# Developing inhibitors of the guanosine triphosphate hydrolysis accelerating activity of Regulator of G protein Signaling-14

**DOI:** 10.1101/2025.06.11.659181

**Authors:** Percy S. Agogo-Mawuli, Isra Sadiya, Tigran M. Abramyan, Dustin E. Bosch, Kyle A. Emmitte, Luis M. Colón-Pérez, Mickey Kosloff, David P. Siderovski

## Abstract

Regulator of G protein Signaling-14 (RGS14), an intracellular inactivator of G protein-coupled receptor (GPCR) signaling, is considered an undruggable protein given its shallow and relatively featureless protein-protein interaction interface combined with a distal allosteric site prone to nonspecific inhibition by thiol-reactive compounds. Here, we identify and validate a tractable chemotype that selectively and non-covalently inhibits RGS14 GTPase-accelerating protein (GAP) activity. Combining structure-guided virtual screening, ligand docking across multiple receptor conformers, and enrichment validation, we progressed from a first-generation active, Z90276197, to over 40 second-generation analogs with improved potency. These inhibitors are predicted to engage the solvent-exposed “canyon” in the RGS14 RGS-box that interacts with the Gα switch I region. Binding pose predictions underscored the importance of non-polar interactions and shape complementarity over polar interactions in engaging this Gα-binding canyon and revealed an “ambidextrous” pattern of R1- and R2-group orientations. GAP inhibition was confirmed in fluorescence-based and gold-standard radioactive GTP hydrolysis assays. Two second-generation analogs, Z55660043 and Z55627844, inhibited RGS14 GAP activity in both assays and without measurable cytotoxicity. Deep learning-based scoring of predicted docking poses further supported observed affinity gains from R3-group additions. One analog demonstrated favorable *in vivo* pharmacokinetics and CNS penetration. Collectively, our findings establish tractable, non-covalent, small molecule inhibition of a G protein regulatory interface and illustrate how machine learning-enhanced docking can guide ligand optimization for shallow protein surfaces. This work opens the door to future development of RGS14 inhibitors as potential therapeutics for central nervous system and metabolic disorders.

## Introduction

The Regulators of G protein Signaling (RGS proteins) were first identified nearly 30 years ago (1–4) as a then “missing regulatory component” (5, 6) of the well-established intracellular pathway of G protein-coupled receptor (GPCR) signaling. Specifically, RGS proteins accelerate the otherwise slow intrinsic guanosine triphosphatase (GTPase) activity of heterotrimeric G protein alpha (Gα) subunits (7–10), leading to GPCR signal termination. As GPCRs historically constitute a significant fraction of the druggable proteome (11, 12), it was thought that uncovering a new intracellular component inhibiting GPCR signaling would quickly unveil novel drug discovery opportunities, especially for central nervous system disorders and diseases (*e.g.*, as reviewed in (13, 14)). For example, soon after discovering RGS14 (15) and identifying shared CNS expression patterns across rodents and humans (*e.g.*, Supplementary Fig. S1a-c), its genetic ablation in mice revealed a key CNS role as a natural suppressor of CA2 neuron synaptic plasticity (16). Loss of mouse RGS14 expression leads to heightened long-term potentiation (LTP) and enhanced hippocampal-based learning and memory (reviewed in (17)). RGS14 knockout mice also have increased levels of peripheral brown adipose tissue (BAT) and exhibit enhancements to metabolism, exercise endurance, and longevity (18, 19), consistent with the idea that RGS14 normally opposes beneficial “beiging” signaling within mouse adipose tissue via the agonist-occupied GPR35 receptor (20). Such preclinical findings of enhanced learning, memory, metabolism, and longevity highlight RGS14 as a potential anti-aging target in both the periphery and CNS. In support of an adipose-centered metabolism function for human RGS14, population genome-wide association studies (GWAS) have established associations between human *RGS14* sequence variations and triglyceride variations (*e.g.*, associations identified in over two million individuals linking *RGS14* variants like SNP rs11746443-G to circulating triglyceride levels [p ∼ 10 ^-15^] (21, 22); Fig. S1d-e); these associations may be direct or instead an indirect result of renal phosphate handling effects that have recently been detailed at the molecular level for RGS14 (23, 24). However, even with all these justifications, only modest progress has been made to date (*e.g.*, (25–27)) in developing RGS protein modulators, possessing reversible binding and physicochemical characteristics common to established therapeutic agents, for preclinical and clinical follow-up for these various indications mentioned above (reviewed in (13, 28)).

The biochemical activity characterizing all 20 “canonical” RGS proteins is Gα-directed GTPase-accelerating protein (GAP) activity (7–10). This GAP activity is technically enzymatic – converting a substrate (Gα·GTP) to a product (Gα·GDP·Pi) without affecting the RGS protein “catalyst”. However, this allosteric activity is in fact mediated by a protein-protein interaction (PPI) between the RGS domain (or “RGS-box”) of the RGS protein and the activated Gα subunit (29, 30). This PPI can be stabilized experimentally when the active site of the Gα subunit is made to mimic the leaving group of inorganic phosphate (Pi), via substitution with the planar ion aluminum tetrafluoride (*i.e*., Gα·GDP·AlF_4_^-^; refs. (29, 31)). Such PPIs are historically more challenging to inhibit with small molecule interactors than GPCRs or enzymes with defined and voluminous active sites, largely because PPI interfaces tend to be large, flat, and topologically featureless, lacking the deep hydrophobic pockets sheltering discrete polar interaction hotspots (*e.g.*, for hydrogen bonding or ionic pairing) that typically accommodate high-affinity small-molecule binding (32, 33). By screening a ∼3,000 compound library using a flow cytometry-based PPI assay that quantified RGS4 binding to the G alpha subunit Gα_o_, an early study identified 4-chloro-N-[methoxy-(4-nitrophenyl)-λ^4^-sulfanylidene]benzenesulfonamide (“CCG-4986”) as an RGS4 GAP activity inhibitor with an *in vitro* IC_50_ of 3∼5 μM; ref. (34)). However, the aryl sulfonamide CCG-4986 was found to be a covalent modifier of cysteines within the RGS-box of RGS4 (35), hindering subsequent drug development based on this early hit. *Reversible* binders, rather than reactive compounds, have better long-term success in a traditional pharmaceutical development pipeline ((36–38); *c.f.* (39, 40)). While an attempt has been made to advance a related compound, CCG-63802, as a reversible, allosteric inhibitor of RGS4 (41), its mechanism of action is also dependent on reactive cysteines within the RGS4 RGS-box but distant from the presumptive Gα-interacting face (42). A different chemotype (1,3-diaryl 1,2,4-(4H)-triazol-5-ones) was originally under exploration for treating urinary incontinence given suspected activity at maxi-K channels (43); this chemotype was subsequently thought to inhibit RGS proteins directed against Gαq based on indirect evidence from model organism genetics (44). However, no direct demonstration of these diaryl-triazolones inhibiting RGS-box GAP function *in vitro* has yet been published in either patent or public literature (44, 45).

Given the advent and democratization of virtual compound screening using protein docking algorithms on high-performance computing platforms (*e.g.*, (46, 47)), we chose to revisit the RGS-box target using newly emerging *in silico* methods and an accumulated volume of RGS-box structural models attained by x-ray crystallography and NMR spectroscopy (*e.g*., (29, 48–51)). We employed the AtomNet® deep-learning system of virtual compound screening on NMR-derived structural models of the RGS-boxes of human RGS14 (PDB id 2JNU; UniProt O43566; ref. (29)) and its closest paralog, human RGS12 (PDB id 2EBZ; UniProt O14924). AtomNet®, a deep convolutional neural network developed for structure-based drug discovery (52–54), was recently applied in one of the largest and most diverse virtual high-throughput screening campaigns to date (55), covering 318 projects across 482 labs in 30 countries and identifying novel drug-like scaffolds across all major therapeutic areas and protein classes. In our application toward RGS-box inhibition, out of 96 AtomNet-predicted interacting compounds obtained from a 2.5-million element “Screening Collection” Enamine chemical library (56), two structurally unrelated compounds were reported (55) as valid micromolar inhibitors *in vitro*. Here, we report on a successful expansion of one of these active compounds into a large class of 1,2,4-triazolo[3,4-b][1,3,4]thiadiazine analogs with verified activities, tested both in the Transcreener® GDP RGScreen™ fluorescence polarization assay of steady-state GTP hydrolysis (57) and in the gold standard of RGS protein GAP activity assay: namely, radioactive GTP single-turnover hydrolysis (7, 58). Given the continuing need for potent and selective RGS protein modulators suitable for preclinical evaluation and eventual therapeutic development, the present study advances this goal by expanding the chemical space targeting RGS14’s orthosteric site -- a shallow Gα-binding canyon.

## Results

### Docking hits validated by *in vitro* GAP inhibition activity

We previously described work with BellBrook Labs creating the Transcreener® GDP RGScreen™ fluorescence polarization assay of steady-state GTP hydrolysis, leveraging a double point-mutant in the Gα enzyme to make GTP hydrolysis, rather than GDP (product) release, the rate-limiting step for multiple rounds of GTP conversion (57, 59). As illustrated in Supplemental Fig. S2A, this *in vitro* assay of RGS-box-accelerated GTPase activity relies on detecting the GDP product, in the presence of excess GTP, using a GDP-selective antibody; displacement of pre-bound fluorescent tracer by GDP binding to this antibody leads to a change in fluorescence anisotropy. Titrating inputs of the double point-mutant (“DM”) Gα_i1_ and RGS-box recombinant proteins (Fig. S2B, C) established a robust, medium-throughput assay for evaluating potential RGS14 and RGS12 inhibitors, with “assay quality” Z’ scores (60) above 0.76 for each target (Fig. S2D). In this assay, BMS-195270, one of the diaryl-triazolones indirectly suspected of having RGS-box inhibitory activity (44) was seen to weakly inhibit both RGS14 and RGS12 RGS-box proteins (estimated IC_50_ for RGS14 of 174 μM [95%CI: 149 to 208 μM], estimated IC_50_ for RGS12 of 75 μM [95%CI: 69 to 81 μM]; Fig. S2E *vs* S2F for the inactive diaryl-triazolone BMS-192364).

We previously reported (55) on the virtual docking of compounds into the conserved Gα-interaction site shared by all RGS proteins (Fig. 1), including RGS14 (61), its closest paralog RGS12 (15), and the archetypal RGS protein RGS4 (7). To focus on human RGS14, the fifth low-energy conformation (PDB id 2JNU “pose” 5) from our NMR-derived structural model of human RGS14 (29) was chosen to provide the original “receptor grid” for docking, based on a survey of all twenty low-energy RGS14 conformations for the largest hydrophobic volume displayed by the conserved Gα-interaction site (55). Two active compounds were identified: the benzimidazole derivative 2-[(5-methoxy-1H-1,3-benzodiazol-2-yl)sulfanyl]-1-(2,4,6-trifluorophenyl)propan-1-one (a.k.a. Enamine Z1098346112; Fig. S3A, published in refs. (55, 62)) and the triazolothiadiazine derivative 2-{3-[(4-chlorophenoxy)methyl]-6-phenyl-5H-[1,2,4]triazolo[3,4-b][1,3,4]thiadiazin-7-yl}acetic acid (a.k.a. Enamine Z90276197). The benzimidazole Z1098346112 failed to spawn a robust analog series of similar or enhanced potency. Of 83 analogs tested in the Transcreener® GDP RGScreen™ assay (Supplementary Table S1), only a single related compound from Enamine’s multi-billion element “REadily AccessibLe” (REAL) library (63) showed comparable potency against RGS14 and RGS12 – namely, Enamine Z1098346193, an N-bridged benzimidazole isostere featuring a triazolo[1,5-a]pyrimidine ring system (Fig. S3B). As actives Z1098346112 and Z1098346193 share a common (2,4,6-trifluorophenyl)propan-1-one group (Fig. S3), several similar-sized, multi-ringed, non-benzimidazole compounds containing a 2,4,6-trifluorobenzoyl group were also tested, but none exhibited potent RGS14 GAP activity inhibition (Supplementary Fig. S4).

**Figure 1.**
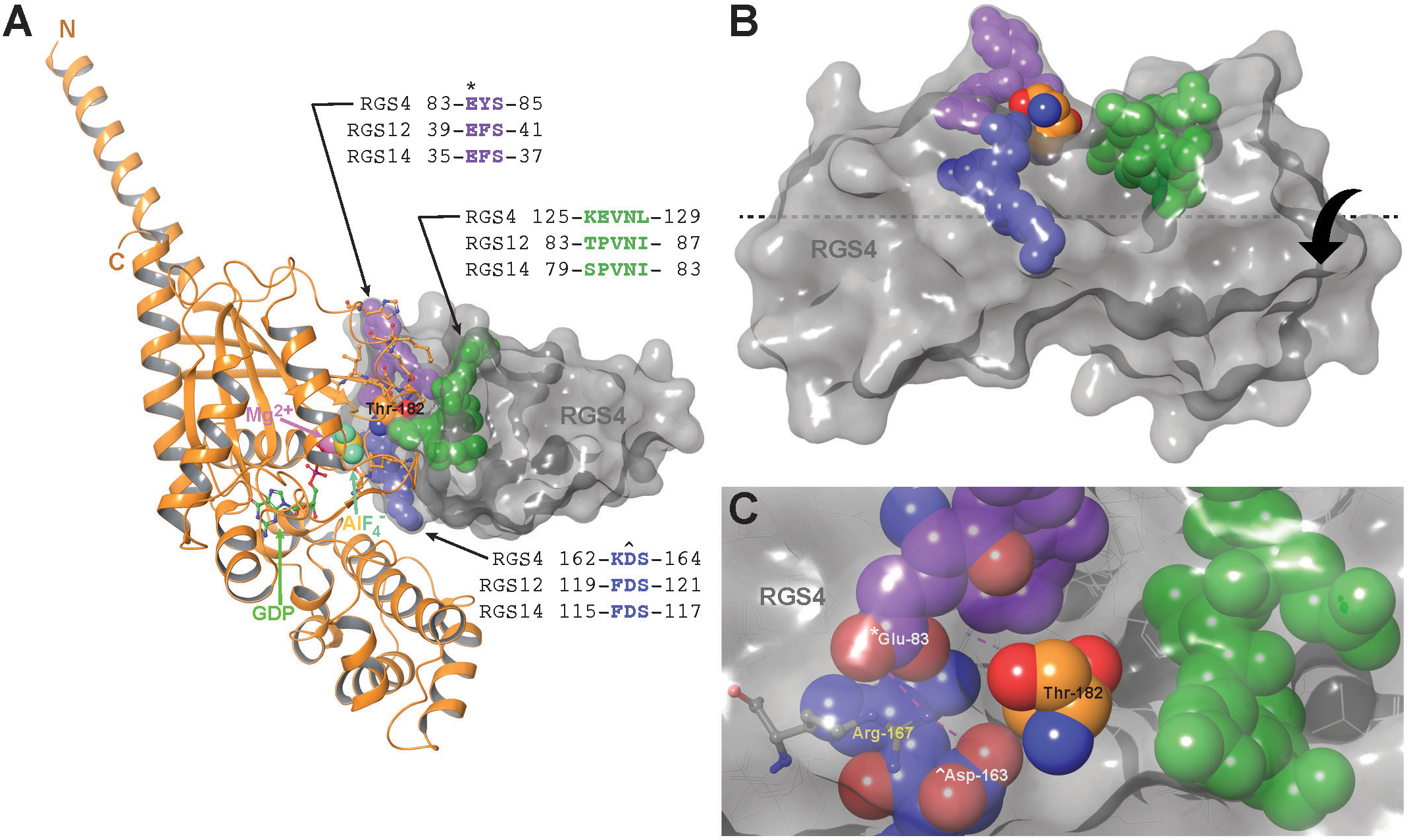
Illustration of the target site used for *in silico* docking of virtual compound libraries into the Gα-interaction face of RGS14 and RGS12 (55), based on the archetypal structural model of the RGS4 / Gα_i1_·GDP·AlF ^-^ complex (PDB id 1AGR; ref. (48)). **(A)** Representation of the RGS4 / Gα_i1_·GDP·AlF4 ^-^ structural model, with the Gα subunit in orange ribbons and side-chains approaching RGS4 in Corey–Pauling–Koltun (CPK)-colored pipes; the Gα_i1_ switch I region’s threonine-182 side-chain (numbered based on PDB id 1AGR) is illustrated in CPK-colored space-filling spheres. The aluminum tetrafluoride planar anion (AlF4 ^-^) and magnesium counterion (Mg ^+^) are also illustrated in space-filling spheres, as indicated. The RGS4 structural model is represented by a gray surface, with key amino acids near the Gα_i1_ threonine-182 side-chain highlighted in purple, green, and blue space-filling spheres; sequences of these three key amino-acid spans are also highlighted in a multiple sequence alignment of corresponding RGS4, RGS12, and RGS14 polypeptides (numbered based on PDB ids 1AGR, 2EBZ, and 2JNU, respectively). **(B)** Side view of the shallow canyon within RGS4 into which the Gα_i1_ switch I region’s threonine-182 side-chain is inserted. **(C)** Top view of the shallow canyon within RGS4 into which the Gα_i1_ switch I region’s threonine-182 side-chain is inserted, highlighting the conserved glutamate-83 (*) and aspartate-163 (^) residues within RGS4 that are each salt-bridged by a terminal nitrogen of RGS4’s arginine-167 (*i.e.*, the residue in CPK-colored ‘pipe’ layout; salt bridges illustrated with purple dashed lines).

### Active compounds inhibit cysteine-less RGS-boxes of RGS14 and RGS12

Given the earlier experience with screening hit CCG-4986 being a covalent modifier of RGS4 cysteine residues (35), we sought to exclude protein-thiol reactivity as a potential mechanism of action of the active compounds derived from the AtomNet® virtual screening effort (55) and SAR campaign follow-ups. To address this possibility and screen out thiol-reactive hits, cysteine-lacking versions of recombinant proteins spanning the RGS-boxes of RGS14 and RGS12 were purified from *E. coli* cultures. These “cysteine-less” proteins were then verified as possessing *in vitro* GAP activity (Figure S5A,B) on the rate-modified mutant (“DM”) of Gα_i1_·GTP (57, 59). Actives from both the benzimidazole and triazolothiadiazine classes were observed to inhibit GDP production by cysteine-less RGS14 and RGS12 RGS-box proteins in these assays (*e.g*., benzimidazole Z1098346112 [Figure S5C,D; estimated IC_50_’s 26∼64 μM] and triazolothiadiazine Z55660043 [Figure S5E,F; estimated IC_50_’s 10∼19 μM]).

### Expansion of the triazolothiadiazine chemotype and analysis of structure-activity relationships

In contrast to our inability to expand within the benzimidazole chemotype, 48 additional active compounds were found within a set of 83 analogs related to the 1,2,4-triazolo[3,4-b][1,3,4]thiadiazine early hit Z90276197 (*e.g.*, eight representatives shown in Figure 2; the entire set of actives is shown across Supplementary Figures S6-S8 and inactives shown in Supplementary Table S2). The early hit Z90276197 (pIC_50_ ^RGS14^ of 4.09 [*i.e.*, IC_50_ of 81 μM]) contains an acetic acid substituent and spawned 20 additional acidic analogs. Some of these acidic analogs trended toward selectivity for RGS14 inhibition over that of RGS12 (*e.g.*, Z90275816, Z90278051, and Z90274338; pIC_50_ ratio estimates of 1.07 to 1.14 as shown in Figure 2). However, these acidic analogs did not progress into substantially improved potency against RGS14 GAP activity (Supplementary Figure S6: n = 21, mean pIC_50_ ^RGS14^ of 4.09, min = 3.52, max = 4.69 [*i.e.*, max IC_50_ of 20 μM]). The acetic acid group was subsequently found dispensable to inhibitory activity (Figure 3A,B *vs* C,D).

**Figure 2.**
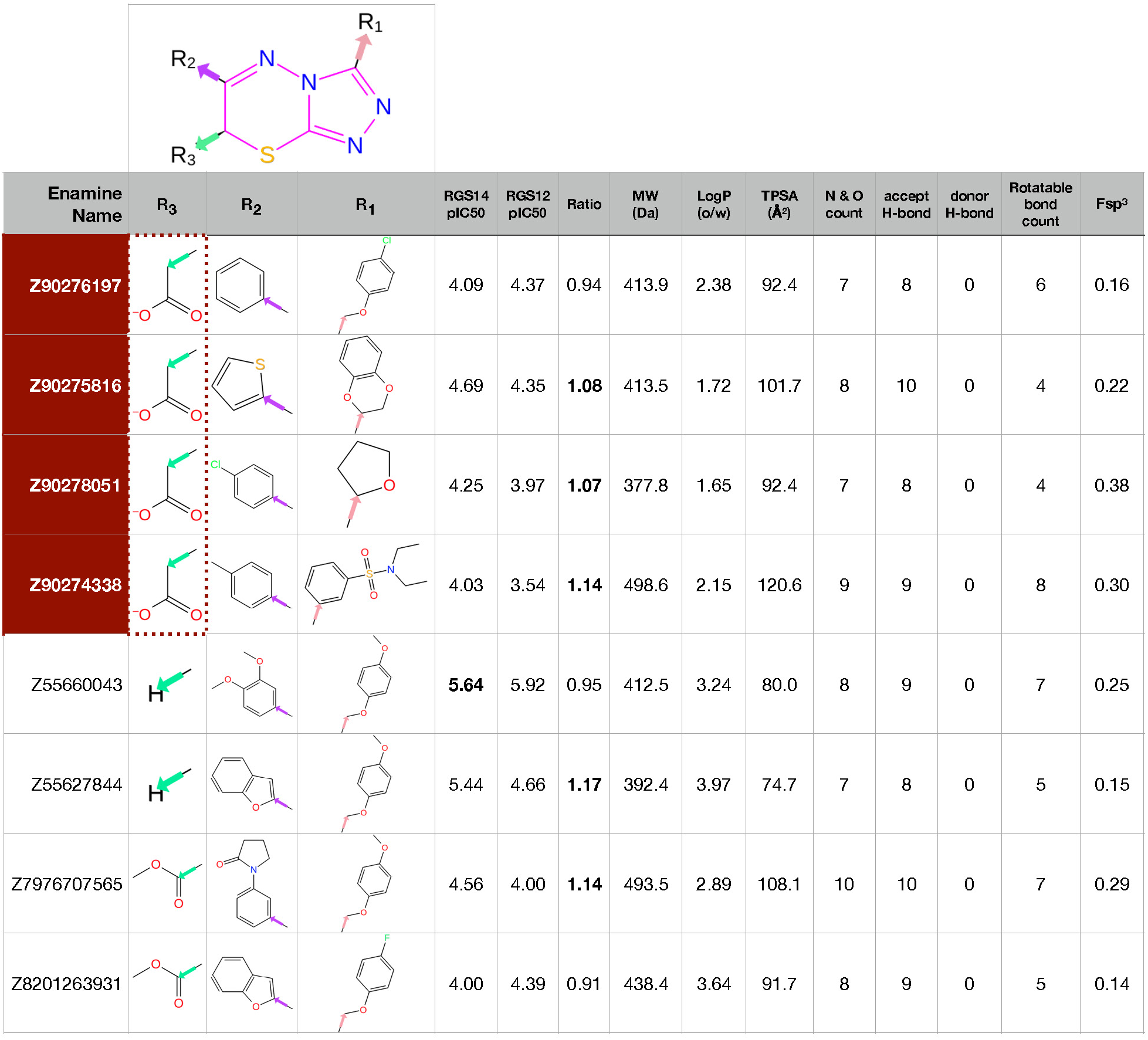
Activities and computed properties of eight representative compounds of the 1,2,4-triazolo[3,4-b][1,3,4]thiadiazine series of RGS-box inhibitors, with constituent R-groups listed below the shared triazolothiadiazine core, including the original screening hit Z90276197 listed first (a member of the unprotonated acetic acids in red backgrounds) and the highest potency analog Z55660043 (highlighted in **bold**: IC_50_ of 2.3 μM against wild-type RGS14 RGS-box). ***RGS14 pIC_50_***, absolute value of log_10_ of the compound concentration providing 50% inhibition of wild-type RGS14 RGS-box GAP activity, as measured in the Transcreener® GDP-detection RGScreen™ assay using the Gα_i1_(DM) subunit; ***RGS12 pIC_50_***, absolute value of log_10_ of the compound concentration providing 50% inhibition of wild-type RGS12 RGS-box GAP activity. ***Ratio*** column represents the calculated ratio of the measured pIC_50_ values for RGS14 inhibition over RGS12 inhibition (values >1.0, reflecting RGS14 selectivity, are highlighted in **bold**). Enumerated and estimated physicochemical properties for each compound were obtained using RDKit Chem Descriptors (95): ***MW***, molecular weight (units of Daltons); ***LogP***, log_10_ of predicted octanol/water (o/w) partition coefficient; ***TPSA***, total polar surface area (units of angstroms-squared); ***N & O count***, enumeration of nitrogens and oxygens within the compound; ***accept H-bond (a.k.a. HBA)***, enumeration of nitrogens, oxygens, and sulfurs within compound (sum used by RDKit as estimation of possible H-bond acceptors); ***donor H-bond** (a.k.a. HBD)*, enumeration of possible H-bond donor atoms; ***rotatable bond count***, the number of bonds in the compound that can be rotated without breaking the molecular structure or altering connectivity; ***Fsp^3^***, fraction of sp^3^ hybridized carbon atoms relative to the total number of carbon atoms in the compound.

**Figure 3.**
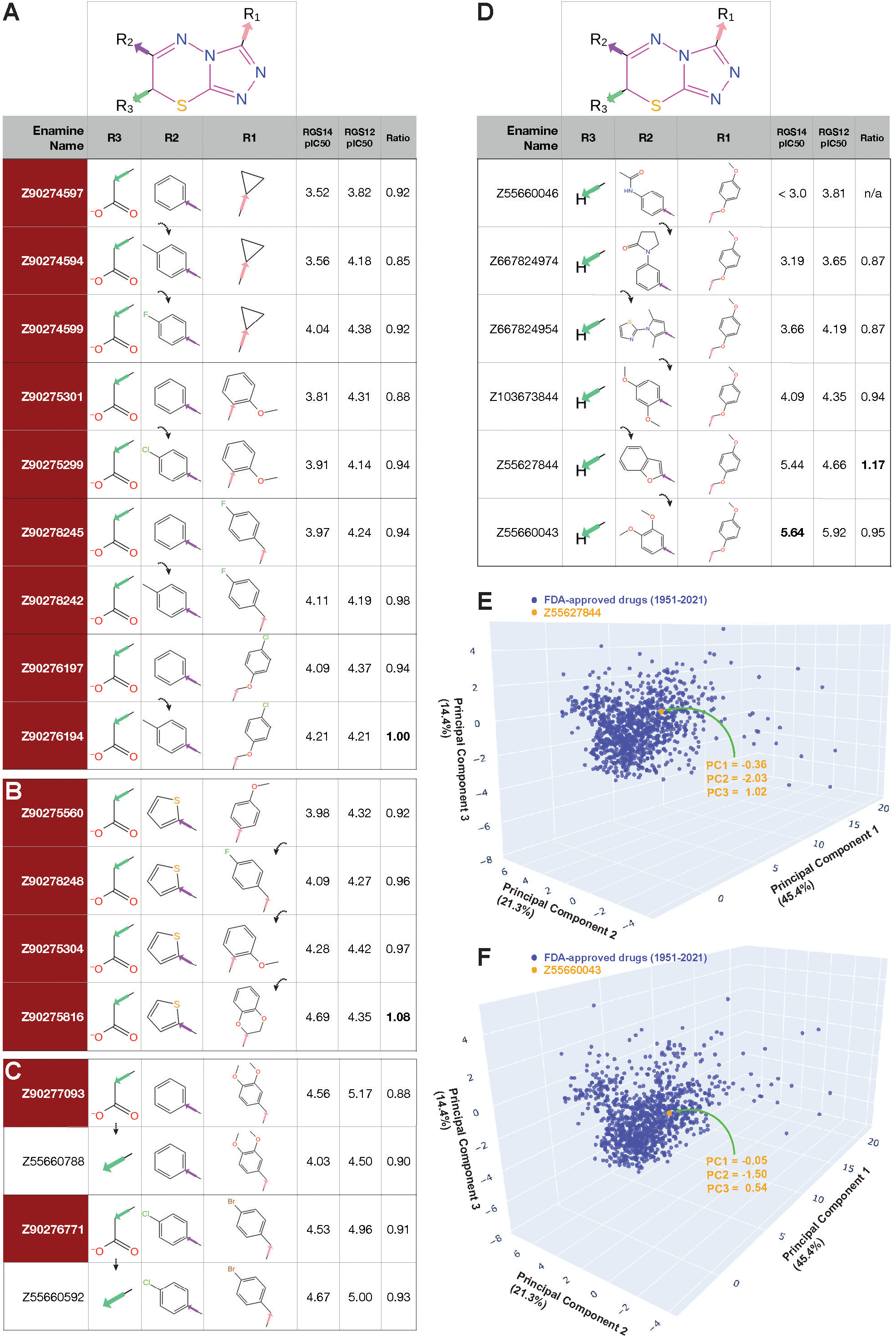
Selected structure-activity relationships observed within the 1,2,4-triazolo[3,4-b][1,3,4]thiadiazine chemotype. **(A)** As highlighted with dotted arrows, *para*-methyl or -halogen substitutions to an R2-group benzene ring within triazolothiadiazine-core acetic acid derivatives (shaded in red background) increase observed potency against RGS14 GAP activity. **(B)** The R1 position of RGS14 inhibitors favors substituted benzene rings, in isolation or as a benzofuran attached by methoxy bridging (*e.g.*, Z90275816). **(C)** An acid moiety at the R3 position is not required for activity; for example, replacement of the acetic acid R3-group within Z90276771 by a methyl group (within Z55660592) increases observed inhibitor potency against both RGS14 and RGS12 GAP activity. **(D)** An R3 group is not required for activity. **(E, F)** Principal component analysis of 1112 FDA-approved drugs from 1951 to 2021 (**blue** dots) using input parameters of MW, LogP, HBA, HBD, and Fsp^3^ (each as defined in Figure 2 legend). Later-stage triazolothiadiazine analogs, like Z55627844 (panel E) and Z55660043 (panel F), are observed (**orange** dots) to vary only minimally from the center of the FDA-approved drugs’ distribution across the three principal component axes.

Varying the other two R-groups of the triazolothiadiazine core revealed that *para*-methyl or *para*-halogen group substitutions to a benzene ring at the R2 position modestly increase potency against RGS14 GAP activity (Fig. 3A). The R1 position was found to favor substituted benzene rings, either in isolation or as a benzofuran moiety attached to the 1,2,4-triazolo[3,4-b][1,3,4]thiadiazine core by a methoxy bridge (Fig. 3B). Preserving a *para*-methoxyphenyl group with a methoxy-methylene linkage at the R1 position and changing the R2 position with various ring structures in the absence of an R3 group resulted in the highest potency observed for this chemotype against both RGS14 and RGS12 GAP activity (Z55660043: RGS14 pIC_50_ of 5.64, RGS12 pIC_50_ of 5.92; Fig. 3D).

### Computed, predicted, and observed drug-like features of the triazolothiadiazine class of inhibitors

Physicochemical properties known to be generally relevant to drug-likeness were predicted across all 49 triazolothiadiazine actives. Z55660043, otherwise known as 6-(3,4-dimethoxyphenyl)-3-[(4-methoxyphenoxy)methyl]-5H-[1,2,4]triazolo[3,4-b][1,3,4]thiadiazine, and similar methoxyphenoxymethyl-substituted triazolothiadiazine actives were predicted to be close in character to the majority of FDA-approved drugs (*e.g.*, Figure 3E, F). Molecular weights amongst all these triazolothiadiazine actives (Figure 4) range from 264.4 to 498.6 Daltons (mean ± SE: 394.0 ± 6.8 Daltons), which is within the size range (< 500 Da) that serves as a first pillar of Lipinski’s Rule of Five originally formulated to avoid problem compounds with poor absorption or permeation in downstream development (64). Atoms with the potential to accept (HBA) or donate (HBD) hydrogen bonds during protein-ligand interactions were also found within the ranges canonized by Lipinski’s Rule of Five (HBA: range 5 - 10, mean ± SE: 7.6 ± 1.2, rule ≤ 10; HBD: range 0 - 1, mean ± SE: 0.1 ± 0.3, rule ≤ 5). Values for the calculated partition coefficient across an octanol/water biphasic system (a.k.a. LogP(o/w)) were also found to be compatible with Lipinski’s Rule of Five for all 49 triazolothiadiazine actives (range 1.02 - 5.03, mean ± SE: 3.0 ± 1.1, rule < 5). Of these four parameters within Lipinski’s Rule of Five, only one was weakly but significantly correlated with RGS14 selectivity in simple linear regression analyses: namely, hydrogen bond acceptors (Figure 4). In addition, calculated total polar surface area was also correlated with RGS14 selectivity by simple linear regression (Figure 4); however, other compound properties were not found to be significantly correlated: specifically, LogP(o/w), rotatable bond count, and fraction of *sp^3^* hybridized carbon atoms relative to total carbon atom count (Fsp^3^), the latter an index of three-dimensional complexity often correlating with desirable pharmacokinetic properties of improved solubility, permeability, and metabolic stability (65).

**Figure 4.**
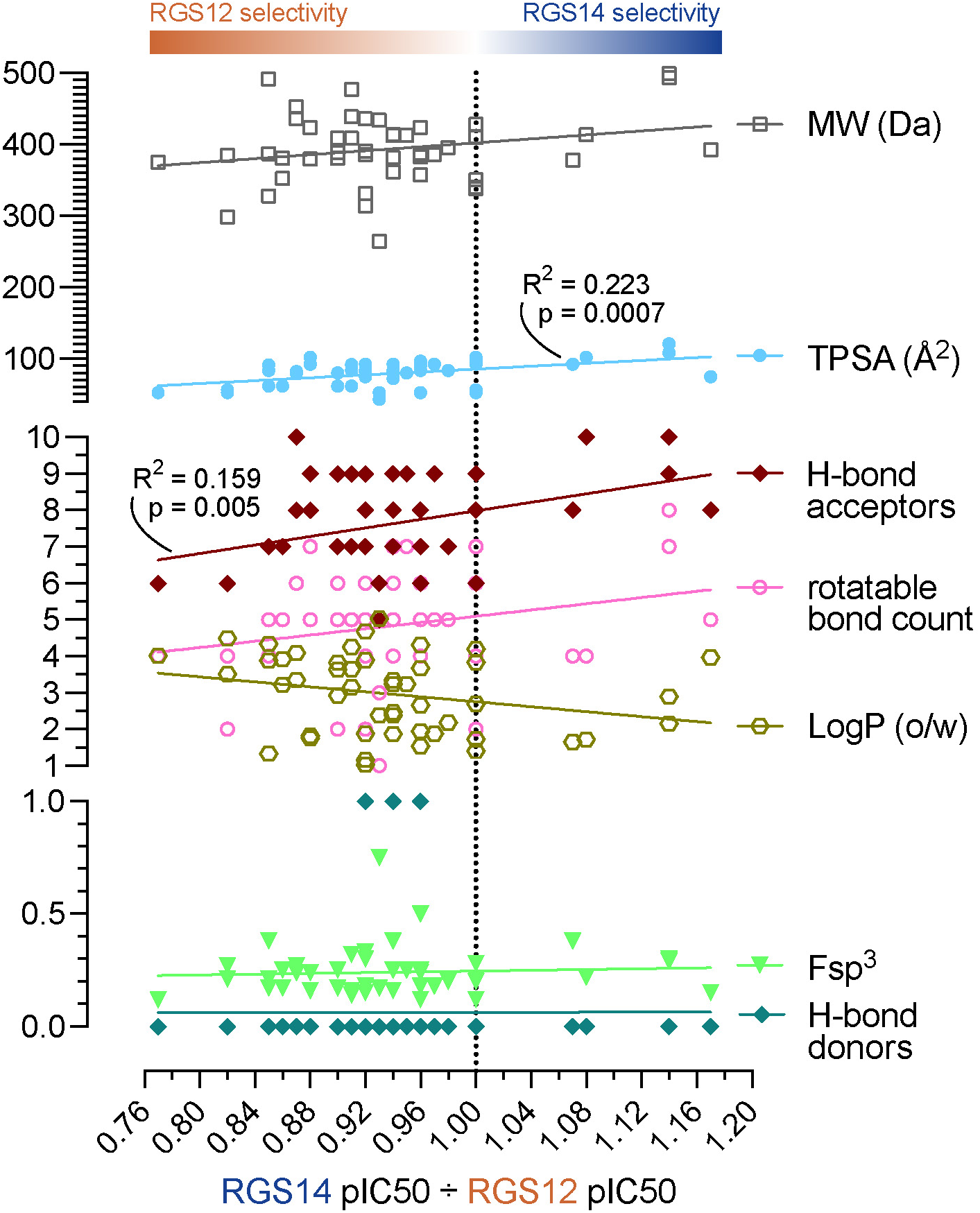
Computed physicochemical properties of the triazolothiadiazine series of RGS14 inhibitors plotted against estimated RGS14 selectivity (*i.e.*, pIC_50_ observed for RGS14 inhibition over pIC_50_ observed for RGS12 inhibition). ***MW***, molecular weight (Daltons); *TPSA*, total polar surface area (angstroms-squared); *H-bond acceptors (a.k.a. HBA)*, enumeration of nitrogens, oxygens, and sulfurs; *rotatable bond count*, number of bonds that can be rotated without breaking the structure; ***LogP***, log_10_ of predicted octanol/water (o/w) partition coefficient; ***Fsp^3^***, fraction of sp^3^ hybridized carbon atoms; ***H-bond donors** (a.k.a. HBD)*, enumeration of possible H-bond donor atoms. Statistics for significant linear associations are indicated (R-squared “goodness-of-fit” and associated p-values), as derived by simple linear regression using GraphPad Prism 10.

According to a recently described deep-learning model for pharmacokinetic and toxicology predictions (66), all triazolothiadiazine actives are predicted to be orally bioavailable, intestinally absorbed, and not affected by P-glycoprotein efflux; furthermore, the majority of them are also anticipated to lack mutagenic/carcinogenic and cardiac toxicities and to penetrate the blood-brain barrier (Supplementary Table S3). To test the latter prediction, two representative triazolothiadiazine derivatives, Z55628018 and Z56944652 (Figure 5A,B), were separately evaluated in cytotoxicity assays and in single-dose pharmacokinetic studies. Neither triazolothiadiazine derivative was observed to induce significant cell death within *in vitro* cultures of the immortalized cell line THP-1 (Figure 5C). Each triazolothiadiazine derivative was then separately administered intraperitoneally to male Swiss-Webster mice, and plasma and brain concentrations were assessed at multiple time points up to 24 hours post-dose using validated LC-MS/MS methods. Systemic exposure and brain penetration were quantified based on compound-specific LCMS calibration curves from plasma extracts and brain tissue homogenates. Z56944652 at a single dose of 10 mg/kg was CNS penetrant, reaching peak brain levels (∼20 ng/g tissue) at 1 hour post-dose. The brain-to-plasma concentration ratio (Kp) approximated 1.0 throughout the measured time course (Figure 5D), indicating effective CNS exposure. In contrast, Z55628018 was only ever detected in one of three mice in plasma at the 1-hour time point (0.17 µg/mL), even after adjusting the PK protocol to increase the dose to 30 mg/kg and pre-dosing with 50 mg/kg 1-aminobenzotriazole (to reduce possible cytochrome P450 metabolism; ref. (67)) two hours prior to Z55628018 administration. No measurable concentrations of Z55628018 were observed in brain tissue at any time point, suggesting poor systemic bioavailability and CNS distribution of this particular triazolothiadiazine derivative under the tested conditions. Z56944652, irrespective of its more favorable CNS penetration character, was subsequently abandoned owing to evidence of interference with Transcreener® GDP detection reagents and potential non-specific inhibition of the intrinsic GTPase activity of an infectious amoeba-sourced Gα subunit (Figure 5E).

**Figure 5.**
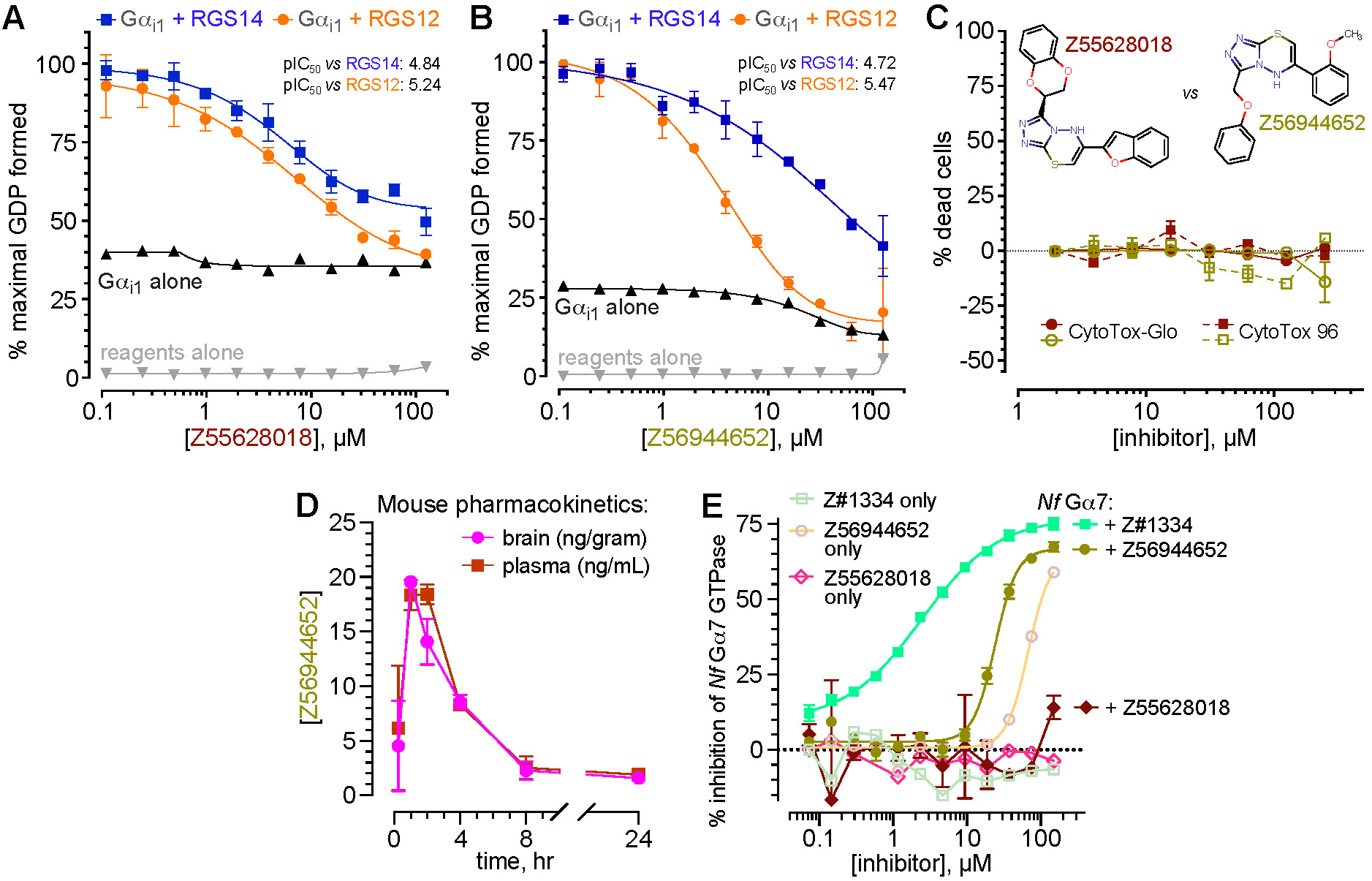
Assessments of cellular toxicity, pharmacokinetics, and non-specific actions of triazolothiadiazine derivatives Z55628018 and Z56944652. (**A,B**) Concentration-dependent inhibition of RGS14 (blue) and RGS12 (orange) RGS-box GAP activity by 6-(1-benzofuran-2-yl)-3-(2,3-dihydro-1,4-benzodioxin-2-yl)-5H-[1,2,4]triazolo[3,4-b][1,3,4]thiadiazine (Z55628018; panel A) and 6-(2-methoxyphenyl)-3-(phenoxymethyl)-5H-[1,2,4]triazolo[3,4-b][1,3,4]thiadiazine (Z56944652; panel B), as measured by the Transcreener® GDP-detection RGScreen™ assay. (**C**) Independent tests of the cytotoxicity of Z55628018 and Z56944652 performed using the immortalized THP-1 leukemic cell line and two different assays for cell viability: a luminescence-based CytoTox-Glo assay for intracellular ATP content, and a colorimetric-based CytoTox 96 assay for lactate dehydrogenase release. (**D**) Evidence of CNS penetration of Z56944652 provided by intraperitoneal injection (10 mg/kg) to three Swiss Webster mice: at this initial dose, Z56944652 achieved a peak CNS level at 1 hr of 19.5 ± 0.3 ng/gram brain tissue (mean±SD). (**E**) In a separate assay of steady-state GDP production by the *Naegleria fowleri* Gα7 GTPase (83), using Transcreener® GDP detection reagents in a previously published protocol (84), Z56944652 was observed to “inhibit” *Nf* Gα7 GTPase activity with an apparent IC_50_ of ∼25 μM ; much of this observation is confounded by compound interference of the fluorescence anisotropy-based readout of GDP production (*i.e.*, parallel reaction conducted without input GTPase: “Z56944652 only”). Also shown are positive results from a previously reported, active small-molecule inhibitor of *Nf* Gα7, coded Z#1334, which does not possess confounding compound interference of the GDP production readout (84).

To further evaluate the functional activity of these triazolothiadiazine derivatives, and especially to eliminate concerns of misascribing GAP-inhibitory activity to potential assay interference, we tested two representative compounds -- Z55660043 and Z55627844 -- in the gold-standard single-turnover radioactive GTP hydrolysis assay (7, 58) (Figure 6). In a concentration-dependent fashion, each of the two actives was observed *in vitro* to inhibit RGS14 RGS-box-mediated acceleration of [γ-^33^P]GTP hydrolysis catalyzed by wild-type Gα_i1_ protein (Figure 6C,D), consistent with genuine inhibition of RGS-box GAP activity. Importantly, both inhibitors demonstrated no overt cytotoxicity in independent cell-based viability assays using the THP-1 cell line, as assessed by both ATP content (CytoTox-Glo) and LDH release (CytoTox 96) measurements (Figure 6, panels E and F). Together, these results validate the RGS GAP-inhibitory activity of Z55660043 and Z55627844 in an orthogonal assay free from potential reagent interference, while also supporting their preliminary tolerability.

**Figure 6.**
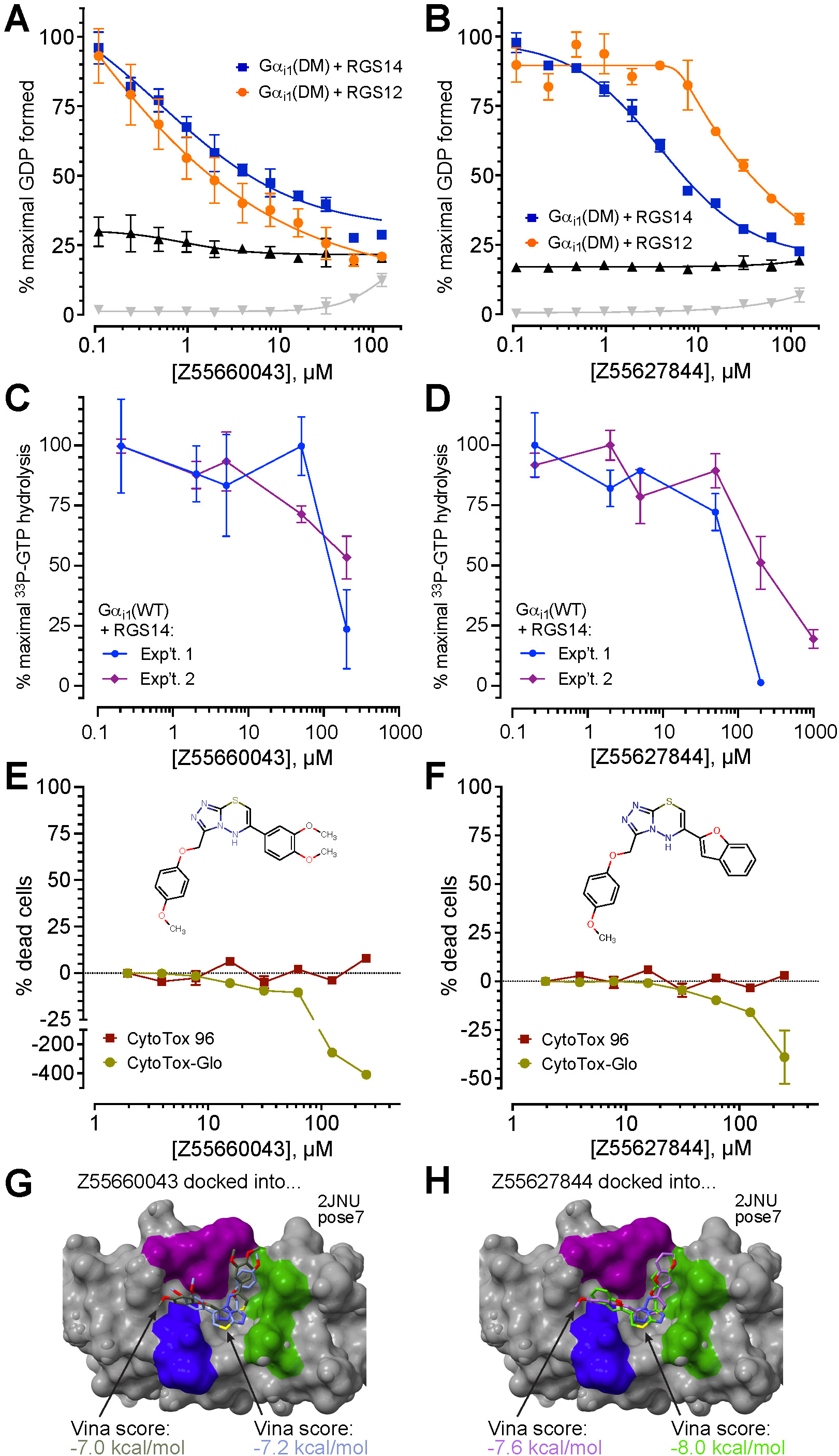
Testing two second-generation inhibitors of the triazolothiadiazine chemotype – Z55660043 and Z55627844 – with the Transcreener® GDP assay, the gold-standard [γ -^33^P]GTP hydrolysis assay, and cytotoxicity assays. **(A,B)** Transcreener® GDP reactions containing indicated compounds, performed as in Fig. S2. Note modest effects (if any) of the compounds on the intrinsic GTPase rate of Gα_i1_(DM) protein alone (interference tracking in **black**) or on the Transcreener® GDP detection reagents alone (interference tracking in **gray**). **(C,D)** Single-turnover [γ-^33^P]GTP hydrolysis with increasing concentrations of the same compounds, measured using human Gα_i1_(WT) protein. Data normalized to maximal observed [^33^P]-Pi production within each independent experiment and plotted as means ± SEM from each independent experiment. **(E,F)** Independent tests of cytotoxicity by Z55660043 and Z55627844 performed using the immortalized THP-1 leukemic cell line and two different assays for cell viability: a luminescence-based CytoTox-Glo assay for intracellular ATP content and a colorimetric-based CytoTox 96 assay for lactate dehydrogenase release. Compound structures presented to highlight observable luciferase-based assay interference presumably caused by presence of the shared *p*-dimethoxybenzyl R-group substituent within Z55660043 and Z55627844, as seen in the CytoTox-Glo assay (*i.e.*, “*negative* % dead cells” at highest concentrations; interference not seen within the alternative, colorimetric-based CytoTox 96 assay). **(E,F)** Representative, top-scoring docking poses for Z55660043 and Z55627844 within the Gα-interaction site of RGS14 as represented by PDB id 2JNU pose 7 and derived from the GNINA implementation of the Vina docking algorithm (69). Color scheme of the RGS14 structural model (**gray** surface), with key Gα-interacting amino acids highlighted in **purple**, **green**, and **blue**, is consistent with the model originally presented in Fig. 1; note that the color of the Vina score matches the color of the inhibitor’s carbons, and that the two inhibitor poses within panels G and H represent flipped orientations of the R1- and R2-groups’ relative positions *versus* the tri-color “signposts” of the central RGS-box canyon.

### Active triazolothiadiazines are predicted to bind RGS14 RGS-box in a conserved and reciprocal fashion

The initial triazolothiadiazine derivative Z90276197 was originally identified via high-throughput virtual screening (55) of an Enamine-sourced compound library against a single, pre-selected docking site within the RGS14 RGS-box. This site was defined by three-dimensional structural coordinates of pose 5 from PDB id 2JNU corresponding to the site homologous to where the switch I Thr-183 side chain of Gα_i1_·GDP·AlF_4_^-^ inserts into RGS4 (Figure 2). To predict the binding mode(s) for Z90276197 and the 48 related triazolothiadiazine derivatives with confirmed RGS-box inhibitor activity, each compound was docked *in silico* against 40 available low-energy conformers of the RGS14 and RGS12 RGS-boxes (20 low-energy conformers each from PDB ids 2JNU and 2EBZ, respectively), as derived by NMR (29). To ensure unbiased binding site predictions, we used GNINA (68), an AutoDock Vina software fork, for this “*49 ligands X 40 receptors*” docking experiment. GNINA was selected because, unlike the original Vina algorithmic implementation (69), it uniquely allows for the entire surface of the protein “receptor” to present possible docking sites for the candidate ligand being docked, as schematized by the dotted box in Figure 7A. The presumptive Gα-interacting site of the seventh low-energy structural pose of the RGS14 RGS-box (“pose 7” of PDB id 2JNU) consistently yielded the highest predicted Vina affinity score (in kcal/mol) for each of the 49 active triazolothiadiazines (Figure 7B).

**Figure 7.**
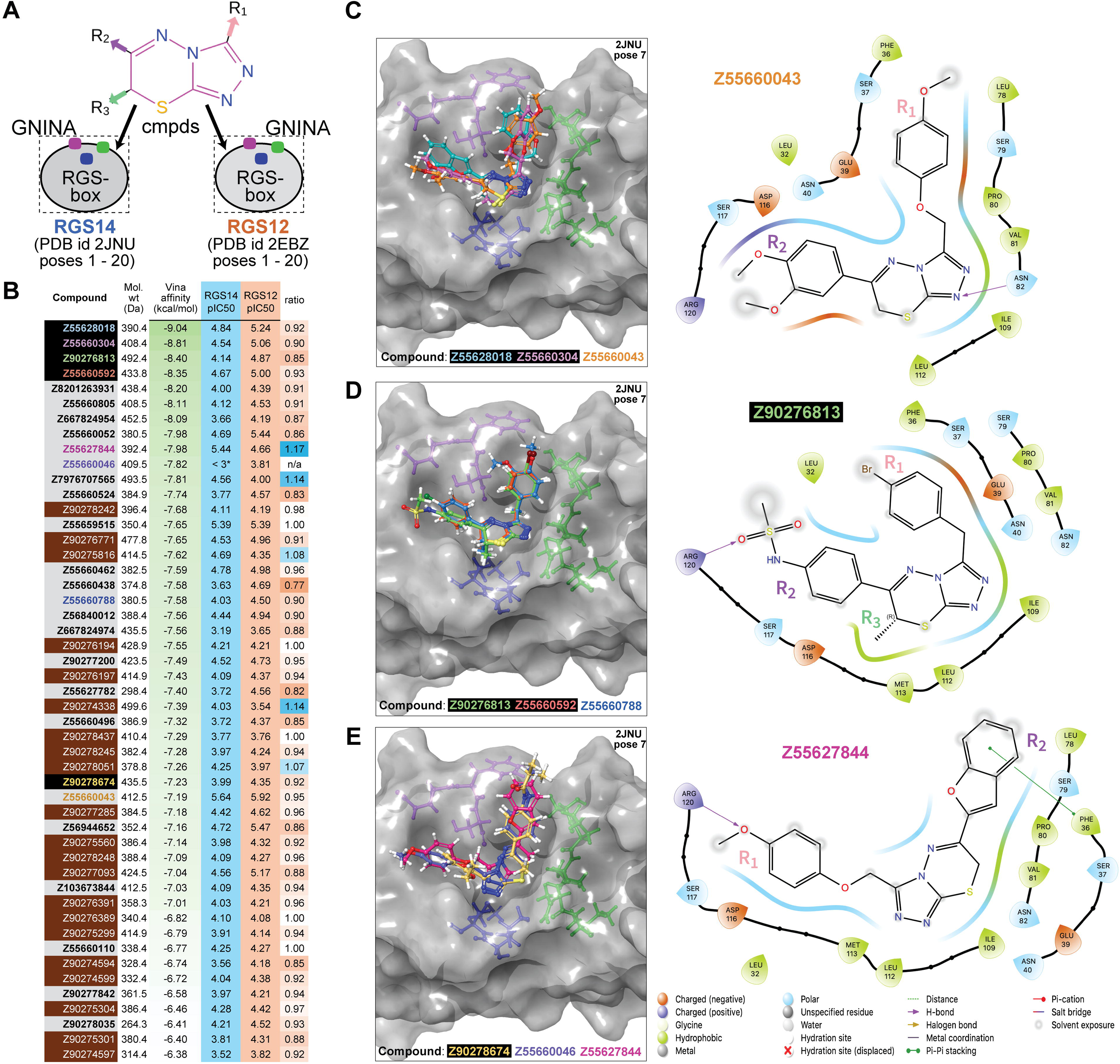
Unbiased assessments of predicted RGS-box binding sites and ligand orientations for the 49 active triazolothiadiazine derivatives. (**A**) Schematic representation of a large-scale re-docking experiment in which all 49 active triazolothiadiazine compounds were docked across 40 structural conformers of RGS-box domains derived from RGS14 (PDB id 2JNU) and its closest paralog RGS12 (PDB id 2EBZ), using GNINA (v1.0) in unbiased docking mode. Color-coded orientation markers (purple, green, and blue) indicate key structural landmarks around the presumptive Gα-binding interface, analogous to the switch-region pocket originally defined in the RGS4/Gα_i1_ complex (see Figure 1). Ligands were allowed to explore the full receptor surface using GNINA’s `--autobox_ligand receptor.pdbqt` mode, without constraints on docking location. (**B**) Summary of Vina-predicted binding affinities and experimental GAP inhibition data for each compound, organized by decreasing Vina score against pose 7 of RGS14 (PDB id 2JNU). This protein conformation was identified as the most discriminating receptor in enrichment analysis (see Supplementary Figure S9) and was therefore used for all downstream pose evaluations. The Compound column is color-coded to highlight structural and biochemical features: compounds with carboxylic-acid R3-groups are boxed in red, while compound identifiers shown in colored text correspond to those depicted structurally in panels C–E. Green gradient shading in the Vina affinity column reflects increasing predicted binding strength (more negative kcal/mol values), whereas background color in the pIC_50_ columns distinguishes between RGS14 (blue) and RGS12 (orange) GAP activity inhibition. Selectivity ratios greater than 1.0 are shaded in blue to indicate RGS14 preference, while ratios less than 1.0 are shaded in orange to reflect greater inhibition of RGS12 GAP activity. (**C–E**) Representative top-scoring docking poses for select compounds within the Gα-interaction site of RGS14 as represented by PDB id 2JNU pose 7. Ligands exhibit diverse “ambidextrous” orientations, with R1 and R2 groups projecting into opposite walls of the shallow canyon (*e.g*., compare R1 and R2 group orientations of Z55660043 (panel C) *versus* Z55627844 (panel E) relative to amino acids of the RGS14 Gα-interaction site in the 2D projections). Ligand core positioning and surface complementarity are consistent with predominantly steric and van der Waals contributions to binding energy, in line with the weak-micromolar IC_50_ values observed experimentally and the feature-poor nature of the protein–protein interface being targeted.

To independently validate the preferential performance of pose 7 of PDB id 2JNU as the top-ranking target (“receptor”) conformation for these active triazolothiadiazines, an additional docking experiment was performed using the Vina docking algorithm, with the presumptive Gα-interacting site now explicitly defined as the docking grid for each of the 40 RGS-box conformers and with a collection of 4160 decoy compounds added to the 49 active triazolothiadiazines as docking ligands to enable enrichment calculations. Across all 40 docking grids, pose 7 of PDB id 2JNU provided the best enrichment performance as measured by both the receiver operating characteristic area under the curve (ROC AUC) and the robust initial enhancement (RIE) metrics (*e.g.*, Supplementary Figure S9). Specifically, the 2JNU pose 7 conformation yielded an ROC AUC of 0.63 and an RIE of 2.58 -- higher than any other RGS14 or RGS12 RGS-box conformer tested in the validation experiment. Although these enrichment scores fall short of the near-ideal ROC values (∼1.0) observed in benchmark systems with highly optimized and discriminatory receptor grids, they nonetheless indicate that 2JNU pose 7 possesses the best available discriminatory power for distinguishing known active triazolothiadiazines from decoys at the presumptive Gα-binding site, particularly in the early enrichment regime emphasized by the RIE metric.

Predicted binding poses of the active triazolothiadiazine derivatives within pose 7 of the RGS14 RGS-box revealed a consistent structural motif -- all ligands were docked within a shallow, solvent-exposed canyon situated at the presumptive insertion point of Gα_i1_ Thr-183 previously described in the homologous RGS4 complex (Figure 1). This binding groove accommodates the planar triazolothiadiazine core in a central anchoring position, with the R1 and R2 substituents projecting outward into adjacent subpockets. Notably, the predicted protein–ligand interactions were dominated by non-polar van der Waals contacts and steric complementarity, with only sporadic hydrogen bonds or π–π stacking interactions observed such as the R2 sulfoxide group of compound Z90276813 predicted to hydrogen bond with Arg120 (Figure 7D), or the R1 ether oxygen and R2 aryl group of compound Z55627844 engaging in polar and π-stacking interactions with Arg120 and Phe36, respectively (Figure 7E).

The R1 and R2 groups are predicted to exhibit recurring inversion of their spatial orientations across the ligand series, such that either substituent could occupy the region proximal to Phe36 or Arg120, depending on the compound. For example, compounds Z55660043 and Z90276813 adopt poses with R1 near Phe36 and R2 near Arg120 (Figures 7C and 7D), while Z55627844 displays the reverse configuration in its highest-scoring pose (Figure 7E). This apparent bidirectional accommodation of ligand termini around a fixed triazolothiadiazine core suggests a flexible, dual-site recognition mode. We refer to this pattern as “ambidextrous binding” -- a symmetric binding behavior in which both R1 and R2 moieties can alternately engage with either subpocket of the shallow canyon, while preserving central scaffold orientation. This mode of binding may enhance the chemical diversity tolerated at the termini of the bound small molecule and reflects the canyon’s permissive topography. Modestly sized R3 groups, such as the methyl moiety of compound Z90276813 (Figure 7D), are accommodated by shallow burial within the canyon-shaped groove. Extension of this position with a methyl ester substituent was predicted to enhance ligand affinity via deeper burial and improved shape complementarity, sometimes accompanied by an inversion of ligand orientation to better accommodate the enlarged R3 group. These computational predictions were borne out experimentally: compounds bearing a methyl ester group at R3 demonstrated improved inhibitory potency in RGS-box GAP assays, but also showed increased cytotoxicity in cell-based viability assays (*e.g.*, Supplementary Figure S10).

## Discussion

Our study suggests that small molecules can selectively and non-covalently inhibit the GAP activity of RGS14 -- a historically undruggable target -- via occupancy of a shallow, solvent-exposed canyon at the presumptive Gα-binding interface of its RGS-box. This structural region, homologous to the RGS4 site originally identified in 1997 as interacting with the Gα_i1_ switch I region (49), lacks the deep hydrophobic pockets that typically support high-affinity ligand binding. Despite this, we identified and validated at least one chemotype -- 1,2,4-triazolo[3,4-b][1,3,4]thiadiazines -- that reproducibly inhibits RGS14 GAP activity in both fluorescence-based and radioactive single-turnover assays. These compounds have calculated properties consistent with favorable physicochemical and ADME characteristics, with at least one representative demonstrating preclinical CNS penetrance *in vivo*.

Interestingly, this shallow canyon of the RGS14 RGS-box is predicted to support an ambidextrous binding mode, in which chemically distinct terminal substituents (R1 and R2) are predicted to alternately engage either side of the binding groove while maintaining a consistent central scaffold orientation. Structural studies of other proteins have revealed other striking examples where a single small molecule or hormone can engage its target binding site in two flipped or ambidextrous orientations, providing strong precedent for the binding behavior predicted for these RGS14 inhibitors. In a seminal NMR study of Bcl-xL inhibitors, Reibarkh and colleagues demonstrated that one compound exhibited conventional single-mode binding, whereas a closely related molecule bound in multiple orientations, flipping dynamically within the same binding site (70, 71). This ambidextrous binding mode was confirmed by line broadening effects that could not be eliminated even under excess ligand conditions (70, 71). This and other (72–74) structural precedents collectively support the concept that shallow, feature-poor binding surfaces can accommodate small molecules in multiple, nearly 180°-flipped orientations, a phenomenon directly analogous to the binding behavior observed in our system. These converging observations suggest that ambidextrous binding may represent a generalizable strategy for expanding ligand compatibility at structurally permissive but pharmacologically elusive protein surfaces.

That such inhibitors were discovered at all, despite the constraints of a relatively featureless protein– protein interaction surface presented by RGS14, underscores both the value and the limitations of *in silico* docking for this type of drug target. Unlike enzyme active sites or GPCR ligand-binding cavities, the RGS-box offers little more than a shallow groove enriched in non-polar interactions. As such, even the most active triazolothiadiazines scored in a relatively narrow and modest range of predicted Vina binding affinities (-6.4 to - 9.0 kcal/mol). For comparison, we note that an original RGS protein inhibitor identified via physiological and genetic screens (BMS-195270) also scored in this lower range (-7.66 kcal/mol predicted Vina affinity for pose 7 of 2JNU; Supplementary Figure S11), suggesting that these low predicted-affinity values reflect the inherent constraints of the target rather than shortcomings in docking performance. Indeed, our best-performing receptor conformation (pose 7 of PDB id 2JNU) yielded respectable enrichment values (ROC AUC = 0.63, RIE = 2.58), despite lacking the discrimination typically afforded by “lock-and-key” binding sites. These findings collectively highlight that docking scores must be interpreted relative to the structural nature of the receptor, particularly when targeting PPIs, and that functional validation remains essential for distinguishing meaningful interactions from scoring noise.

Looking forward, our results also suggest that deep-learning-based docking methods -- such as the convolutional neural network (CNN) scoring function implemented in GNINA -- may offer particular advantages for shallow or otherwise undruggable targets. As shown in Figure S10, we applied GNINA’s CNN-based affinity predictions to evaluate how adding a methyl ester group at the R3 position influenced predicted binding to the shallow groove of the RGS14 RGS-box. Compared to their unsubstituted R3-free counterparts, the methyl ester-substituted analogs consistently received improved CNN affinity scores, which paralleled their experimentally observed increases in RGS-box GAP inhibitory potency. These improvements likely reflect the CNN model’s ability to integrate nuanced shape and polarity features that are underweighted in traditional scoring functions like those in AutoDock Vina. As such, this study supports the growing utility of AI-augmented docking, particularly in identifying and optimizing ligands for targets traditionally resistant to conventional drug discovery. In the case of the “undruggable” RGS-box target, conserved cysteines distal to the Gα-binding site form a reactive ‘hot-spot’ for allosteric inhibition of the RGS-box/Gα·GTP protein/protein interaction (42). This hot-spot has proven to be an Achilles’ heel for conventional library screening, repeatedly yielding thiol-reactive inhibitory compounds when traditional small-molecule collections are screened *in vitro* for GAP inhibitors (*e.g.*, refs. (35), (42)); in the present study, we briefly encountered this pitfall again (Fig. S12). By focusing virtual docking on the actual PPI groove -- using a boxed volume around the RGS-box’s shallow canyon -- and proactively eliminating thiol-reactive or otherwise promiscuously reactive compounds prior to *in vitro* validation, we can better prioritize non-covalent, chemically tractable scaffolds and accelerate the discovery of more potent GAP activity inhibitors still needed in this space.

## Acknowledgements

Work was supported in part by the US National Institute on Drug Abuse (R01 grant DA048153, to D.P.S), a Team Science award from the HSC Division of Research and Innovation (to K.A.E., L.C.-P., and D.P.S.), the Israel Science Foundation (ISF) and the Azrieli Foundation (grant 3512/19 to M.K.), a grant from the Council for Higher Education through the Data Science Research Center at the University of Haifa (to M.K.), and an HSC Presidential Endowment to the Chair of Pharmacology and Neuroscience (to D.P.S.).

## Methods

### AtomNet® screening and computational assessments of identified compounds

ICM Pocket Finder and fpocket analyses (75, 76) were applied to all twenty low-energy NMR-derived structural conformations reported for the RGS14 RGS-box (PDB id 2JNU; ref. (29)), leading to the identification of pose 5 as having the largest hydrophobic volume for its presumptive Gα-interaction site. This pose 5 receptor grid was entered into the AtomNet® screening protocol against a ∼2.5-million element Enamine in-stock virtual chemical library (circa 2020) as previously described (55). Compounds were scored by AtomNet®, and the top 30,000 hits were filtered using Lipinski’s Rule of Five criteria (in an attempt to avoid future problems with poor absorption or permeation) (64), followed by Butina clustering (77) with ECFP4 Tanimoto coefficient < 0.35 to ensure chemical diversity (78). Ninety-six compounds were selected for testing in the Transcreener® GDP RGScreen™ assay (detailed below), leading to identification of the active triazolothiadiazine derivative Z90276197; subsequent use of the same receptor grid in AtomNet® screening of the ∼16-billion element Enamine REAL library and Transcreener® GDP assay testing of an additional 96 compounds resulted in the identification of 48 additional active triazolothiadiazine derivatives. R-group analyses across these triazolothiadiazines, as well as physicochemical property quantifications and predictions for each compound, were performed using Canvas within the Schrödinger software suite (79) and tested statistically for association with RGS-box inhibition ratio by simple linear regression using GraphPad Prism 10 (Dotmatics; Boston, MA). Principal component analyses of physicochemical properties, and comparisons to those of all FDA-approved compounds from 1951-2021 (obtained from ref. (80)), were performed using an RDKit-evoking (81) Python script “Walter_3D_1.0” (82).

### Transcreener® GDP RGScreen™ assay

Reagents and protocols for the fluorescence anisotropy-based Transcreener® GDP RGScreen™ assay using a rate-altered, double point-mutant, recombinant Gα_i1_ protein have been extensively described previously (43, 44); reactions were always performed so that total GTP converted over the time course was consistently less than 10% of reaction maximum input. Specific methods for its application in assessing RGS14 and RGS12 GAP activity are summarized as follows. Open-reading frames (ORFs) encoding wild-type versions of the human RGS14 RGS-box (aa 46-195 of UniProt O43566) and human RGS12 RGS-box (aa 695-843 of UniProt O14924) were each sub-cloned in-frame with an N-terminal hexahistidine (His_6_) expression tag in the plasmid pET30a, expressed in *E. coli* cultures, and then purified by nickel-nitriloacetate (Ni-NTA) and size-exclusion (Superdex 200) column chromatographies by GenScript using previously described methods (35). These recombinant His_6_-tag fusion proteins were stored at -80 °C in storage buffer (150 mM NaCl, 50 mM Tris-HCl pH 8.0, 10% glycerol) before thawing and use. Cysteine-less versions were similarly produced by ORF site-directed mutagenesis: for RGS14, codons for cysteine residues 47 and 130 (numbered as in PDB id 2JNU) were replaced with serine codons; for RGS12, codons for cysteines 4, 51, 78, and 134 (numbered in relation to PDB id 2EBZ) were replaced with serine codons. Use of Transcreener® GDP-detection reagents to assess compound inhibition of *N. fowleri* Gα7 protein GTPase activity (83) has previously been described (84).

### Cytotoxicity assays

To evaluate cellular toxicity of active compounds, we employed the immortalized human monocytic THP1-Dual™ cell line (InvivoGen) in two orthogonal cytotoxicity assays: ATP-dependent luminescence (CytoTox-Glo) and colorimetric lactate dehydrogenase (LDH) release (CytoTox 96). For each assay, compounds were serially diluted 2-fold starting at 250 μM and added to 96-well flat-bottom tissue culture plates (Thermo Scientific Nunc, Cat# 165306). THP1-Dual™ cells were then seeded at 20,000–30,000 cells per well in 100 μL RPMI 1640 media supplemented with 10% fetal bovine serum (Gibco). Plates were gently mixed and incubated overnight at 37°C in a humidified 5% CO₂ incubator. For the CytoTox-Glo assay (Promega, Cat# G9291), 50 μL of assay reagent was added to each well. Plates were mixed briefly, sealed, and incubated at room temperature for 15 minutes before reading luminescence on a TECAN Infinite M1000 Pro plate reader to quantify dead-cell signal. Subsequently, 50 μL of a digitonin-containing lysis solution was added to each well, followed by a second luminescence read after 15 minutes to assess total cell content. Percent cytotoxicity was calculated per manufacturer instructions. For the CytoTox 96 Non-Radioactive Cytotoxicity Assay (Promega, Cat# G1780), after overnight compound treatment, LDH release into the culture medium was measured by adding the provided substrate mix. Absorbance was read at 490 or 492 nm within one hour of adding the stop solution. Assay optimization and data normalization were conducted according to the manufacturer’s technical manual.

### Pharmacokinetic studies

Single-dose pharmacokinetic (PK) studies were conducted under IACUC-approved protocols of the UNTSCP Preclinical Services laboratory at the University of North Texas Health Science Center. Male Swiss-Webster (CFW) mice (∼30 g; Charles River) were used in both studies, housed in groups of three per cage with *ad libitum* access to food and water and acclimated for a minimum of three days prior to dosing. Each compound was evaluated in a single-dose, time-course design using three mice per time point across seven collection points (0.25, 0.5, 1, 2, 4, 8, and 24 hours post-dose). For Z56944652, mice received a single intraperitoneal (IP) dose of 10 mg/kg (0.2 mL at 1.5 mg/mL). At each designated time point, mice were euthanized via CO_2_ inhalation, and blood was collected by cardiac puncture for plasma isolation (K₂-EDTA tubes). Brains were aseptically removed, flash-frozen, and stored at -80::J°C. For Z55628018, due to poor detection in preliminary studies, the protocol was modified to enhance systemic exposure. Mice were pre-dosed orally with 50 mg/kg of the pan-CYP inhibitor 1-aminobenzotriazole (ABT; ref. (67)) two hours prior to administration of Z55628018. The compound was then administered via IP injection at a single dose of 30 mg/kg (0.2 mL at 1.5 mg/mL). Sample collection and processing procedures were identical to those described above for Z56944652. Despite protocol adjustments, Z55628018 was undetectable in all plasma and brain samples except for one animal at the 1-hour time point, which exhibited a plasma concentration of 0.17 µg/mL (just above the lower limit of detection).

LC-MS/MS analyses were performed at UNTHSC using an Agilent 1260 Infinity HPLC system coupled to an Agilent 6460 Triple Quadrupole mass spectrometer equipped with Jet Stream electrospray ionization (ESI) in positive ion mode. Separation was carried out on a Poroshell 120 EC-C18 column (50 × 3.0 mm, 2.7 µm particle size) maintained at 30::J°C, with a mobile phase consisting of 0.1% formic acid in water (A) and 0.1% formic acid in acetonitrile (B) under a standard gradient elution at a flow rate of 0.5 mL/min. The injection volume was 2 µL. Multiple reaction monitoring (MRM) transitions were optimized for each compound: Z56944652 was detected at m/z 353.1 → 227/132/126, and Z55628018 at m/z 391 → 134.8/247.9/132/121. The internal standard, carbamazepine, was monitored at m/z 237.2 → 167. Plasma samples (30–40 µL) were extracted with 1000–1200 µL of either acetonitrile or a 50:50 acetonitrile:methanol mixture containing carbamazepine (150–500 ng/mL), followed by vortexing, sonication, centrifugation, and supernatant collection for injection. Brain samples were homogenized in organic solvent (2 mL per 0.5 g tissue), vortexed, centrifuged, and processed using the same analytical workflow. Matrix-matched calibration curves were prepared using blank plasma or brain extract and validated over appropriate dynamic ranges: 0.09–46 µg/mL (plasma) and 0.547–70 ng/mL (brain) for Z56944652; 0.1–12.5 µg/mL (plasma) and 0.1–3.13 ng/mL (brain) for Z55628018. Lower limits of quantification (LLOQ) were 0.09 µg/mL and 0.547 ng/mL for Z56944652, and 0.1 µg/mL and 0.1 ng/mL for Z55628018. Retention times were 5.0 ± 0.01 min for Z56944652 and 3.3 ± 0.01 min for Z55628018, with the internal standard eluting at 4.0 ± 0.02 min.

### Vina docking and enrichment calculations

To evaluate and rank each of the twenty low-energy NMR-derived RGS14 and RGS12 RGS-box receptor conformers for their suitability as docking sites for the active triazolothiadiazine series, we first employed GNINA (v1.0; ref. (68)), an AutoDock Vina derivative implementing both Vina scoring and convolutional neural network (CNN)-based scoring functions for docking affinity predictions. GNINA (compiled with CUDA acceleration) was run on a 96 Intel CPU / four Tesla V100 (32 GB HBM2) NVIDIA GPU high-performance computer under Rocky Linux 9.5, initially in unbiased docking mode using the `--autobox_ligand receptor.pdbqt` switch (68) to enable candidate ligands to explore the entire surface of each RGS-box conformer as a potential binding site. For each of the 49 active triazolothiadiazine derivatives, docking was performed across all twenty NMR-derived low-energy structural conformers of the RGS14 RGS-box (PDB id 2JNU) and its paralog RGS12 (PDB id 2EBZ), totaling 40 distinct docking environments. To validate the discriminatory power of the resultant single receptor conformation demonstrating the highest binding affinities across the triazolothiadiazine series, a follow-up enrichment study was conducted using AutoDock Vina (v1.2.5; ref. (69)). Ligands included the 49 triazolothiadiazine actives plus 4160 matched-property decoy molecules drawn from a curated subset of the DUD-E database (85). All ligands were pre-processed using Open Babel (86) to convert from 3D .sdf format and generate .pdbqt files with appropriate Gasteiger partial charges, using a custom Python script built upon Open Babel’s command-line interface.

Docking was conducted in batch mode (with custom Python scripting for GNINA that evoked GNU Parallel parallelization (87); using the `--batch` switch for Vina) for all 40 receptor conformers, with receptor files prepared using Protein Preparation Wizard (under Schrödinger Maestro; ref. (88)) followed by Open Babel processing under physiological pH (pH 7.4) and Gasteiger partial charge assignment to provide appropriately formatted receptor.pdbqt files (verified by manual inspection using ChimeraX; ref. (89)). Docking grid volumes, when formally specified, were defined to fully encompass the Gα-binding interface across the RGS14 and RGS12 conformers. Enrichment performance was assessed via ROC AUC and robust initial enrichment (RIE) metrics using Schrödinger’s Enrichment Calculator (90).

### Single-turnover [**γ** -^33^P]GTP hydrolysis

The single-turnover radiolabeled-GTP hydrolysis assay is the gold standard assay for RGS-box GAP activity, first described using [γ -^32^P]GTP by Berman *et al.* (7) and recently modified for modern performance (91–94) to use readily available [γ-^33^P]GTP substrate and perchloric acid quenching. The assay was performed by first pre-loading wild-type (WT) Gα protein (*i.e.*, 750 nM purified, recombinant, wild-type-sequence human Gα_i1_ protein) with a slight excess (1 μM) of [γ-^33^P]GTP for 15 min at 30 °C. Assay initiation involved adding purified recombinant RGS protein previously pre-incubated for 30 min at 30 °C with a candidate inhibitor. Final concentrations of reactants and reagents were as follows: 500 nM Gα_i1_(WT), 200 nM RGS protein, 100 μM cold (non-radioactive) GTP, and variable concentrations of candidate RGS-box inhibitor (from 0.015 to 1000 μM). This reaction mixture was then incubated on wet-ice (4 °C)for 30 seconds, terminated by adding 3x volume of 5% perchloric acid, and then all except released Pi was precipitated with the addition of neutral (pH 7.5) activated charcoal slurry. Supernatant aliquots were then added to liquid scintillation fluid and [^33^P]-Pi content (*i.e.*, hydrolyzed GTP) was quantified with a scintillation counter with an appropriate β-transmission window.

## Data Availability

All original data have been presented in the main text or supplementary information sections of this manuscript; all structural models used have been publicly available and indicated as such throughout the manuscript.

