## Supplementary Information for "Developing inhibitors of the guanosine triphosphate hydrolysis accelerating activity of Regulator of G protein Signaling-14"

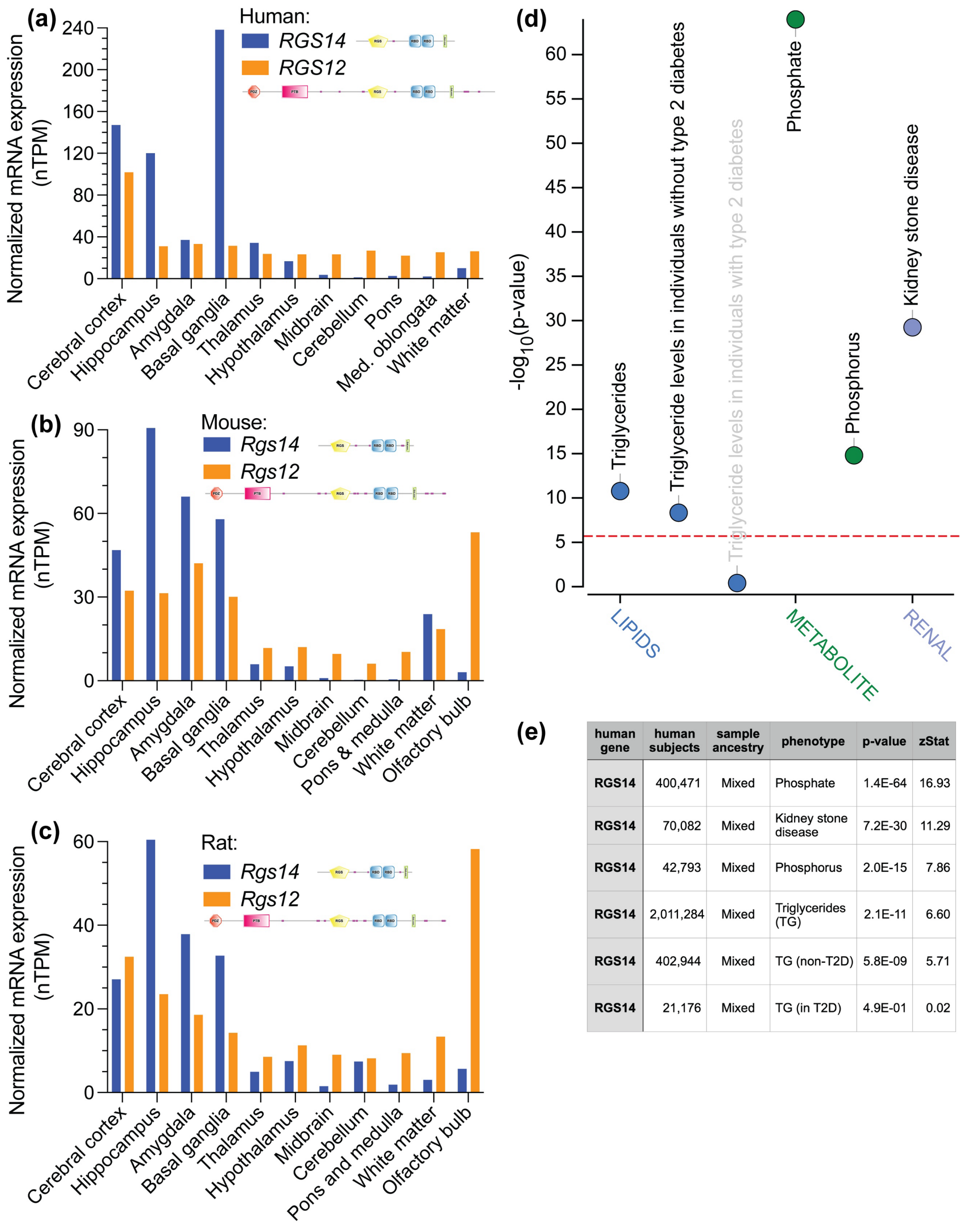

| human gene | human subjects | sample ancestry | phenotype | p-value | zStat |
|---|---|---|---|---|---|
| RGS14 | 400,471 | Mixed | Phosphate | 1.4E-64 | 16.93 |
| RGS14 | 70,082 | Mixed | Kidney stone disease | 7.2E-30 | 11.29 |
| RGS14 | 42,793 | Mixed | Phosphorus | 2.0E-15 | 7.86 |
| RGS14 | 2,011,284 | Mixed | Triglycerides (TG) | 2.1E-11 | 6.60 |
| RGS14 | 402,944 | Mixed | TG (non-T2D) | 5.8E-09 | 5.71 |
| RGS14 | 21,176 | Mixed | TG (in T2D) | 4.9E-01 | 0.02 |

**Supplementary Figure S1. Comparative analyses of RGS14 and RGS12 protein domain organizations, brain-region gene expression patterns, and phenome-wide associations for human RGS14 genetic variation.** Normalized transcript per million (nTPM) values are shown for RGS14 (blue bars) and RGS12 (orange bars) across homologous brain regions in (**a**) human, (**b**) mouse, and (**c**) rat. Human and mouse transcript expression data were obtained from the Human Protein Atlas (HPA) transcriptomics database (1) -- RGS14 (ENSG00000169220) and RGS12 (ENSG00000159788) -- each representing bulk RNA-seq expression across defined anatomical structures. Rat brain region data were curated from BioGPS BioDatabase System (2) dataset BDS_00054, which provides organ- and region-specific nTPM values derived from bulk RNA-seq (including discrete subregions such as the basolateral amygdala, corpus striatum, and subiculum). Region labels in the rat dataset were manually harmonized to match those in the human and mouse panels. Specifically, "corpus striatum" and "nucleus accumbens" were merged into the "Basal ganglia" category; "subiculum" was grouped with the "Hippocampus"; and "brainstem" was used to approximate "Pons and medulla". The olfactory bulb, absent from human HPA data, is shown for rat and mouse only. Expression values represent the arithmetic mean across multiple biological replicates per brain region, where applicable. Domain layouts within graph legends were generated from SMART outputs using the following UniProt entries: RGS14_HUMAN, RGS12_HUMAN, RGS14_MOUSE, RGS12_MOUSE, RGS14_RAT, and Swiss-Prot id O08774 for rat RGS12 protein sequence, respectively. Protein domains are color-coded: PDZ (orange hexagon), PTB (rose-pink rectangle), RGS-box (yellow pentagon), Ras-binding domain (RBD, cyan square), GoLoco (lime green rectangle), and predicted low-complexity regions (pink lines). Panels (**d**) and (**e**) represent graphical and numeric readouts, respectively, obtained from the Type 2 Diabetes Knowledge Portal (https://t2d.hugeamp.org; RRID: SCR_003743; ref. (3)) when queried for phenotypic associations to common human RGS14 gene sequence variations, calculated from bottom-line genetic associations using the MAGMA (Multi-marker Analysis of GenoMic Annotation) method (4). The generally accepted threshold for significance of MAGMA results ($p \leq 2.5 \times 10^{-6}$) is denoted with a red dotted line in the phenome-wide association (PheWAS) plot of panel (**d**), highlighting that the triglyceride levels in individuals without type 2 diabetes are significantly associated with human RGS14 gene variations (p-value $5.8 \times 10^{-9}$), but not triglyceride levels in individuals with type 2 diabetes. Sample size ("*human subjects*") in panel (**e**) refers to the number of individuals whose genotypes contributed to the bottom-line analysis of genetic associations for that phenotype. The MAGMA method's "Z-stat" is a standardized effect size from gene-level regression, indicating the strength and direction of association; positive values suggest stronger associations with increased trait values (4).

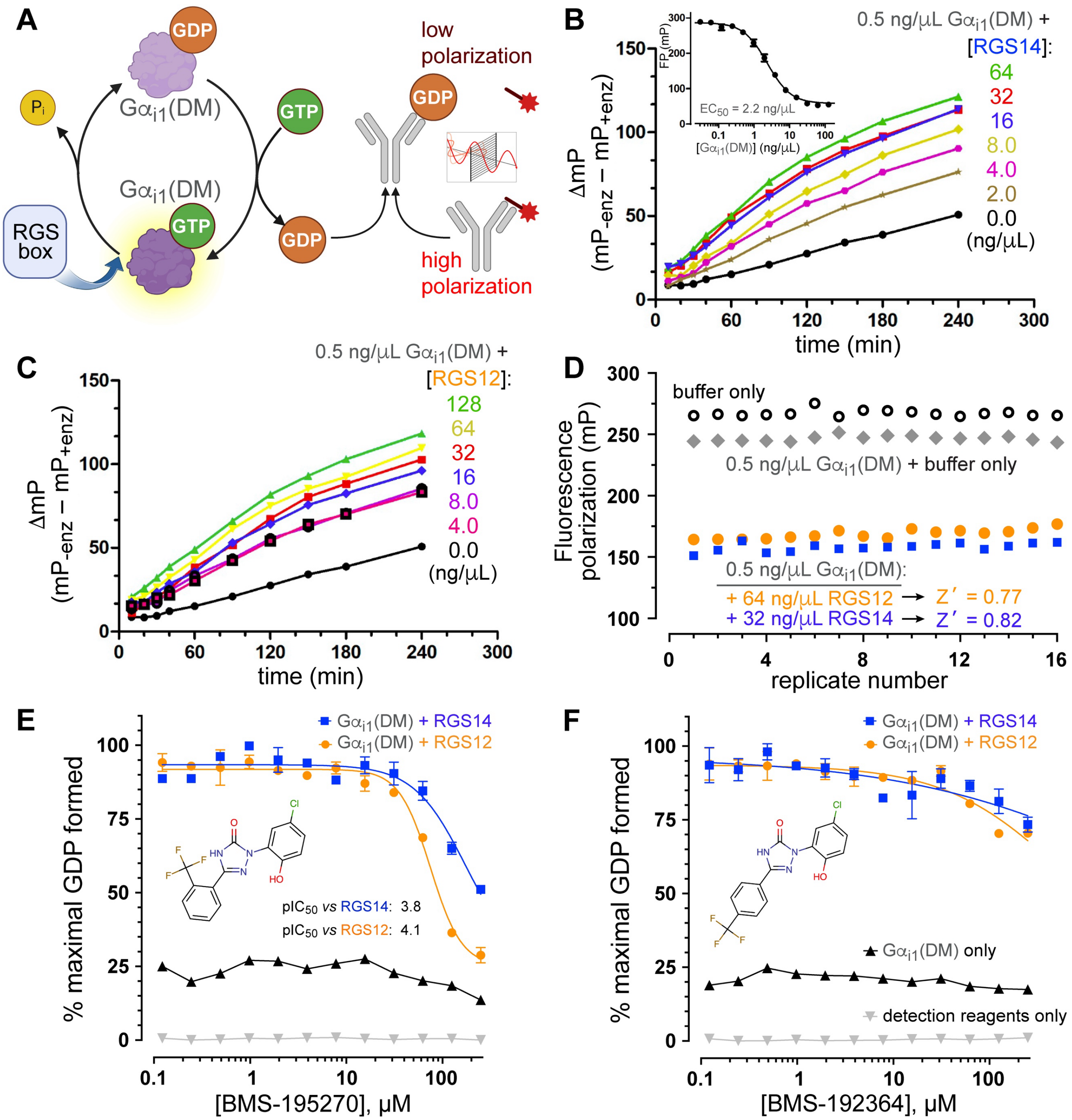

**Supplementary Figure S2. Using the Transcreener® GDP RGScreen™ assay to measure $G\alpha_{i1}$(DM) GTPase activity in the presence of wild-type RGS14 and RGS12 RGS-box proteins. (A)** Schematic representation of the Transcreener® GDP fluorescence polarization immunoassay as applied to measuring steady-state guanosine triphosphate (GTP) hydrolysis activity (and resultant GDP production) by a rate-altered, double point-mutant ("DM") form of $G\alpha_{i1}$ protein when its intrinsic GTP hydrolysis rate is accelerated by the GAP activity of an RGS-box protein (reaction details extensively explored in refs. (5, 6)). Fluorescent tracer is illustrated with a jagged red oval; when bound to the GDP-selective monoclonal antibody, emitted light remains polarized, whereas there is low polarization of emitted light when this tracer is displaced by free GDP. **(B, C)** Recombinant wild-type RGS14 RGS-box protein (panel B) and wild-type RGS12 RGS-box protein (panel C) are both observed, in a concentration-dependent fashion, to increase GDP production by 0.5 ng/μL $G\alpha_{i1}$(DM) protein (representing the $EC_{20}$ for $G\alpha_{i1}$(DM) in a 20 μL final reaction volume), as measured by the change in polarization (ΔmP) at each indicated time point ("enz" = $G\alpha_{i1}$(DM) protein a.k.a. "enzyme"). *Inset to panel B:* Initial titration (to establish $EC_{20}$) of recombinant $G\alpha_{i1}$(DM) protein into Transcreener® GDP detection reagents and measurement of fluorescence polarization output (in units of milliP or "milli-polarization units") after reaction incubation for 120 minutes at 30 °C. **(D)** Sixteen (16) replicate measures of final fluorescence polarization were obtained after 180 min with indicated reaction conditions to calculate the assay's Z' factor: a dimensionless statistic (7) assessing both signal dynamic range and variation within data from both positive and negative controls. An observed Z' factor > 0.5 is generally considered permissible for assay launch in a screening campaign (7). **(E, F)** Tests of concentration-dependent inhibition of RGS14 (blue) and RGS12 (orange) RGS-box GAP activity on $G\alpha_{i1}$(DM) protein by Bristol Myers Squibb-identified 1,3-diaryl 1,2,4-(4H)-triazol-5-ones: BMS-195270 (panel E; estimated $IC_{50}^{RGS14}$ of 174 μM [$pIC_{50}^{RGS14}$ of 3.8] and estimated $IC_{50}^{RGS12}$ of 75 μM [$pIC_{50}^{RGS14}$ of 4.1] and BMS-192364 (panel F; considered inactive against RGS14 and RGS12)).

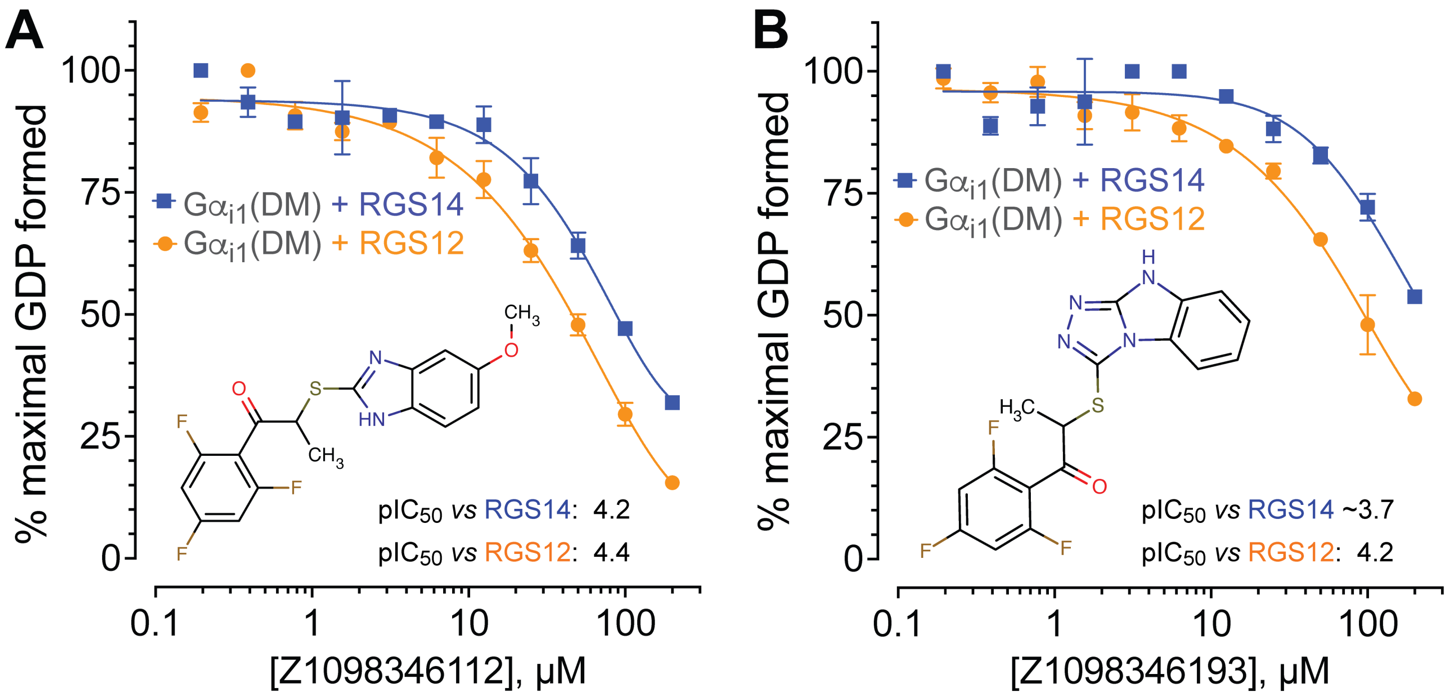

**Supplementary Figure S3. RGS14 and RGS12 inhibition by initial hit compound Z1098346112 and the related compound Z1098346193.** (**A**) Concentration-dependent inhibition of RGS14 (**blue**) and RGS12 (**orange**) RGS-box GAP activity by original hit compound 2-[(5-methoxy-1H-1,3-benzodiazol-2-yl)sulfanyl]-1-(2,4,6-trifluorophenyl) propane-1-one (Z1098346112; estimated $pIC_{50}^{RGS14}$ of 4.2 and estimated $IC_{50}^{RGS12}$ of 4.4) (from ref. (8)) and the sole Enamine-sourced analog found with activity, 2-(2,4,5,7-tetraaza-tricyclo[6.4.0.0$^{2}$,$^{6}$]dodeca-1(12),3,5,8,10-pentaen-3-ylsulfanyl)-1-(2,4,6-trifluorophenyl)propan-1-one (Z1098346193; estimated $pIC_{50}^{RGS14}$ of ~3.7 and estimated $pIC_{50}^{RGS12}$ of 4.2), obtained after having screened >80 related, but ultimately inactive, benzimidazoles (listed in Supplementary Table S1).

| SMILES | Enamine identifier | Tanimoto coefficient (ECFP4, *vs* Z1098346112) | $pIC_{50}$ (RGS14) | 2JNU pose 7 predicted Vina affinity (kcal/mol) |
|---|---|---|---|---|
| **Initial docking hit, initial followup active, and initial (negative) SAR around benzimidazoles** | | | | |
| COC1=CC2=C(C=C1)N=C(SC(C)C(=O)C1=C(F)C=C(F)C=C1F)N2 | **Z1098346112** | **1.00** | **4.2** | **-7.34** |
| CC(SC1=NNC2=NC3=C(C=CC=C3)N21)C(=O)C1=C(F)C=C(F)C=C1F | **Z1098346193** | **0.43** | **~3.7** | **-7.87** |
| COC1=CC=C(C(=O)C(C)SC2=NC3=C(C=C(OC)C=C3)N2)C=C1 | Z16077705 | 0.71 | <3.0 | -6.88 |
| COC1=CC2=C(C=C1)N=C(SC(C)C(=O)C1=CC=CC=C1)N2 | Z16077228 | 0.69 | <3.0 | -6.92 |
| COC1=CC2=C(C=C1)N=C(SCCC(=O)C1=CC=C(F)C=C1)N2 | Z16077963 | 0.52 | <3.0 | -6.78 |
| COC1=CC2=C(C=C1)N=C(SCC(=O)C1=CC=C(O)C=C1)N2 | Z112468740 | 0.47 | <3.0 | -6.79 |
| COC1=CC2=C(C=C1)N=C(SCC(O)C1=CC=CC=C1)N2 | Z56784116 | 0.45 | <3.0 | -6.23 |
| CCOC1=CC2=C(C=C1)NC(SCCC(=O)C1=CC=C(F)C=C1)=N2 | Z19828508 | 0.33 | <3.0 | -6.51 |
| **Benzimidazoles that interfere with Transcreener® GDP detection reagents** | | | | |
| CC(SC1=NC=2C=CC=CC2N1C(F)F)C(=O)C=3C(F)=CC(F)=CC3F | Z1098351974 | 0.44 | <3.0 | -7.28 |
| CCOC(=O)C=1C=CC=2N(CC)C(SC(C)C(=O)C=3C(F)=CC(F)=CC3F)=NC2C1 | Z1098347818 | 0.42 | <3.0 | -7.02 |
| CCCCN1C(SC(C)C(=O)C=2C(F)=CC(F)=CC2F)=NC=3C=C(C=CC13)S(=O)(=O)N(C)C | Z1098346992 | 0.36 | <3.0 | -7.27 |
| CCOC(=O)C=1C=CC=2N(CC)C(SC(C)C(=O)C=3C=CC(NC(=O)C(C)C)=CC3)=NC2C1 | Z65654925 | 0.24 | <3.0 | -6.94 |
| **Benzimidazoles that exhibit non-specific inhibition (affecting Gαi1(DM) GTPase activity alone)** | | | | |
| COC=1C=CC=2N=C(SC(C)C(=O)C=3C=CC(NC(=O)C)=CC3)NC2C1 | Z16077185 | 0.62 | <3.0 | -7.20 |
| COC=1C=CC=2N=C(SC(C)C(=O)C=3C=CC=4N(CCC4C3)C(=O)C)NC2C1 | Z165591262 | 0.54 | <3.0 | -7.58 |
| COC=1C=CC=2N=C(SCC(=O)C=3C=CC(F)=CC3)NC2C1 | Z16077977 | 0.53 | <3.0 | -6.76 |
| COC=1C=CC(=CC1)N2C(SC(C)C(=O)C=3C=C(C)C(C)=CC3C)=NC=4C=CC=CC24 | Z19576531 | 0.32 | <3.0 | -7.23 |
| CC(C)C=1C=CC(C(=O)C(C)SC2=NC=3C=CC=CC3N2C(C)C)=C(C1)C(C)C | Z54156331 | 0.26 | <3.0 | -7.43 |
| CCC=1C=CC(=CC1)C(=O)C(C)SC2=NC=3C=C(C=CC3N2CC)S(=O)(=O)N | Z51418852 | 0.24 | <3.0 | -7.29 |
| CCN1C(SC(C)C(=O)C=2C=CC=3N(CCC3C2)C(=O)C)=NC=4C=CC=CC14 | Z165501154 | 0.23 | <3.0 | -7.04 |
| **Inactive unsubstituted benzimidazoles** | | | | |
| COC=1C=CC=2N=C(SC(C)C(=O)C=3C=CC(F)=CC3F)NC2C1 | Z167736700 | 0.78 | <3.0 | -7.15 |
| COC=1C=CC=2N=C(SC(C)C(=O)C=3C=CC(F)=CC3)NC2C1 | Z16077798 | 0.75 | <3.0 | -7.05 |
| COC=1C=CC=2N=C(SC(C)C(=O)C=3C=CC(F)=C(F)C3)NC2C1 | Z16077536 | 0.73 | <3.0 | -7.41 |
| COC=1C=CC=2N=C(SC(C)C(=O)C=3C=CC=C(F)C3)NC2C1 | Z5032180758 | 0.73 | <3.0 | -7.03 |
| COC=1C=CC=2N=C(SC(C)C(=O)NC=3C=CC(F)=CC3F)NC2C1 | Z112468618 | 0.68 | <3.0 | -7.46 |
| COC=1C=CC(C(=O)C(C)SC2=NC=3C=CC(OC)=CC3N2)=C(OC)C1 | Z16077640 | 0.67 | <3.0 | -6.78 |
| COC=1C=CC=2N=C(SC(C)C(=O)C=3C=CC(OC(F)F)=CC3)NC2C1 | Z16077231 | 0.67 | <3.0 | -7.08 |
| COC=1C=CC=2N=C(SC(C)C(=O)C=3C=C(C)C(C)=CC3C)NC2C1 | Z16077531 | 0.66 | <3.0 | -6.91 |
| COC=1C=CC=2N=C(SC(C)C(=O)C=3C=CC(Br)=CC3)NC2C1 | Z16077584 | 0.66 | <3.0 | -6.86 |
| CC(SC1=NC=2C=C(C)C=CC2N1)C(=O)C=3C(F)=CC(F)=CC3F | Z1098347474 | 0.65 | <3.0 | -7.41 |
| COC=1C=CC=2N=C(SC(C)C(=O)NC=3C=C(F)C=CC3F)NC2C1 | Z112468620 | 0.65 | <3.0 | -7.36 |
| COC=1C=CC=2N=C(SC(C)C(=O)C=3C=CC(C)=C(C)C3)NC2C1 | Z16077488 | 0.65 | <3.0 | -7.29 |
| COC=1C=CC=2N=C(SC(C)C(=O)C=3C=CC(OC)=C(OC)C3)NC2C1 | Z940567722 | 0.65 | <3.0 | -7.02 |
| COC=1C=CC=2N=C(SC(C)C(=O)N(C)C)NC2C1 | Z92077129 | 0.65 | <3.0 | -5.81 |
| COC=1C=CC=2N=C(SC(C)C(=O)C=3C=CC=C(Cl)C3)NC2C1 | Z16077229 | 0.64 | <3.0 | -6.98 |
| COC=1C=CC=2N=C(SC(C)C(=O)C=3C=C(C)C=CC3C)NC2C1 | Z16077483 | 0.64 | <3.0 | -6.98 |
| CCC=1C=CC(=CC1)C(=O)C(C)SC2=NC=3C=CC(OC)=CC3N2 | Z16077530 | 0.64 | <3.0 | -7.12 |
| COC=1C=CC=2N=C(SC(C)C(=O)C=3C=CC(=CC3)C(C)(C)C)NC2C1 | Z16077565 | 0.64 | <3.0 | -6.97 |
| COC=1C=CC=2N=C(SC(C)C(=O)C3=CC=CO3)NC2C1 | Z3277796272 | 0.64 | <3.0 | -6.69 |
| COC=1C=CC=2N=C(SC(C)C(=O)C=3C=CC=CC3C(F)(F)F)NC2C1 | Z5032180788 | 0.63 | <3.0 | -7.20 |
| COC=1C=CC=2N=C(SC(C)C(=O)C=3NC(C)=C(C(=O)C)C3C)NC2C1 | Z98717471 | 0.63 | <3.0 | -6.76 |
| COC=1C=CC=2N=C(SC(C)C(=O)C=3C=CC=4CCCC4C3)NC2C1 | Z16076970 | 0.61 | <3.0 | -7.53 |
| COC=1C=CC=2N=C(SC(C)C(=O)C=3C=CC(NC(=O)C(C)C)=CC3)NC2C1 | Z65841956 | 0.61 | <3.0 | -7.09 |
| COC=1C=CC=2N=C(SC(C)C(=O)C=3C=CC(NC(=O)C(C)(C)C)=CC3)NC2C1 | Z50965385 | 0.59 | <3.0 | -7.21 |
| COC=1C=CC=2N=C(SC(C)C(=O)NC=3C=CC(C)=CC3)NC2C1 | Z16077036 | 0.58 | <3.0 | -7.01 |
| CCCC(=O)NC=1C=CC(=CC1)C(=O)C(C)SC2=NC=3C=CC(OC)=CC3N2 | Z50965416 | 0.57 | <3.0 | -6.97 |
| CCC(SC1=NC=2C=CC(OC)=CC2N1)C(=O)C=3C=CC(OC)=CC3 | Z5032180767 | 0.56 | <3.0 | -6.87 |
| COC=1C=CC=2N=C(SC(C(C)C)C(=O)OC(C)(C)C)NC2C1 | Z5032180761 | 0.55 | <3.0 | -6.12 |
| CCC(SC1=NC=2C=CC(OC)=CC2N1)C(=O)C=3C=CC=CC3 | Z447938100 | 0.54 | <3.0 | -7.00 |
| CCC(SC1=NC=2C=CC(OC)=CC2N1)C(=O)C=3C=CC(Cl)=CC3 | Z5032180785 | 0.52 | <3.0 | -6.87 |
| CC(SC1=NC=2C=C(C)C=CC2N1)C(=O)C=3C=CC(F)=CC3F | Z167804418 | 0.50 | <3.0 | -7.24 |
| CCC(SC1=NC=2C=CC(OC)=CC2N1)C(=O)C=3C=CC(NC(=O)C)=CC3 | Z1340653159 | 0.49 | <3.0 | -7.33 |
| COC=1C=CC=2NC(SC(C(=O)N)C=3C=CC(F)=CC3)=NC2C1 | Z234729962 | 0.49 | <3.0 | -7.31 |
| CC(SC1=NC=2C=C(C)C=CC2N1)C(=O)C=3C=CC(F)=CC3 | Z18443583 | 0.48 | <3.0 | -7.17 |
| CC(SC1=NC=2C=C(C)C=CC2N1)C(=O)C=3C=CC(OC(F)F)=CC3 | Z18443016 | 0.46 | <3.0 | -7.19 |
| CC(SC1=NC=2C=C(C)C=CC2N1)C(=O)C=3C=CC=C(F)C3 | Z5032180743 | 0.46 | <3.0 | -7.16 |
| COC=1C=CC(=CC1Cl)C(=O)C(C)SC2=NC=3C=C(C)C=CC3N2 | Z5032180749 | 0.43 | <3.0 | -7.27 |
| CCC=1C=CC(=CC1)C(=O)C(C)SC2=NC=3C=C(C)C=CC3N2 | Z18443315 | 0.39 | <3.0 | -7.27 |
| COC=1C=CC=2N=C(SCCOCC3CCCO3)NC2C1 | Z1258859165 | 0.38 | <3.0 | -5.82 |
| COC=1C=CC=2NC(SCC(=O)C=3C=CC(O)=C(O)C3)=NC2C1 | Z223822242 | 0.37 | <3.0 | -7.02 |
| CCOC=1C=CC=2N=C(SCC(=O)C=3C=CC(CC)=CC3)NC2C1 | Z55030990 | 0.36 | <3.0 | -6.71 |
| CC(SC1=NC=2C=C(C)C=CC2N1)C(=O)C=3C=CC=4N(CCC4C3)C(=O)C | Z165332254 | 0.34 | <3.0 | -7.68 |
| **Inactive N-substituted benzimidazoles** | | | | |
| COC=1C=CC(=CC1)N2C(SC(C)C(=O)C=3C(F)=CC(F)=CC3F)=NC=4C=CC=CC24 | Z5032180751 | 0.52 | <3.0 | -7.27 |
| CCN1C(SC(C)C(=O)C=2C(F)=CC(F)=CC2F)=NC=3C=CC=CC13 | Z1098345881 | 0.45 | <3.0 | -6.94 |
| CCN1C(SC(C)C(=O)C=2C(F)=CC(F)=CC2F)=NC=3C=C(C=CC13)S(=O)(=O)N4CCCCC4 | Z5032180738 | 0.38 | <3.0 | -7.73 |
| CCN1C(SC(C)C(=O)C=2C=CC(OC)=CC2)=NC=3C=C(C=CC13)S(=O)(=O)N | Z51419018 | 0.33 | <3.0 | -7.01 |
| COC=1C=CC(=CC1)N2C(SC(C)C(=O)C=3C=C(C)C=CC3C)=NC=4C=CC=CC24 | Z19576483 | 0.32 | <3.0 | -6.73 |
| COCCN1C(SC(C)C(=O)C=2C=CC(OC)=CC2)=NC=3C=CC=CC13 | Z16216221 | 0.32 | <3.0 | -6.59 |
| CCN1C(SC(C)C(=O)C=2C=CC(F)=CC2)=NC=3C=C(C=CC13)S(=O)(=O)N | Z51419107 | 0.29 | <3.0 | -7.05 |
| COC=1C=CC(=CC1)C(SC2=NC=3C=CC=CC3N2C)C(=O)C=4C=CC(OC)=CC4 | Z15478257 | 0.28 | <3.0 | -7.43 |
| CCOC=1C=CC(=CC1)C(=O)C(C)SC2=NC=3C=C(C=CC3N2CC)S(=O)(=O)N | Z51419689 | 0.28 | <3.0 | -6.80 |
| CCOC=1C=CC(=CC1)C(=O)C(C)SC2=NC=3C=CC=CC3N2C | Z15479255 | 0.27 | <3.0 | -6.77 |
| COCCN1C(SC(C)C(=O)C=2C=CC(Br)=CC2)=NC=3C=CC=CC13 | Z16216100 | 0.25 | <3.0 | -6.51 |
| CCCCN1C(SC(C(=O)C=2C=CC(OC)=CC2)C=3C=CC(OC)=CC3)=NC=4C=C(C=CC14)S(=O)(=O)N | Z19162246 | 0.24 | <3.0 | -7.01 |
| CCN1C(SC(C)C(=O)C=2C=CC(Cl)=CC2)=NC=3C=C(C=CC13)S(=O)(=O)N | Z51418567 | 0.24 | <3.0 | -7.06 |
| CCN1C(SC(C)C(=O)C=2C=CC(Br)=CC2)=NC=3C=C(C=CC13)S(=O)(=O)N | Z51418902 | 0.24 | <3.0 | -7.04 |
| CCN1C(SC(C)C(=O)C=2C=CC=C(Cl)C2)=NC=3C=C(C=CC13)S(=O)(=O)N | Z51418566 | 0.23 | <3.0 | -6.98 |
| CCN1C(SC(C)C(=O)C=2C=C(C)C=CC2C)=NC=3C=C(C=CC13)S(=O)(=O)N | Z51418807 | 0.23 | <3.0 | -6.54 |
| CCN1C(SC(C)C(=O)C=2C=CC(NS(=O)(=O)C)=CC2)=NC=3C=CC=CC13 | Z50932618 | 0.23 | <3.0 | -6.73 |
| COCCN1C(SC(C)C(=O)C=2C=CC=3N(CCC3C2)C(=O)C)=NC=4C=CC=CC14 | Z165550978 | 0.23 | <3.0 | -7.00 |
| CCN1C(SC(C)C(=O)C=2C=CC(NC(=O)C)=CC2)=NC=3C=C(C=CC13)S(=O)(=O)N | Z51418527 | 0.23 | <3.0 | -7.23 |
| CCN1C(SC(C)C(=O)C=2C=CC=3CCCC3C2)=NC=4C=C(C=CC14)S(=O)(=O)N | Z51418316 | 0.22 | <3.0 | -7.13 |
| CCN1C(SC(C)C(=O)C=2C=CC=CC2)=NC=3C=C(C=CC13)S(=O)(=O)N4CCOCC4 | Z16669878 | 0.22 | <3.0 | -7.43 |
| CC(SC1=NC=2C=CC=CC2N1C)C(=O)C=3C=CC(NS(=O)(=O)C)=CC3 | Z50932699 | 0.22 | <3.0 | -6.93 |
| CCN1C(SC(C)C(=O)C=2C=CC=3CCCC3C2)=NC=4C=C(C=CC14)S(=O)(=O)N5CCOCC5 | Z16669629 | 0.21 | <3.0 | -7.57 |

**Supplementary Table S1. Chemical, computational, and experimental characteristics of benzimidazole-based RGS14 inhibitor candidates.** This table summarizes the structural and screening data for benzimidazole derivatives tested during early structure-activity relationship (SAR) exploration. Columns include: the compound SMILES string, the Enamine identifier, the Tanimoto similarity coefficient (ECFP4) *versus* initial active benzimidazole derivative Z1098346112, experimentally determined $pIC_{50}$ values against RGS14-catalyzed GTP hydrolysis (where available), and the AutoDock Vina-predicted binding affinity (in kcal/mol) for pose 7 of the RGS14 RGS-box (PDB id 2JNU). Compounds are grouped by classification based on experimental outcomes, including initial actives, signal-interfering compounds, non-specific inhibitors of $G\alpha_{i1}$(DM) GTPase activity, and inactive analogs with either unsubstituted or N-substituted benzimidazole scaffolds. Where applicable, lack of $pIC_{50}$ values reflects either inactivity or interference with assay components.

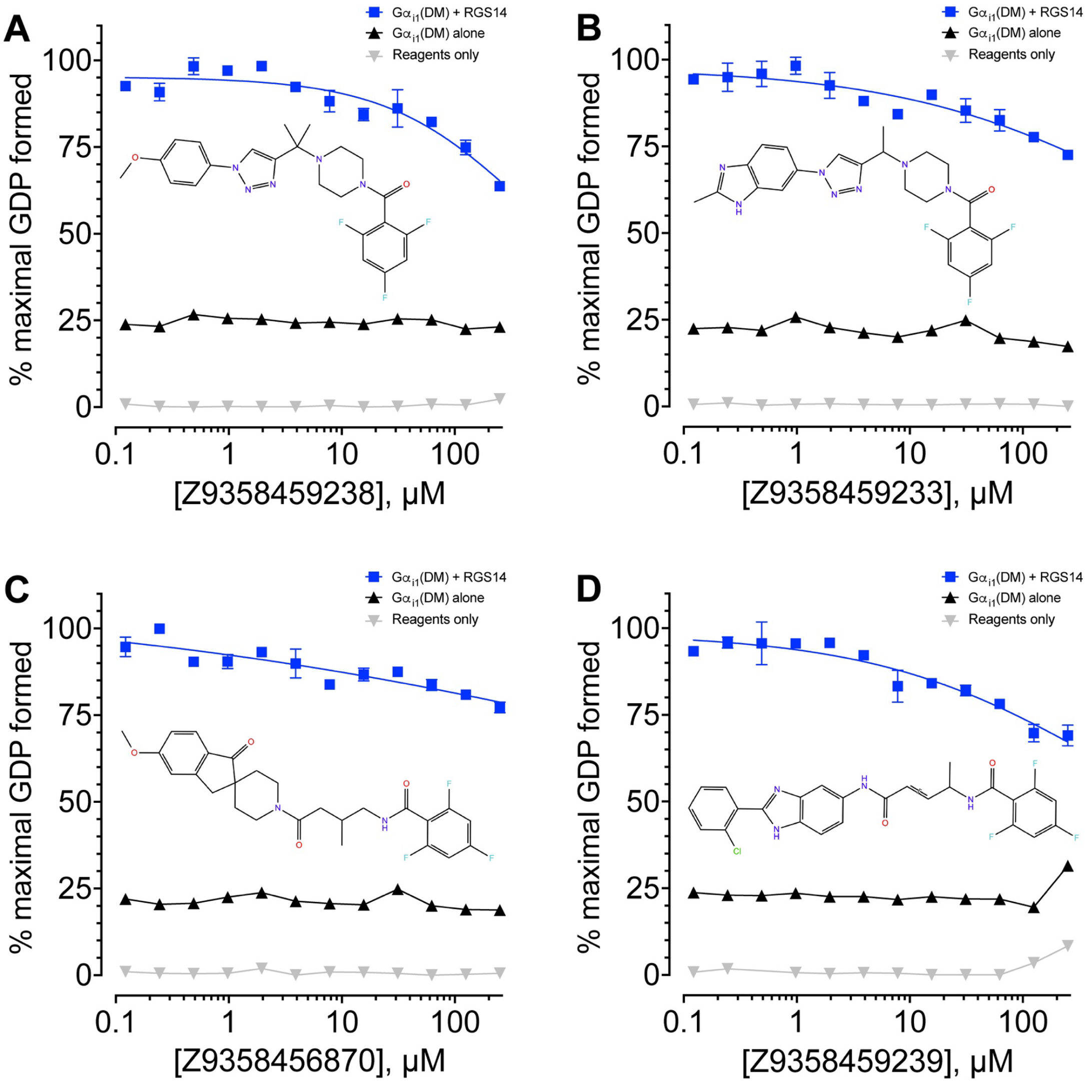

**Supplementary Figure S4. Multi-ringed 2,4,6-trifluorobenzoyl analogs fail to inhibit RGS14 GAP activity.**
(**A-D**) Transcreener® GDP-detection RGScreen™ assays assessing the effect of four multi-ringed, non-benzimidazole analogs -- each containing a 2,4,6-trifluorobenzoyl substituent -- on GTP hydrolysis catalyzed by $G\alpha_{i1}$(DM) protein in the presence or absence of recombinant RGS14 RGS-box protein. Structures were selected based on shared substructure with the active benzimidazole compounds Z1098346112 and Z1098346193, both of which possess a (2,4,6-trifluorophenyl)propan-1-one moiety. Despite their similar size and functional group presentation, none of these four analogs exhibited meaningful inhibition of RGS14-accelerated GAP activity across the tested concentration range. Negative controls ($G\alpha_{i1}$(DM) alone and reagents only) are shown for comparison.

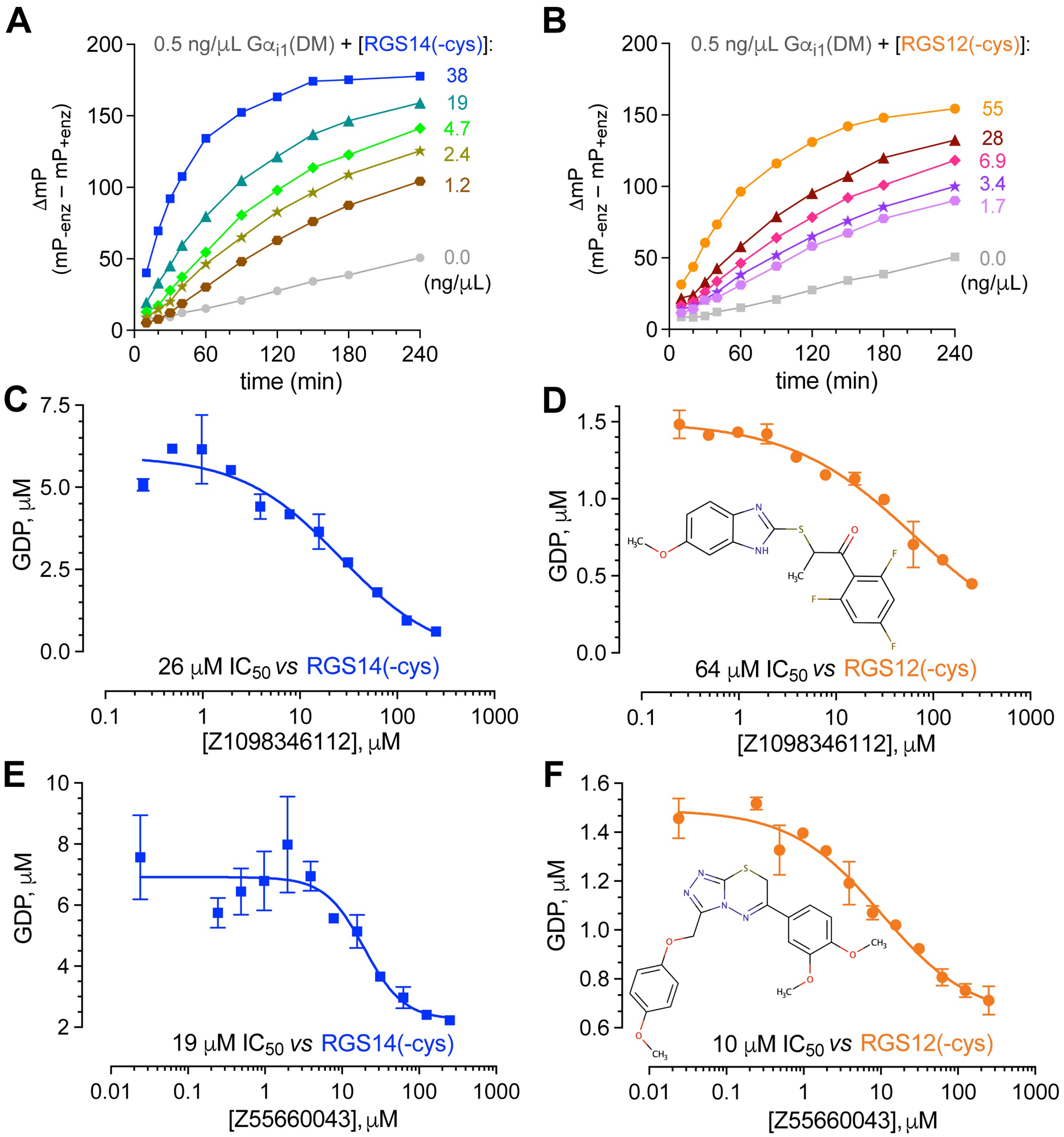

**Supplementary Figure S5. RGS-box inhibitors of the benzimidazole and 1,2,4-triazolo[3,4-b][1,3,4]thiadiazine series inhibit the GAP activities of cysteine-less RGS-box proteins. (A,B)** Recombinant, cysteine residue-lacking ("-cys") RGS14 RGS-box protein (panel A) and cysteine residue-lacking RGS12 RGS-box protein (panel A) were both observed, in a concentration-dependent fashion (indicated in ng/μL by color), to increase steady-state GDP production by 0.5 ng/μL $G\alpha_{i1}$(DM) protein in a 20 μL final reaction volume, as measured by the Transcreener® GDP-detection RGScreen™ assay and detected by the change in polarization (ΔmP) at each indicated time point ("enz" = $G\alpha_{i1}$(DM) protein a.k.a. "enzyme"). (**C,D**) End-point GDP production measured at 120 min in the presence of indicated concentrations of initial hit compound Z1098346112, as produced by the $G\alpha_{i1}$(DM) subunit and accelerated by the GAP activity of cysteine residue-lacking RGS14 RGS-box protein ("RGS14(-cys)"; panel C) or an RGS12 RGS-box recombinant protein lacking all wild-type cysteine residues ("RGS12(-cys)"; panel D). (**E,F**) GDP production in the presence of indicated concentrations of triazolothiadiazine compound Z55660043, as produced in reactions similar to panels C and D, respectively. Using standard curves of mP signal from defined GDP input, Transcreener® GDP detection output was converted to GDP produced (y-axes).

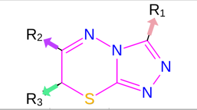

| SMILES | Enamine Name | R3 | R2 | R1 | RGS14 pIC50 | RGS12 pIC50 | Ratio | MW (Da) | cLogP (o/w) | TPSA (Å²) | accept H-bond | donor H-bond | Rotatable bond count | $Fsp^3$ |
|---|---|---|---|---|---|---|---|---|---|---|---|---|---|---|
| O=C([O-])C[C@@H]1Sc2nnc(COc3ccc(Cl)cc3)n2N=C1c1ccccc1 | Z90276197 | 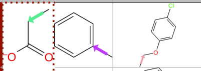 | | | 4.09 | 4.37 | 0.94 | 414.9 | 2.171 | 89.3 | 6 | 2 | 6 | 0.11 |
| O=C([O-])C[C@@H]1Sc2nnc([C@H]3COc4ccccc4O3)n2N=C1c1cccs1 | Z90275816 | 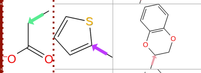 | | | 4.69 | 4.35 | **1.08** | 414.5 | 1.362 | 98.5 | 7 | 2 | 4 | 0.17 |
| COc1ccc(Cc2nnc3n2N=C(c2ccccc2)[C@H](CC(=O)[O-])S3)cc1OC | Z90277093 | 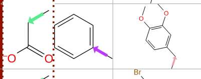 | | | 4.56 | 5.17 | 0.88 | 424.5 | 1.569 | 98.5 | 7 | 2 | 4 | 0.19 |
| O=C([O-])C[C@@H]1Sc2nnc(Cc3ccc(Br)cc3)n2N=C1c1ccc(Cl)cc1 | Z90276771 | 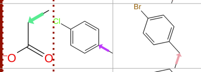 | | | 4.53 | 4.96 | 0.91 | 477.8 | 3.108 | 80.0 | 5 | 2 | 5 | 0.11 |
| O=C([O-])C[C@@H]1Sc2nnc(CCC3CCCCC3)n2N=C1c1ccccc1 | Z90277285 | 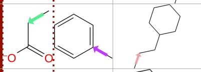 | | | 4.42 | 4.62 | 0.96 | 384.5 | 2.796 | 80.0 | 5 | 2 | 6 | 0.45 |
| COc1ccccc1-c1nnc2n1N=C(c1cccs1)[C@@H](CC(=O)[O-])S2 | Z90275304 | 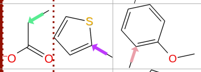 | | | 4.28 | 4.42 | 0.97 | 386.4 | 1.707 | 89.3 | 6 | 2 | 5 | 0.12 |
| O=C([O-])C[C@@H]1Sc2nnc([C@@H]3CCCO3)n2N=C1c1ccc(Cl)cc1 | Z90278051 | 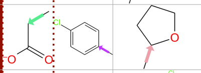 | | | 4.25 | 3.97 | **1.07** | 378.8 | 1.097 | 89.3 | 6 | 2 | 4 | 0.31 |
| Cc1ccc(C2=Nn3c(COc4ccc(Cl)cc4)nnc3S[C@@H]2CC(=O)[O-])cc1 | Z90276194 | 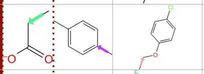 | | | 4.21 | 4.21 | **1.00** | 428.9 | 2.684 | 89.3 | 6 | 2 | 6 | 0.15 |
| Cc1ccc(C2=Nn3c(Cc4ccc(F)cc4)nnc3S[C@@H]2CC(=O)[O-])cc1 | Z90278242 | 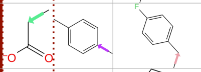 | | | 4.11 | 4.19 | 0.98 | 396.4 | 2.540 | 80.0 | 5 | 2 | 5 | 0.15 |
| O=C([O-])C[C@@H]1Sc2nnc(-c3ccco3)n2N=C1c1ccccc1 | Z90276389 | 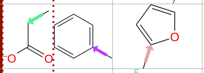 | | | 4.10 | 4.08 | **1.00** | 340.4 | 1.012 | 93.2 | 5 | 2 | 4 | 0.06 |
| O=C([O-])C[C@H]1Sc2nnc(Cc3ccc(F)cc3)n2N=C1c1cccs1 | Z90278248 | 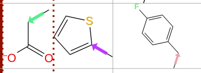 | | | 4.09 | 4.27 | 0.96 | 388.4 | 1.940 | 80.0 | 5 | 2 | 5 | 0.12 |
| O=C([O-])C[C@@H]1Sc2nnc(C3CC3)n2N=C1c1ccc(F)cc1 | Z90274599 | 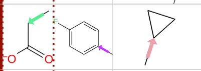 | | | 4.04 | 4.38 | 0.92 | 332.4 | 0.972 | 80.0 | 5 | 2 | 4 | 0.27 |
| CCN(CC)S(=O)(=O)c1cccc(-c2nnc3n2N=C(c2ccc(C)cc2)[C@@H](CC(=O)[O-])S3)c1 | Z90274338 | 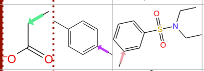 | | | 4.03 | 3.54 | **1.14** | 499.6 | 2.067 | 117.4 | 7 | 2 | 7 | 0.26 |
| O=C([O-])C[C@@H]1Sc2nnc(-c3ccco3)n2N=C1c1ccc(F)cc1 | Z90276391 | 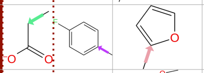 | | | 4.03 | 4.21 | 0.96 | 358.3 | 1.154 | 93.2 | 5 | 2 | 4 | 0.06 |
| COc1ccc(-c2nnc3n2N=C(c2cccs2)[C@H](CC(=O)[O-])S3)cc1 | Z90275560 | 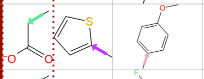 | | | 3.98 | 4.32 | 0.92 | 386.4 | 1.707 | 89.3 | 6 | 2 | 5 | 0.12 |
| O=C([O-])C[C@H]1Sc2nnc(Cc3ccc(F)cc3)n2N=C1c1ccccc1 | Z90278245 | 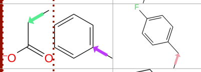 | | | 3.97 | 4.24 | 0.94 | 382.4 | 2.027 | 80.0 | 5 | 2 | 5 | 0.11 |
| COc1ccccc1-c1nnc2n1N=C(c1ccc(Cl)cc1)[C@H](CC(=O)[O-])S2 | Z90275299 | 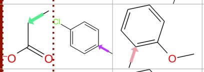 | | | 3.91 | 4.14 | 0.94 | 414.9 | 2.160 | 89.3 | 6 | 2 | 5 | 0.11 |
| COc1ccccc1-c1nnc2n1N=C(c1ccccc1)[C@H](CC(=O)[O-])S2 | Z90275301 | 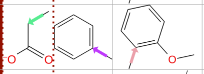 | | | 3.81 | 4.31 | 0.88 | 380.4 | 1.633 | 89.3 | 6 | 2 | 5 | 0.11 |
| COc1cccc(OCc2nnc3n2N=C(c2ccccc2)[C@H](CC(=O)[O-])S3)c1 | Z90278437 | 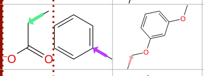 | | | 3.77 | 3.76 | **1.00** | 410.4 | 1.409 | 98.5 | 7 | 2 | 7 | 0.15 |
| Cc1ccc(C2=Nn3c(nnc3C3CC3)S[C@H]2CC(=O)[O-])cc1 | Z90274594 | 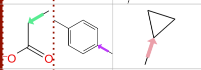 | | | 3.56 | 4.18 | 0.85 | 328.4 | 1.343 | 80.0 | 5 | 2 | 4 | 0.31 |
| O=C([O-])C[C@@H]1Sc2nnc(C3CC3)n2N=C1c1ccccc1 | Z90274597 | 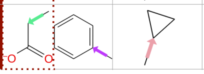 | | | 3.52 | 3.82 | 0.92 | 314.4 | 0.829 | 80.0 | 5 | 2 | 4 | 0.27 |

**Supplementary Figure S6. Activity and computed physicochemical properties of triazolothiadiazine derivatives bearing a carboxylic acid R3-group.** This panel displays a collection of active triazolothiadiazine compounds that share a carboxylic acid functional group at the R3 position. Each compound is shown with its SMILES string and annotated with its Enamine identifier, R-group substitutions, and experimentally measured RGS14 and RGS12 $pIC_{50}$ values (absolute $log_{10}$ of the compound concentration yielding 50% inhibition in the Transcreener® GDP-detection RGScreen™ assay). The selectivity ratio (RGS14 $pIC_{50}$ / RGS12 $pIC_{50}$) is included, with values >1.0 (bold) indicating preferential inhibition of RGS14. Additional physicochemical descriptors were computed using RDKit Chem Descriptors: molecular weight (MW, in Daltons), predicted octanol/water partition coefficient (cLogP), total polar surface area (TPSA, Å²), number of hydrogen bond acceptors (HBA) and donors (HBD), rotatable bond count, and the fraction of $sp^3$-hybridized carbon atoms ($Fsp^3$). This dataset highlights both the structural diversity and selective activity retained within the subset of acidic analogs.

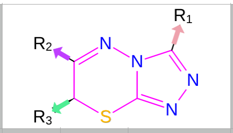

| SMILES | Enamine Name | R3 | R2 | R1 | RGS14 pIC50 | RGS12 pIC50 | Ratio | MW (Da) | cLogP (o/w) | TPSA (Å²) | accept H-bond | donor H-bond | Rotatable bond count | Fsp³ |
|---|---|---|---|---|---|---|---|---|---|---|---|---|---|---|
| COc1ccc(OCc2nnc3n2N=C(c2ccc(OC)c(OC)c2)CS3)cc1 | Z55660043 | H | 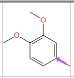 | 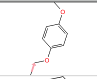 | **5.64** | 5.92 | 0.95 | 412.5 | 1.586 | 79.7 | 7 | 1 | 7 | 0.20 |
| Cc1ccc(C2=Nn3c(COc4ccccc4)nnc3SC2)c(C)c1 | Z55659515 | H | 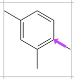 | 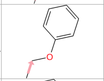 | 5.39 | 5.39 | **1.00** | 350.4 | 3.086 | 52.0 | 4 | 1 | 4 | 0.16 |
| c1ccc2c(c1)OC[C@@H](c1nnc3n1N=C(c1cc4ccccc4o1)CS3)O2 | Z55628018 | H | 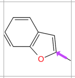 | 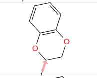 | 4.84 | 5.24 | 0.92 | 390.4 | 2.021 | 74.3 | 5 | 1 | 2 | 0.10 |
| CCCc1ccc(C2=Nn3c(COc4ccccc4F)nnc3SC2)cc1 | Z55660462 | H | 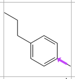 | 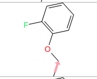 | 4.78 | 4.98 | 0.96 | 382.5 | 3.605 | 52.0 | 4 | 1 | 6 | 0.20 |
| COc1ccccc1C1=Nn2c(COc3ccccc3)nnc2SC1 | Z56944652 | H | 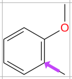 | 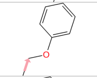 | 4.72 | 5.47 | 0.86 | 352.4 | 1.902 | 61.2 | 5 | 1 | 5 | 0.11 |
| c1ccc2c(c1)OC[C@@H](c1nnc3n1N=C(c1ccc4c(c1)OCCO4)CS3)O2 | Z55660304 | H | 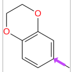 | 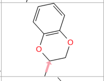 | 4.54 | 5.06 | 0.90 | 408.4 | 1.455 | 79.7 | 7 | 1 | 2 | 0.20 |
| COc1ccc(OCc2nnc3n2N=C(c2cc4ccccc4o2)CS3)cc1 | Z55627844 | H | 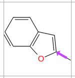 | 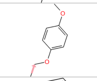 | 5.44 | 4.66 | 1.17 | 392.4 | 1.981 | 74.3 | 5 | 1 | 5 | 0.10 |
| FC(F)Oc1ccc(C2=Nn3c(COc4ccccc4)nnc3SC2)cc1 | Z56840012 | H | 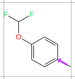 | 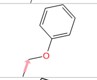 | 4.44 | 4.94 | 0.90 | 388.4 | 2.828 | 61.2 | 5 | 1 | 6 | 0.11 |
| CC(C)(C)c1ccc(C2=Nn3c(nnc3-c3ccco3)SC2)cc1 | Z55660110 | H | 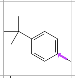 | 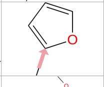 | 4.25 | 4.27 | **1.00** | 338.4 | 3.049 | 55.9 | 3 | 1 | 3 | 0.22 |
| COc1ccc(OCc2nnc3n2N=C(c2ccc(OC)cc2OC)CS3)cc1 | Z103673844 | H | 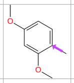 | 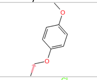 | 4.09 | 4.35 | 0.94 | 412.5 | 1.586 | 79.7 | 7 | 1 | 7 | 0.20 |
| Cc1ccc(C2=Nn3c(COc4ccc(Cl)cc4)nnc3SC2)c(C)c1 | Z55660524 | H | 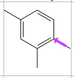 | 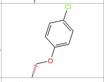 | 3.77 | 4.57 | 0.82 | 384.9 | 3.690 | 52.0 | 4 | 1 | 4 | 0.16 |
| CC(C)c1nnc2n1N=C(c1cc3ccccc3o1)CS2 | Z55627782 | H | 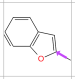 | 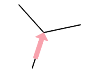 | 3.72 | 4.56 | 0.82 | 298.4 | 1.865 | 55.9 | 3 | 1 | 2 | 0.20 |
| COc1cccc(C2=Nn3c(COc4ccc(Cl)cc4)nnc3SC2)c1 | Z55660496 | H | 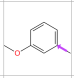 | 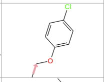 | 3.72 | 4.37 | 0.85 | 386.9 | 2.506 | 61.2 | 5 | 1 | 5 | 0.11 |
| COc1ccc(OCc2nnc3n2N=C(c2cc(C)n(-c4nccs4)c2C)CS3)cc1 | Z667824954 | H | 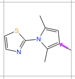 | 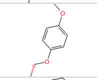 | 3.66 | 4.19 | 0.87 | 452.5 | 2.605 | 79.0 | 6 | 1 | 6 | 0.19 |
| Fc1ccccc1OCc1nnc2n1N=C(c1ccc(Cl)cc1)CS2 | Z55660438 | H | 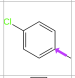 | 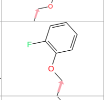 | 3.63 | 4.69 | 0.77 | 374.8 | 2.806 | 52.0 | 4 | 1 | 4 | 0.06 |
| COc1ccc(OCc2nnc3n2N=C(c2cccc(N4CCCC4=O)c2)CS3)cc1 | Z667824974 | H | 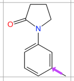 | 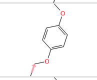 | 3.19 | 3.65 | 0.87 | 435.5 | 1.230 | 81.5 | 6 | 1 | 6 | 0.23 |
| COc1ccc(OCc2nnc3n2N=C(c2ccc(NC(C)=O)cc2)CS3)cc1 | Z55660046 | H | 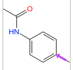 | 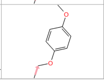 | < 3.00 | 3.81 | | 409.5 | 1.139 | 90.3 | 6 | 2 | 6 | 0.15 |

**Supplementary Figure S7. Activity and computed physicochemical properties of triazolothiadiazine derivatives lacking an R3-group.** This panel displays a collection of active triazolothiadiazine compounds that lack a functional group at the R3 position. Each compound is shown with its SMILES string and annotated with its Enamine identifier, R-group substitutions, and experimentally measured RGS14 and RGS12 $pIC_{50}$ values (absolute $\log_{10}$ of the compound concentration yielding 50% inhibition in the Transcreener® GDP-detection RGScreen™ assay). The selectivity ratio (RGS14 $pIC_{50}$ / RGS12 $pIC_{50}$) is included, with values equal to 1.0 (bold) indicating no preferential inhibition of RGS14 or RGS12. Additional physicochemical descriptors were computed using RDKit Chem Descriptors: molecular weight (MW, in Daltons), predicted octanol/water partition coefficient (cLogP), total polar surface area (TPSA, Å²), number of hydrogen bond acceptors (HBA) and donors (HBD), rotatable bond count, and the fraction of $sp^3$-hybridized carbon atoms ($Fsp^3$).

| | | 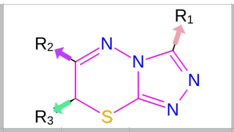 | | | | | | | | | | | | |
|---|---|---|---|---|---|---|---|---|---|---|---|---|---|---|
| **SMILES** | **Enamine Name** | **R3** | **R2** | **R1** | **RGS14 pIC50** | **RGS12 pIC50** | **Ratio** | **MW (Da)** | **cLogP (o/w)** | **TPSA (Å²)** | **accept H-bond** | **donor H-bond** | **Rotatable bond count** | **Fsp³** |
| **COC(=O)[C@@H]1Sc2nnc(COc3ccc(OC)cc3)n2N=C1c1cccc(N2CCCC2=O)c1** | Z7976707565 | 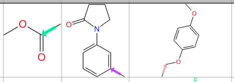 | | | 4.56 | 4.00 | **1.14** | 493.5 | 1.863 | 108.1 | 7 | 0 | 8 | 0.29 |
| **COC(=O)[C@H]1Sc2nnc(COc3ccc(F)cc3)n2N=C1c1cc2ccccc2o1** | Z8201263931 | 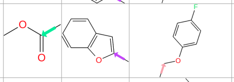 | | | 4.00 | 4.39 | 0.91 | 438.4 | 2.235 | 91.4 | 5 | 1 | 6 | 0.10 |
| **COc1ccc(OCc2nnc3n2N=C(c2ccc(C)cc2)[C@@H](C)S3)cc1** | Z55660052 | 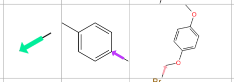 | | | 4.69 | 5.44 | 0.86 | 380.5 | 2.564 | 61.2 | 5 | 1 | 5 | 0.20 |
| **C[C@H]1Sc2nnc(Cc3ccc(Br)cc3)n2N=C1c1ccc(Cl)cc1** | Z55660592 | 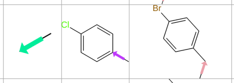 | | | 4.67 | 5.00 | 0.93 | 433.8 | 3.899 | 42.7 | 3 | 1 | 3 | 0.11 |
| **COc1ccc(-c2nnc3n2N=C(c2ccc(NC(C)=O)cc2)[C@@H](C)S3)cc1OC** | Z90277200 | 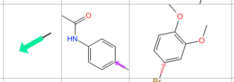 | | | 4.52 | 4.73 | 0.96 | 423.5 | 1.516 | 90.3 | 6 | 2 | 5 | 0.19 |
| **C[C@H]1Sc2nnc(Cc3ccc(Br)cc3)n2N=C1c1ccc([N-]S(C)(=O)=O)cc1** | Z90276813 | 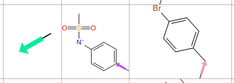 | | | 4.14 | 4.87 | 0.85 | 492.4 | 1.668 | 88.9 | 5 | 2 | 4 | 0.16 |
| **COc1ccc(Cc2nnc3n2N=C(c2ccc(C)c(C)c2)[C@@H](C)S3)cc1OC** | Z55660805 | 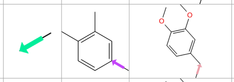 | | | 4.12 | 4.53 | 0.91 | 408.5 | 3.237 | 61.2 | 5 | 1 | 5 | 0.27 |
| **COc1ccc(Cc2nnc3n2N=C(c2ccccc2)[C@@H](C)S3)cc1OC** | Z55660788 | 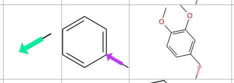 | | | 4.03 | 4.50 | 0.90 | 380.5 | 2.210 | 61.2 | 5 | 1 | 5 | 0.20 |
| **CCOc1ccccc1-c1nnc2n1N=C(c1ccc(NC(=O)C(C)C)cc1)[C@@H](C)S2** | Z90278674 | 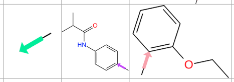 | | | 3.99 | 4.35 | 0.92 | 435.5 | 3.274 | 81.1 | 5 | 2 | 6 | 0.26 |
| **CSc1nnc2n1N=C(c1ccc(NC(=O)C(C)C)cc1)[C@H](C)S2** | Z90277842 | 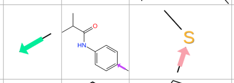 | | | 3.97 | 4.21 | 0.94 | 361.5 | 3.275 | 72.2 | 4 | 1 | 4 | 0.37 |
| **C1CC[C@H]2Sc3nnc([C@H]4CCCO4)n3N=C2C1** | Z90278035 | 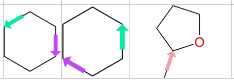 | | | 4.21 | 4.52 | 0.93 | 264.3 | 1.382 | 52.3 | 4 | 0 | 1 | 0.75 |

**Supplementary Figure S8. Activity and computed physicochemical properties of triazolothiadiazine derivatives with R3-groups different than a carboxylic-acid moiety.** This panel displays a collection of active triazolothiadiazine compounds that possess various different functional groups at the R3 position. Each compound is shown with its SMILES string and annotated with its Enamine identifier, R-group substitutions, and experimentally measured RGS14 and RGS12 $pIC_{50}$ values (absolute $\log_{10}$ of the compound concentration yielding 50% inhibition in the Transcreener® GDP-detection RGScreen™ assay). The selectivity ratio (RGS14 $pIC_{50}$ / RGS12 $pIC_{50}$) is included, with values >1.0 (bold) indicating preferential inhibition of RGS14. Additional physicochemical descriptors were computed using RDKit Chem Descriptors: molecular weight (MW, in Daltons), predicted octanol/water partition coefficient (cLogP), total polar surface area (TPSA, Å²), number of hydrogen bond acceptors (HBA) and donors (HBD), rotatable bond count, and the fraction of sp³-hybridized carbon atoms (Fsp³).

**Supplementary Table S2. List of 34 triazolothiadiazines tested for RGS-box inhibitory activity and found inactive *in vitro*.**

| SMILES | Name | Vina affinity (kcal/mol)* | logS @ pH 7.4§ | IUPAC name |
|---|---|---|---|---|
| Cc1ccc(C2=Nn3c(nnc3[C@@H]3COc4ccccc4O3)S[C@H]2C)cc1C | Z55660317 | -8.81 | -7.36 | (7S)-3-[(2R)-2,3-dihydro-1,4-benzodioxin-2-yl]-6-(3,4-dimethylphenyl)-7-methyl-7H-[1,2,4]triazolo[3,4-b][1,3,4]thiadiazine |
| COc1cccc(OCc2nnc3n2N=C(c2ccccc2)[C@@H](C(=O)Nc2ccc4c(c2)OCO4)S3)c1 | Z90278454 | -8.27 | -6.65 | N-(2H-1,3-benzodioxol-5-yl)-3-[(3-methoxyphenoxy)methyl]-6-phenyl-5H-[1,2,4]triazolo[3,4-b][1,3,4]thiadiazine-7-carboxamide |
| CCc1cccc2c(C3=Nn4c(COc5ccc(OC)cc5)nnc4S[C@H]3c3ccccc3)c[nH]c12 | Z90276677 | -8.23 | -7.67 | 7-ethyl-3-{3-[(4-methoxyphenoxy)methyl]-7-phenyl-5H-[1,2,4]triazolo[3,4-b][1,3,4]thiadiazin-6-yl}-1H-indole |
| Cc1ccc(C2=Nn3c(nnc3[C@H]3COc4ccccc4O3)S[C@H]2CC(=O)[O-])cc1 | Z90275810 | -8.20 | -1.51 | 2-{3-[(2S)-2,3-dihydro-1,4-benzodioxin-2-yl]-6-(4-methylphenyl)-5H-[1,2,4]triazolo[3,4-b][1,3,4]thiadiazin-7-yl}acetic acid |
| Fc1ccc(OCc2nnc3n2N=C(c2cc4ccccc4o2)CS3)cc1 | Z141354104 | -8.13 | -6.12 | 6-(1-benzofuran-2-yl)-3-[(4-fluorophenoxy)methyl]-5H-[1,2,4]triazolo[3,4-b][1,3,4]thiadiazine |
| COc1ccc(OCc2nnc3n2N=C(c2ccc4c(c2)NC(=O)CO4)CS3)cc1 | Z55627846 | -8.12 | -4.86 | 6-{3-[(4-methoxyphenoxy)methyl]-5H-[1,2,4]triazolo[3,4-b][1,3,4]thiadiazin-6-yl}-3,4-dihydro-2H-1,4-benzoxazin-3-one |
| Cc1ccc(C2=Nn3c(COc4c(Cl)cccc4Cl)nnc3S[C@@H]2C)cc1C | Z141361328 | -8.03 | -8.24 | (7R)-3-[(2,6-dichlorophenoxy)methyl]-6-(3,4-dimethylphenyl)-7-methyl-7H-[1,2,4]triazolo[3,4-b][1,3,4]thiadiazine |
| c1ccc(C2=Nn3c(Cc4cccc5ccccc45)nnc3SC2)cc1 | Z55659738 | -7.97 | -6.38 | 3-[(naphthalen-1-yl)methyl]-6-phenyl-5H-[1,2,4]triazolo[3,4-b][1,3,4]thiadiazine |
| COc1ccc(C2=Nn3c(nnc3[C@@H]3COc4ccccc4O3)SC2)cc1OC | Z55660314 | -7.95 | -5.21 | 3-[(2R)-2,3-dihydro-1,4-benzodioxin-2-yl]-6-(3,4-dimethoxyphenyl)-5H-[1,2,4]triazolo[3,4-b][1,3,4]thiadiazine |
| COc1ccc(Cc2nnc3n2N=C(c2ccc(Cl)cc2)[C@@H](C)S3)cc1OC | Z55660789 | -7.93 | -5.13 | 6-(4-chlorophenyl)-3-[(3,4-dimethoxyphenyl)methyl]-7-methyl-5H-[1,2,4]triazolo[3,4-b][1,3,4]thiadiazine |
| COc1cccc(OCc2nnc3n2N=C(c2c(C)[nH]c4ccccc24)[C@H](C)S3)c1 | Z90278458 | -7.82 | -6.75 | 3-[(7S)-3-[(3-methoxyphenoxy)methyl]-7-methyl-7H-[1,2,4]triazolo[3,4-b][1,3,4]thiadiazin-6-yl]-2-methyl-1H-indole |
| COc1ccccc1C1=Nn2c(Cc3cccc4ccccc34)nnc2SC1 | Z55659743 | -7.72 | -6.33 | 6-(2-methoxyphenyl)-3-[(naphthalen-1-yl)methyl]-5H-[1,2,4]triazolo[3,4-b][1,3,4]thiadiazine |
| COc1ccc(OCc2nnc3n2N=C(c2ccc4c(c2)CCO4)CS3)cc1 | Z667825018 | -7.70 | -5.04 | 6-(2,3-dihydro-1-benzofuran-5-yl)-3-[(4-methoxyphenoxy)methyl]-5H-[1,2,4]triazolo[3,4-b][1,3,4]thiadiazine |
| COC(=O)[C]1Sc2nnc(COc3ccc(OC)cc3)n2N=C1c1ccc2c(c1)CCO2 | Z8040061307 | -7.64 | -5.57 | 6-(2,3-dihydro-1-benzofuran-5-yl)-7-(methoxycarbonyl)-3-[(4-methoxyphenoxy)methyl]-7H-[1,2,4]triazolo[3,4-b][1,3,4]thiadiazin-7-yl |
| Cc1ccc(C2=Nn3c(COc4ccc(F)cc4)nnc3S[C@@H]2CC(=O)[O-])cc1 | Z141354596 | -7.56 | -1.37 | 2-{3-[(4-fluorophenoxy)methyl]-6-(4-methylphenyl)-5H-[1,2,4]triazolo[3,4-b][1,3,4]thiadiazin-7-yl}acetic acid |
| CCOc1ccccc1OCc1nnc2n1N=C(c1ccc(C)c(C)c1)[C@@H](C)S2 | Z141358858 | -7.55 | -7.16 | (7R)-6-(3,4-dimethylphenyl)-3-[(2-ethoxyphenoxy)methyl]-7-methyl-7H-[1,2,4]triazolo[3,4-b][1,3,4]thiadiazine |
| COc1ccc(OCc2nnc3n2N=C(c2ccc(F)cc2)[C@@H](CC(=O)[O-])S3)cc1 | Z90276647 | -7.49 | -1.29 | 2-[6-(4-fluorophenyl)-3-[(4-methoxyphenoxy)methyl]-5H-[1,2,4]triazolo[3,4-b][1,3,4]thiadiazin-7-yl]acetic acid |
| COc1ccc(OCc2nnc3n2N=C(c2cc(C)[nH]c2C)CS3)cc1 | Z667825046 | -7.49 | -4.26 | 3-{3-[(4-methoxyphenoxy)methyl]-5H-[1,2,4]triazolo[3,4-b][1,3,4]thiadiazin-6-yl}-2,5-dimethyl-1H-pyrrole |
| COc1ccc(OCc2nnc3n2N=C(c2ccc(NC(=O)CC(C)C)cc2)CS3)cc1 | Z55660047 | -7.47 | -6.02 | N-(4-{3-[(4-methoxyphenoxy)methyl]-5H-[1,2,4]triazolo[3,4-b][1,3,4]thiadiazin-6-yl}phenyl)-3-methylbutanamide |
| CCOC(=O)[C@H]1Sc2nnc(COc3ccc(OC)cc3)n2N=C1c1ccc(NC(C)=O)cc1 | Z8166491807 | -7.46 | -5.81 | ethyl (7S)-6-(4-acetamidophenyl)-3-[(4-methoxyphenoxy)methyl]-7H-[1,2,4]triazolo[3,4-b][1,3,4]thiadiazine-7-carboxylate |
| CCOC(=O)[C-]1Sc2nnc(COc3ccc(OC)cc3)n2N=C1c1ccc([N+](=O)[O-])cc1 | Z8186118910 | -7.37 | -5.63 | ethyl 3-[(4-methoxyphenoxy)methyl]-6-(4-nitrophenyl)-5H-[1,2,4]triazolo[3,4-b][1,3,4]thiadiazine-7-carboxylate |
| CCOC(=O)[C@H]1Sc2nnc(COc3ccc(OC)cc3)n2N=C1c1ccc(NC(=O)CC(C)C)cc1 | Z8024365455 | -7.37 | -6.97 | ethyl (7S)-3-[(4-methoxyphenoxy)methyl]-6-[4-(3-methylbutanamido)phenyl]-7H-[1,2,4]triazolo[3,4-b][1,3,4]thiadiazine-7-carboxylate |
| CCOC(=O)[C@H]1Sc2nnc(COc3ccc(OC)cc3)n2N=C1c1ccc(N)cc1 | Z7310720580 | -7.33 | -5.65 | ethyl (7S)-6-(4-aminophenyl)-3-[(4-methoxyphenoxy)methyl]-7H-[1,2,4]triazolo[3,4-b][1,3,4]thiadiazine-7-carboxylate |
| Cc1ccc(C2=Nn3c(nnc3-c3ccco3)S[C@H]2CC(=O)[O-])cc1 | Z90276386 | -7.23 | -1.73 | 2-[3-(furan-2-yl)-6-(4-methylphenyl)-5H-[1,2,4]triazolo[3,4-b][1,3,4]thiadiazin-7-yl]acetic acid |
| COC(=O)[C@H]1Sc2nnc(COc3ccc(OC)cc3)n2N=C1c1ccc(OC)cc1OC | Z8024373548 | -7.20 | -5.41 | methyl (7S)-6-(2,4-dimethoxyphenyl)-3-[(4-methoxyphenoxy)methyl]-7H-[1,2,4]triazolo[3,4-b][1,3,4]thiadiazine-7-carboxylate |
| CC(=O)Nc1ccc(C2=Nn3c(nnc3-c3cccnc3)SC2)cc1 | Z55659325 | -7.12 | -4.51 | N-{4-[3-(pyridin-3-yl)-5H-[1,2,4]triazolo[3,4-b][1,3,4]thiadiazin-6-yl]phenyl}acetamide |
| COC(=O)[C]1Sc2nnc(COc3ccc(OC)cc3)n2N=C1c1cccs1 | Z8024491322 | -7.10 | -5.36 | 7-(methoxycarbonyl)-3-[(4-methoxyphenoxy)methyl]-6-(thiophen-2-yl)-7H-[1,2,4]triazolo[3,4-b][1,3,4]thiadiazin-7-yl |
| Clc1ccc(C2=Nn3c(nnc3-c3cccnc3)SC2)cc1 | Z55659297 | -7.00 | -5.16 | 3-[6-(4-chlorophenyl)-5H-[1,2,4]triazolo[3,4-b][1,3,4]thiadiazin-3-yl]pyridine |
| COc1ccc(OCc2nnc3n2N=C(c2cccs2)CS3)cc1 | Z55627843 | -6.96 | -4.81 | 3-[(4-methoxyphenoxy)methyl]-6-(thiophen-2-yl)-5H-[1,2,4]triazolo[3,4-b][1,3,4]thiadiazine |
| COc1ccc(OCc2nnc3n2N=C(c2ccc[nH]2)CS3)cc1 | Z667824920 | -6.93 | -4.30 | 2-{3-[(4-methoxyphenoxy)methyl]-5H-[1,2,4]triazolo[3,4-b][1,3,4]thiadiazin-6-yl}-1H-pyrrole |
| CCc1nnc2n1N=C(c1ccc(C)cc1)[C@@H](CC(=O)[O-])S2 | Z90278306 | -6.65 | -0.16 | 2-[3-ethyl-6-(4-methylphenyl)-5H-[1,2,4]triazolo[3,4-b][1,3,4]thiadiazin-7-yl]acetic acid |
| COc1ccc(C2=Nn3c(nnc3-c3ccncc3)SC2)cc1OC | Z55659418 | -6.54 | -4.40 | 4-[6-(3,4-dimethoxyphenyl)-5H-[1,2,4]triazolo[3,4-b][1,3,4]thiadiazin-3-yl]pyridine |
| CCOc1ccccc1-c1nnc2n1N=C(c1ccc(OC)c(OC)c1)CS2 | Z90279873 | -6.52 | -5.84 | 6-(3,4-dimethoxyphenyl)-3-(2-ethoxyphenyl)-5H-[1,2,4]triazolo[3,4-b][1,3,4]thiadiazine |
| COc1ccccc1C1=Nn2c(nnc2-c2ccccc2OC)SC1 | Z55659692 | -6.41 | -5.58 | 3,6-bis(2-methoxyphenyl)-5H-[1,2,4]triazolo[3,4-b][1,3,4]thiadiazine |

* Best Vina-predicted binding affinity score (in kcal/mol) as derived by GNINA docking into pose 7 of RGS14 (PDB id 2JNU).

§ Predicted aqueous solubility at pH 7.4 (in $\log_{10}$[mol/L]) as derived by Chemaxon's chemicalize.com webserver (9, 10).

Column headers (as printed): Enamine Name; [Absorption/Human Oral Bioavailability 20%] Interpretation (Confidence); [Absorption/Human Oral Bioavailability 50%] Interpretation (Confidence); [Absorption/Human Intestinal Absorption] Interpretation (Confidence); [Absorption/P-Glycoprotein Inhibitor] Interpretation (Confidence); [Absorption/P-Glycoprotein Substrate] Interpretation (Confidence); [Distribution/Blood-Brain Barrier] Interpretation (Confidence); [Metabolism/CYP 1A2 Inhibitor] Interpretation (Confidence); [Metabolism/CYP 1A2_substrate] Interpretation (Confidence); [Metabolism/CYP 2C19 Inhibitor] Interpretation (Confidence); [Metabolism/CYP 2C19_substrate] Interpretation (Confidence); [Metabolism/CYP 2C9 Inhibitor] Interpretation (Confidence); [Metabolism/CYP 2C9 Substrate] Interpretation (Confidence); [Metabolism/CYP 2D6 Inhibitor] Interpretation (Confidence); [Metabolism/CYP 2D6 Substrate] Interpretation (Confidence); [Metabolism/CYP 3A4 Inhibitor] Interpretation (Confidence); [Metabolism/CYP 3A4 Substrate] Interpretation (Confidence); [Toxicity/AMES Mutagenesis] Interpretation (Confidence); [Toxicity/ Carcinogenesis] Interpretation (Confidence); [Toxicity/hERG Blockers] Interpretation (Confidence)

| Enamine Name | [Absorption/Human Oral Bioavailability 20%] Interpretation (Confidence) | [Absorption/Human Oral Bioavailability 50%] Interpretation (Confidence) | [Absorption/Human Intestinal Absorption] Interpretation (Confidence) | [Absorption/P-Glycoprotein Inhibitor] Interpretation (Confidence) | [Absorption/P-Glycoprotein Substrate] Interpretation (Confidence) | [Distribution/Blood-Brain Barrier] Interpretation (Confidence) | [Metabolism/CYP 1A2 Inhibitor] Interpretation (Confidence) | [Metabolism/CYP 1A2_substrate] Interpretation (Confidence) | [Metabolism/CYP 2C19 Inhibitor] Interpretation (Confidence) | [Metabolism/CYP 2C19_substrate] Interpretation (Confidence) | [Metabolism/CYP 2C9 Inhibitor] Interpretation (Confidence) | [Metabolism/CYP 2C9 Substrate] Interpretation (Confidence) | [Metabolism/CYP 2D6 Inhibitor] Interpretation (Confidence) | [Metabolism/CYP 2D6 Substrate] Interpretation (Confidence) | [Metabolism/CYP 3A4 Inhibitor] Interpretation (Confidence) | [Metabolism/CYP 3A4 Substrate] Interpretation (Confidence) | [Toxicity/AMES Mutagenesis] Interpretation (Confidence) | [Toxicity/ Carcinogenesis] Interpretation (Confidence) | [Toxicity/hERG Blockers] Interpretation (Confidence) |
|---|---|---|---|---|---|---|---|---|---|---|---|---|---|---|---|---|---|---|---|
| Z90276197 | Bioavailable (Medium) | Bioavailable (Medium) | Absorbed (High) | Non-Inhibitor (High) | Non-Substrate (High) | Penetrable (High) | Inhibitor (High) | Non-Substrate (High) | Non-Inhibitor (Medium) | Non-Substrate (Low) | Inhibitor (Medium) | Non-Substrate (Medium) | Non-Inhibitor (High) | Non-Substrate (Low) | Non-Inhibitor (Medium) | Non-Substrate (Medium) | Safe (High) | Safe (Medium) | Safe (High) |
| Z90275816 | Bioavailable (Low) | Bioavailable (Medium) | Absorbed (High) | Non-Inhibitor (High) | Non-Substrate (High) | Penetrable (High) | Inhibitor (High) | Non-Substrate (High) | Inhibitor (Low) | Non-Substrate (Low) | Inhibitor (Medium) | Non-Substrate (Medium) | Non-Inhibitor (High) | Non-Substrate (Medium) | Inhibitor (Medium) | Non-Substrate (High) | Toxic (Low) | Safe (High) | Safe (Low) |
| Z90277093 | Bioavailable (Low) | Bioavailable (Medium) | Absorbed (High) | Non-Inhibitor (High) | Non-Substrate (High) | Non-Penetrable (Medium) | Non-Inhibitor (High) | Non-Substrate (High) | Non-Inhibitor (High) | Non-Substrate (Medium) | Non-Inhibitor (Low) | Non-Substrate (High) | Non-Inhibitor (High) | Non-Substrate (High) | Non-Inhibitor (Medium) | Non-Substrate (High) | Safe (High) | Safe (Low) | Safe (High) |
| Z90276771 | Bioavailable (Medium) | Bioavailable (Medium) | Absorbed (High) | Non-Inhibitor (High) | Non-Substrate (High) | Penetrable (High) | Inhibitor (High) | Non-Substrate (High) | Non-Inhibitor (Medium) | Non-Substrate (Low) | Non-Inhibitor (Low) | Non-Substrate (Medium) | Non-Inhibitor (Medium) | Non-Substrate (Medium) | Non-Inhibitor (Medium) | Non-Substrate (Medium) | Safe (High) | Safe (Medium) | Safe (High) |
| Z90277285 | Bioavailable (Medium) | Bioavailable (Low) | Absorbed (High) | Non-Inhibitor (High) | Non-Substrate (High) | Penetrable (High) | Inhibitor (Medium) | Non-Substrate (High) | Non-Inhibitor (High) | Non-Substrate (Low) | Non-Inhibitor (Medium) | Non-Substrate (Low) | Non-Inhibitor (High) | Non-Substrate (Low) | Non-Inhibitor (Low) | Non-Substrate (High) | Safe (High) | Safe (High) | Safe (High) |
| Z90275304 | Bioavailable (Medium) | Bioavailable (Medium) | Absorbed (High) | Non-Inhibitor (High) | Non-Substrate (High) | Penetrable (High) | Non-Inhibitor (Low) | Non-Substrate (High) | Non-Inhibitor (Medium) | Non-Substrate (Medium) | Non-Inhibitor (Medium) | Non-Substrate (High) | Non-Inhibitor (High) | Non-Substrate (High) | Non-Inhibitor (High) | Non-Substrate (High) | Safe (Low) | Safe (Low) | Safe (Low) |
| Z90278051 | Bioavailable (Medium) | Bioavailable (Low) | Absorbed (High) | Non-Inhibitor (High) | Non-Substrate (High) | Penetrable (Medium) | Inhibitor (High) | Non-Substrate (Medium) | Non-Inhibitor (Low) | Non-Substrate (Low) | Non-Inhibitor (High) | Non-Substrate (High) | Non-Inhibitor (High) | Non-Substrate (Low) | Non-Inhibitor (High) | Non-Substrate (Low) | Safe (Medium) | Safe (High) | Safe (High) |
| Z90276194 | Bioavailable (Medium) | Bioavailable (Medium) | Absorbed (High) | Non-Inhibitor (High) | Non-Substrate (High) | Penetrable (Low) | Inhibitor (Medium) | Non-Substrate (High) | Inhibitor (Low) | Non-Substrate (Medium) | Inhibitor (Medium) | Non-Substrate (High) | Non-Inhibitor (High) | Non-Substrate (High) | Non-Inhibitor (High) | Substrate (Low) | Safe (High) | Safe (Low) | Safe (Medium) |
| Z90278242 | Bioavailable (Medium) | Bioavailable (Medium) | Absorbed (High) | Non-Inhibitor (High) | Non-Substrate (High) | Penetrable (High) | Non-Inhibitor (Low) | Non-Substrate (High) | Inhibitor (Low) | Non-Substrate (Medium) | Non-Inhibitor (Low) | Non-Substrate (High) | Non-Inhibitor (High) | Non-Substrate (Medium) | Non-Inhibitor (High) | Non-Substrate (Low) | Safe (High) | Toxic (Low) | Safe (High) |
| Z90276389 | Bioavailable (Medium) | Bioavailable (Low) | Absorbed (High) | Non-Inhibitor (High) | Non-Substrate (High) | Penetrable (High) | Inhibitor (High) | Non-Substrate (Medium) | Non-Inhibitor (Low) | Non-Substrate (Medium) | Non-Inhibitor (Medium) | Non-Substrate (High) | Non-Inhibitor (High) | Non-Substrate (Low) | Non-Inhibitor (High) | Non-Substrate (High) | Safe (High) | Toxic (Low) | Safe (High) |
| Z90278248 | Bioavailable (Medium) | Bioavailable (Medium) | Absorbed (High) | Non-Inhibitor (High) | Non-Substrate (High) | Penetrable (High) | Inhibitor (High) | Non-Substrate (High) | Non-Inhibitor (Medium) | Non-Substrate (Low) | Inhibitor (Low) | Non-Substrate (Low) | Inhibitor (High) | Non-Substrate (Medium) | Non-Inhibitor (Low) | Non-Substrate (High) | Safe (High) | Safe (Medium) | Safe (High) |
| Z90274599 | Bioavailable (Medium) | Bioavailable (Medium) | Absorbed (High) | Non-Inhibitor (High) | Non-Substrate (High) | Penetrable (High) | Inhibitor (High) | Non-Substrate (Medium) | Non-Inhibitor (Medium) | Non-Substrate (Low) | Non-Inhibitor (High) | Non-Substrate (Medium) | Non-Inhibitor (High) | Non-Substrate (Medium) | Non-Inhibitor (High) | Non-Substrate (High) | Safe (High) | Safe (High) | Safe (High) |
| Z90274338 | Bioavailable (Medium) | Bioavailable (Medium) | Absorbed (High) | Non-Inhibitor (High) | Non-Substrate (High) | Non-Penetrable (High) | Inhibitor (Low) | Non-Substrate (High) | Non-Inhibitor (Medium) | Non-Substrate (Medium) | Inhibitor (High) | Non-Substrate (High) | Non-Inhibitor (High) | Non-Substrate (High) | Non-Inhibitor (Medium) | Non-Substrate (High) | Safe (High) | Safe (Medium) | Toxic (High) |
| Z90276391 | Bioavailable (High) | Bioavailable (Low) | Absorbed (High) | Non-Inhibitor (High) | Non-Substrate (High) | Penetrable (High) | Inhibitor (High) | Non-Substrate (Medium) | Inhibitor (Low) | Non-Substrate (Medium) | Non-Inhibitor (Medium) | Non-Substrate (High) | Non-Inhibitor (High) | Non-Substrate (Medium) | Non-Inhibitor (High) | Non-Substrate (High) | Safe (High) | Safe (Low) | Safe (High) |
| Z90275560 | Bioavailable (Low) | Bioavailable (Low) | Absorbed (High) | Non-Inhibitor (High) | Non-Substrate (High) | Penetrable (Low) | Non-Inhibitor (Low) | Non-Substrate (High) | Non-Inhibitor (Medium) | Non-Substrate (Medium) | Non-Inhibitor (Medium) | Non-Substrate (High) | Non-Inhibitor (High) | Non-Substrate (High) | Non-Inhibitor (High) | Non-Substrate (High) | Safe (Low) | Safe (Low) | Safe (Medium) |
| Z90278245 | Bioavailable (Medium) | Bioavailable (Medium) | Absorbed (High) | Non-Inhibitor (High) | Non-Substrate (High) | Penetrable (High) | Inhibitor (High) | Non-Substrate (High) | Non-Inhibitor (Medium) | Non-Substrate (Low) | Non-Inhibitor (Low) | Non-Substrate (Medium) | Non-Inhibitor (High) | Non-Substrate (Medium) | Non-Inhibitor (Medium) | Non-Substrate (High) | Safe (High) | Safe (Medium) | Safe (High) |
| Z90275299 | Bioavailable (Medium) | Bioavailable (Medium) | Absorbed (High) | Non-Inhibitor (High) | Non-Substrate (High) | Penetrable (Medium) | Inhibitor (Low) | Non-Substrate (High) | Non-Inhibitor (Low) | Non-Substrate (High) | Non-Inhibitor (Low) | Non-Substrate (High) | Non-Inhibitor (High) | Non-Substrate (High) | Non-Inhibitor (High) | Non-Substrate (High) | Safe (High) | Safe (Low) | Safe (Low) |
| Z90275301 | Bioavailable (Medium) | Bioavailable (Low) | Absorbed (High) | Non-Inhibitor (High) | Non-Substrate (High) | Penetrable (Medium) | Non-Inhibitor (Low) | Non-Substrate (High) | Non-Inhibitor (Medium) | Non-Substrate (High) | Non-Inhibitor (Medium) | Non-Substrate (Medium) | Non-Inhibitor (High) | Non-Substrate (Medium) | Non-Inhibitor (High) | Non-Substrate (High) | Safe (High) | Safe (Low) | Safe (Medium) |
| Z90278437 | Bioavailable (Low) | Bioavailable (Medium) | Absorbed (High) | Non-Inhibitor (High) | Non-Substrate (High) | Penetrable (Low) | Inhibitor (Medium) | Non-Substrate (High) | Non-Inhibitor (High) | Non-Substrate (Medium) | Inhibitor (Low) | Non-Substrate (High) | Non-Inhibitor (High) | Non-Substrate (High) | Non-Inhibitor (High) | Non-Substrate (High) | Safe (High) | Safe (Low) | Safe (Medium) |
| Z90274594 | Bioavailable (Medium) | Bioavailable (Medium) | Absorbed (High) | Non-Inhibitor (High) | Non-Substrate (High) | Penetrable (Low) | Inhibitor (Low) | Non-Substrate (High) | Non-Inhibitor (High) | Non-Substrate (Medium) | Non-Inhibitor (Medium) | Non-Substrate (High) | Non-Inhibitor (High) | Non-Substrate (High) | Non-Inhibitor (High) | Non-Substrate (Medium) | Safe (Medium) | Safe (Low) | Safe (High) |
| Z90274597 | Bioavailable (Medium) | Bioavailable (Low) | Absorbed (High) | Non-Inhibitor (High) | Non-Substrate (High) | Penetrable (High) | Inhibitor (High) | Non-Substrate (Medium) | Non-Inhibitor (High) | Non-Substrate (Low) | Non-Inhibitor (High) | Non-Substrate (High) | Non-Inhibitor (High) | Non-Substrate (Low) | Non-Inhibitor (High) | Non-Substrate (High) | Safe (High) | Safe (Medium) | Safe (High) |
| Z55660043 | Bioavailable (Medium) | Bioavailable (Medium) | Absorbed (High) | Non-Inhibitor (High) | Non-Substrate (Medium) | Penetrable (High) | Inhibitor (High) | Substrate (Low) | Inhibitor (High) | Non-Substrate (Low) | Inhibitor (High) | Non-Substrate (Medium) | Non-Inhibitor (High) | Non-Substrate (Low) | Inhibitor (High) | Substrate (Low) | Safe (High) | Safe (High) | Safe (Medium) |
| Z55659515 | Bioavailable (Medium) | Bioavailable (Medium) | Absorbed (High) | Inhibitor (High) | Non-Substrate (Medium) | Penetrable (High) | Inhibitor (High) | Substrate (Low) | Inhibitor (High) | Substrate (Low) | Inhibitor (High) | Non-Substrate (High) | Non-Inhibitor (High) | Non-Substrate (Low) | Inhibitor (Medium) | Substrate (Medium) | Safe (High) | Safe (Low) | Safe (High) |
| Z55628018 | Bioavailable (Medium) | Bioavailable (Low) | Absorbed (High) | Inhibitor (High) | Non-Substrate (High) | Penetrable (High) | Inhibitor (High) | Substrate (Medium) | Inhibitor (High) | Non-Substrate (Low) | Inhibitor (High) | Non-Substrate (High) | Non-Inhibitor (High) | Non-Substrate (Low) | Inhibitor (Medium) | Non-Substrate (High) | Toxic (Medium) | Toxic (Low) | Toxic (Medium) |
| Z55660462 | Bioavailable (Medium) | Bioavailable (Medium) | Absorbed (High) | Inhibitor (High) | Non-Substrate (High) | Penetrable (High) | Inhibitor (High) | Non-Substrate (Low) | Inhibitor (High) | Non-Substrate (Low) | Inhibitor (High) | Non-Substrate (Low) | Non-Inhibitor (Medium) | Non-Substrate (Low) | Inhibitor (Medium) | Substrate (Medium) | Safe (High) | Safe (Low) | Safe (Medium) |
| Z56944652 | Bioavailable (Medium) | Bioavailable (Medium) | Absorbed (High) | Non-Inhibitor (High) | Non-Substrate (High) | Penetrable (High) | Inhibitor (High) | Substrate (Low) | Inhibitor (High) | Non-Substrate (Low) | Inhibitor (High) | Non-Substrate (High) | Non-Inhibitor (High) | Non-Substrate (Low) | Inhibitor (Medium) | Non-Substrate (Low) | Safe (Low) | Safe (Medium) | Safe (High) |
| Z55660304 | Bioavailable (Medium) | Bioavailable (Medium) | Absorbed (High) | Non-Inhibitor (Medium) | Non-Substrate (Medium) | Penetrable (High) | Inhibitor (High) | Substrate (Low) | Inhibitor (High) | Non-Substrate (Low) | Inhibitor (High) | Non-Substrate (Medium) | Non-Inhibitor (High) | Non-Substrate (Low) | Inhibitor (High) | Substrate (Medium) | Toxic (Medium) | Safe (Low) | Toxic (High) |
| Z55627844 | Bioavailable (Medium) | Bioavailable (Medium) | Absorbed (High) | Inhibitor (High) | Non-Substrate (High) | Penetrable (High) | Inhibitor (High) | Substrate (Medium) | Inhibitor (High) | Non-Substrate (Low) | Inhibitor (High) | Non-Substrate (High) | Non-Inhibitor (High) | Non-Substrate (Low) | Inhibitor (High) | Non-Substrate (High) | Safe (Low) | Safe (Low) | Safe (High) |
| Z56840012 | Bioavailable (High) | Bioavailable (Medium) | Absorbed (High) | Non-Inhibitor (High) | Non-Substrate (High) | Penetrable (High) | Inhibitor (High) | Substrate (Low) | Inhibitor (High) | Non-Substrate (Low) | Inhibitor (High) | Substrate (Medium) | Non-Inhibitor (High) | Non-Substrate (Low) | Non-Inhibitor (Low) | Substrate (Low) | Safe (High) | Safe (Medium) | Toxic (Medium) |
| Z55660110 | Bioavailable (Medium) | Bioavailable (Low) | Absorbed (High) | Inhibitor (High) | Non-Substrate (High) | Penetrable (High) | Inhibitor (High) | Substrate (Low) | Inhibitor (High) | Non-Substrate (Low) | Inhibitor (High) | Non-Substrate (High) | Non-Inhibitor (Medium) | Substrate (Low) | Non-Inhibitor (Medium) | Non-Substrate (High) | Safe (High) | Toxic (Low) | Safe (Low) |
| Z103673844 | Bioavailable (Medium) | Bioavailable (Medium) | Absorbed (High) | Non-Inhibitor (High) | Non-Substrate (Medium) | Penetrable (High) | Inhibitor (High) | Substrate (Low) | Inhibitor (High) | Non-Substrate (Low) | Inhibitor (High) | Non-Substrate (Medium) | Non-Inhibitor (High) | Non-Substrate (Low) | Inhibitor (High) | Substrate (Low) | Safe (High) | Safe (High) | Safe (Medium) |
| Z55660524 | Bioavailable (High) | Bioavailable (Medium) | Absorbed (High) | Inhibitor (High) | Non-Substrate (High) | Penetrable (High) | Inhibitor (High) | Substrate (Low) | Inhibitor (High) | Substrate (Low) | Inhibitor (High) | Non-Substrate (High) | Non-Inhibitor (High) | Substrate (Low) | Inhibitor (High) | Substrate (High) | Safe (High) | Safe (Low) | Safe (High) |
| Z55627782 | Bioavailable (Medium) | Bioavailable (Low) | Absorbed (High) | Inhibitor (Medium) | Non-Substrate (High) | Penetrable (High) | Inhibitor (High) | Substrate (Medium) | Inhibitor (High) | Non-Substrate (Low) | Inhibitor (High) | Non-Substrate (High) | Non-Inhibitor (High) | Non-Substrate (Low) | Non-Inhibitor (High) | Non-Substrate (High) | Safe (Low) | Toxic (Low) | Safe (High) |
| Z55660496 | Bioavailable (High) | Bioavailable (Medium) | Absorbed (High) | Non-Inhibitor (High) | Non-Substrate (High) | Penetrable (High) | Inhibitor (High) | Substrate (Low) | Inhibitor (High) | Non-Substrate (Low) | Inhibitor (High) | Non-Substrate (Medium) | Non-Inhibitor (Low) | Non-Substrate (Low) | Inhibitor (High) | Substrate (Low) | Safe (High) | Safe (Medium) | Safe (Medium) |
| Z667824954 | Bioavailable (Medium) | Bioavailable (Medium) | Absorbed (High) | Inhibitor (High) | Non-Substrate (Medium) | Penetrable (High) | Inhibitor (High) | Non-Substrate (Medium) | Inhibitor (High) | Non-Substrate (Low) | Inhibitor (High) | Non-Substrate (High) | Non-Inhibitor (High) | Non-Substrate (Low) | Inhibitor (High) | Substrate (Medium) | Safe (Medium) | Safe (Medium) | Toxic (Low) |
| Z55660438 | Bioavailable (High) | Bioavailable (High) | Absorbed (High) | Inhibitor (High) | Non-Substrate (High) | Penetrable (High) | Inhibitor (High) | Non-Substrate (Low) | Inhibitor (High) | Non-Substrate (Low) | Inhibitor (High) | Non-Substrate (Low) | Non-Inhibitor (High) | Non-Substrate (Low) | Inhibitor (Low) | Substrate (Medium) | Safe (High) | Safe (Low) | Safe (High) |
| Z667824974 | Bioavailable (Medium) | Bioavailable (Medium) | Absorbed (High) | Inhibitor (High) | Non-Substrate (Medium) | Penetrable (High) | Inhibitor (High) | Non-Substrate (Low) | Inhibitor (High) | Non-Substrate (Low) | Inhibitor (High) | Non-Substrate (High) | Non-Inhibitor (High) | Non-Substrate (Low) | Inhibitor (High) | Substrate (High) | Safe (High) | Safe (Medium) | Safe (Medium) |
| Z55660046 | Bioavailable (Medium) | Bioavailable (Medium) | Absorbed (High) | Inhibitor (High) | Non-Substrate (High) | Penetrable (High) | Inhibitor (High) | Non-Substrate (Medium) | Inhibitor (Medium) | Non-Substrate (Low) | Inhibitor (High) | Non-Substrate (Medium) | Non-Inhibitor (High) | Non-Substrate (Low) | Inhibitor (High) | Substrate (Medium) | Safe (High) | Safe (Medium) | Safe (Low) |
| Z7976707565 | Bioavailable (Low) | Bioavailable (Medium) | Absorbed (High) | Inhibitor (High) | Non-Substrate (Medium) | Non-Penetrable (High) | Inhibitor (High) | Non-Substrate (Medium) | Inhibitor (Medium) | Non-Substrate (Low) | Inhibitor (High) | Non-Substrate (High) | Non-Inhibitor (High) | Non-Substrate (Medium) | Inhibitor (High) | Substrate (High) | Safe (High) | Safe (High) | Safe (Medium) |
| Z8201263931 | Bioavailable (Medium) | Bioavailable (Medium) | Absorbed (High) | Inhibitor (High) | Non-Substrate (High) | Non-Penetrable (High) | Inhibitor (High) | Non-Substrate (Low) | Inhibitor (High) | Non-Substrate (Medium) | Inhibitor (High) | Non-Substrate (Medium) | Non-Inhibitor (High) | Non-Substrate (Medium) | Inhibitor (Medium) | Non-Substrate (High) | Safe (High) | Safe (High) | Toxic (High) |
| Z55660052 | Bioavailable (Medium) | Bioavailable (Medium) | Absorbed (High) | Inhibitor (Medium) | Non-Substrate (High) | Penetrable (High) | Inhibitor (High) | Non-Substrate (Low) | Inhibitor (High) | Non-Substrate (Low) | Inhibitor (High) | Non-Substrate (High) | Non-Inhibitor (High) | Non-Substrate (Low) | Inhibitor (High) | Substrate (Low) | Safe (High) | Safe (Medium) | Toxic (Medium) |
| Z55660592 | Bioavailable (High) | Bioavailable (High) | Absorbed (High) | Inhibitor (Low) | Non-Substrate (High) | Penetrable (High) | Inhibitor (High) | Substrate (Low) | Inhibitor (High) | Substrate (Low) | Inhibitor (High) | Non-Substrate (High) | Non-Inhibitor (High) | Non-Substrate (Low) | Inhibitor (Low) | Substrate (Medium) | Safe (High) | Safe (Medium) | Toxic (Low) |
| Z90277200 | Bioavailable (Medium) | Bioavailable (Medium) | Absorbed (High) | Inhibitor (High) | Non-Substrate (Medium) | Penetrable (High) | Non-Inhibitor (Low) | Non-Substrate (Low) | Inhibitor (High) | Non-Substrate (Low) | Inhibitor (High) | Non-Substrate (Low) | Non-Inhibitor (High) | Non-Substrate (Low) | Inhibitor (Medium) | Substrate (Medium) | Safe (High) | Safe (Medium) | Toxic (High) |
| Z90276813 | Bioavailable (Medium) | Bioavailable (Medium) | Absorbed (High) | Inhibitor (High) | Non-Substrate (High) | Penetrable (High) | Inhibitor (Medium) | Non-Substrate (Medium) | Inhibitor (High) | Non-Substrate (Low) | Inhibitor (High) | Non-Substrate (High) | Inhibitor (Medium) | Non-Substrate (Low) | Inhibitor (Medium) | Substrate (High) | Safe (High) | Safe (Medium) | Toxic (Medium) |
| Z55660805 | Bioavailable (Medium) | Bioavailable (Medium) | Absorbed (High) | Inhibitor (High) | Non-Substrate (Medium) | Penetrable (High) | Inhibitor (High) | Substrate (Low) | Inhibitor (High) | Substrate (Low) | Inhibitor (High) | Non-Substrate (Medium) | Non-Inhibitor (High) | Substrate (Low) | Inhibitor (High) | Substrate (High) | Safe (High) | Safe (Medium) | Safe (Medium) |
| Z55660788 | Bioavailable (Medium) | Bioavailable (Medium) | Absorbed (High) | Inhibitor (Medium) | Non-Substrate (Medium) | Penetrable (High) | Inhibitor (High) | Substrate (Low) | Inhibitor (High) | Non-Substrate (Low) | Inhibitor (High) | Non-Substrate (Medium) | Non-Inhibitor (High) | Substrate (Low) | Inhibitor (High) | Substrate (Low) | Safe (High) | Safe (Medium) | Safe (Low) |
| Z90278674 | Bioavailable (Medium) | Bioavailable (Low) | Absorbed (High) | Inhibitor (High) | Non-Substrate (Medium) | Penetrable (High) | Non-Inhibitor (Low) | Non-Substrate (Low) | Inhibitor (High) | Substrate (Low) | Inhibitor (High) | Non-Substrate (Medium) | Non-Inhibitor (High) | Non-Substrate (Low) | Inhibitor (High) | Substrate (Medium) | Safe (Medium) | Safe (Medium) | Toxic (Medium) |
| Z90277842 | Bioavailable (Medium) | Bioavailable (Medium) | Absorbed (High) | Non-Inhibitor (High) | Non-Substrate (High) | Penetrable (High) | Inhibitor (High) | Non-Substrate (Low) | Inhibitor (Medium) | Substrate (Low) | Inhibitor (High) | Non-Substrate (Medium) | Non-Inhibitor (High) | Non-Substrate (Low) | Inhibitor (Low) | Substrate (Low) | Safe (High) | Safe (Low) | Safe (High) |
| Z90278035 | Bioavailable (Medium) | Bioavailable (Medium) | Absorbed (High) | Non-Inhibitor (High) | Non-Substrate (High) | Penetrable (High) | Inhibitor (High) | Non-Substrate (Low) | Non-Inhibitor (High) | Substrate (Low) | Non-Inhibitor (High) | Substrate (Medium) | Non-Inhibitor (High) | Substrate (Low) | Non-Inhibitor (High) | Substrate (Low) | Safe (High) | Safe (High) | Safe (High) |

**Supplementary Table S3. Predicted pharmacokinetic and toxicity profiles of triazolothiadiazine analogs using the Deep-PK deep-learning platform.** Pharmacokinetic and toxicity properties of the triazolothiadiazine analogs were computationally predicted using the Deep-PK platform (11), a graph neural network-based tool trained on curated experimental datasets encompassing 73 ADMET-related endpoints. Each row lists a triazolothiadiazine compound previously characterized in GAP inhibition assays, with the compound name color-coded as in prior figures (e.g., red background denotes the presence of a carboxylic acid $R_3$-group). The table columns reflect Deep-PK predictions across key ADMET categories: absorption (including oral bioavailability and intestinal uptake), distribution (e.g., blood–brain barrier penetration), metabolism (inhibition and substrate status for major cytochrome P450 isoforms), and toxicity (mutagenicity, carcinogenicity, and hERG blockade). Each cell includes both a qualitative classification (e.g., "Bioavailable", "Safe", "Non-Inhibitor") and a parenthetical confidence score assigned by the model. Background shading of table cells provides interpretative cues: green cells denote "favorable" or "safe" predictions with medium to high confidence, whereas yellow cells indicate predictions that may warrant caution, reflecting borderline confidence levels, potentially concerning classifications (*e.g.*, weak CYP inhibition), or lower prediction certainty.

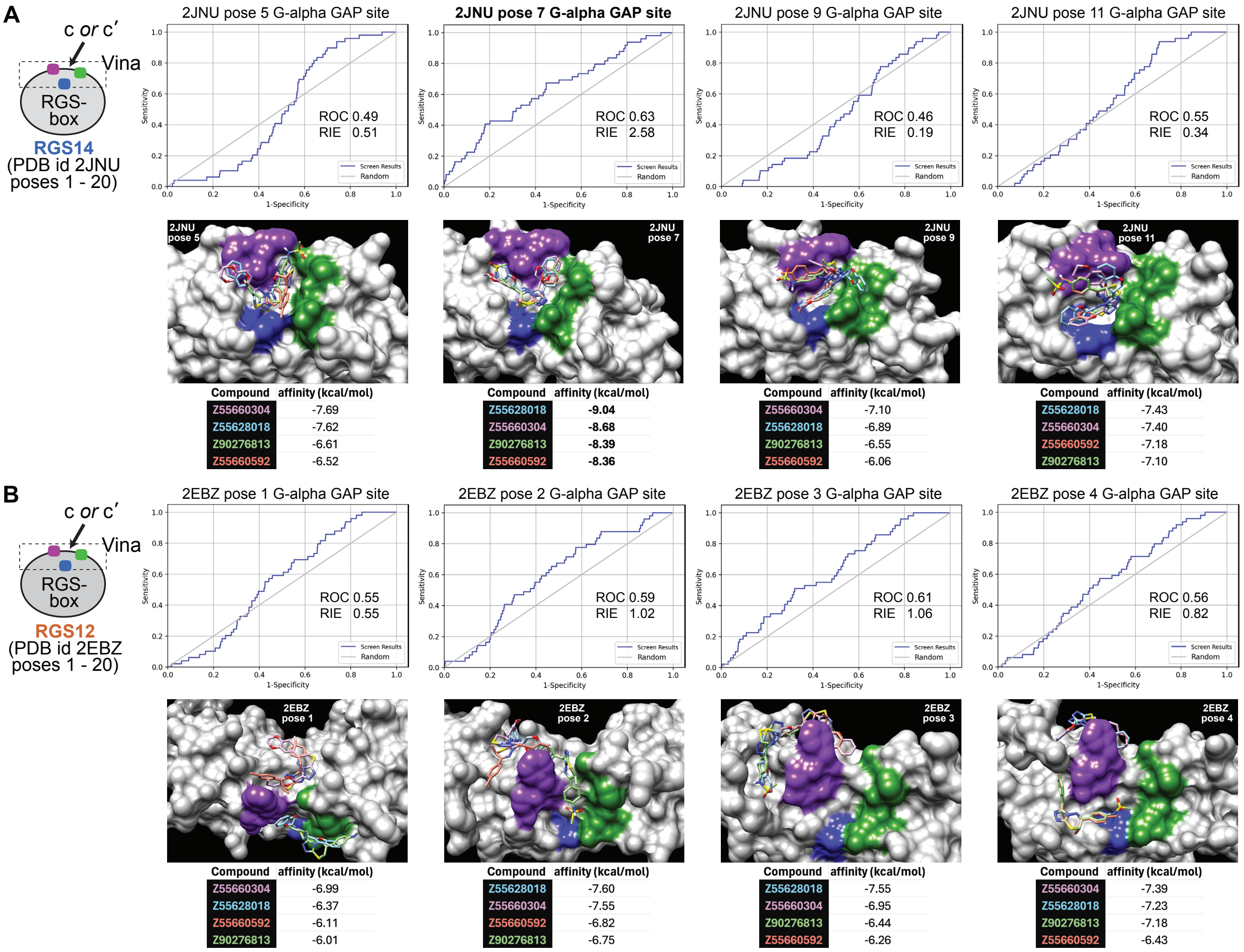

**Supplementary Figure S9. Enrichment of active triazolothiadiazine ligands into the Gα-binding site across multiple RGS-box conformers.** (**A-B**) Structure-based virtual screening results for selected conformers of the RGS14 (PDB id 2JNU; panel A) and RGS12 (PDB id 2EBZ; panel B) RGS-box domains, each assessed using AutoDock Vina (v1.2.5), for their ability to discriminate between active and decoy ligands. Ligand sets comprised either the 49 active triazolothiadiazine compounds (denoted as "compound" or "c") or 4,160 physicochemically matched decoy structures generated by DUD-E (denoted as "compound-prime" or c′). Each receptor conformer is labeled with its pose number and the location of the presumptive Gα-binding groove (color-coded as in Figure 2). ROC AUC and robust initial enrichment (RIE) values are shown for each pose, reflecting the degree to which active compounds were preferentially ranked within the top of the docking score distribution. Pose 7 of 2JNU yielded the highest enrichment (ROC AUC = 0.63, RIE = 2.58).

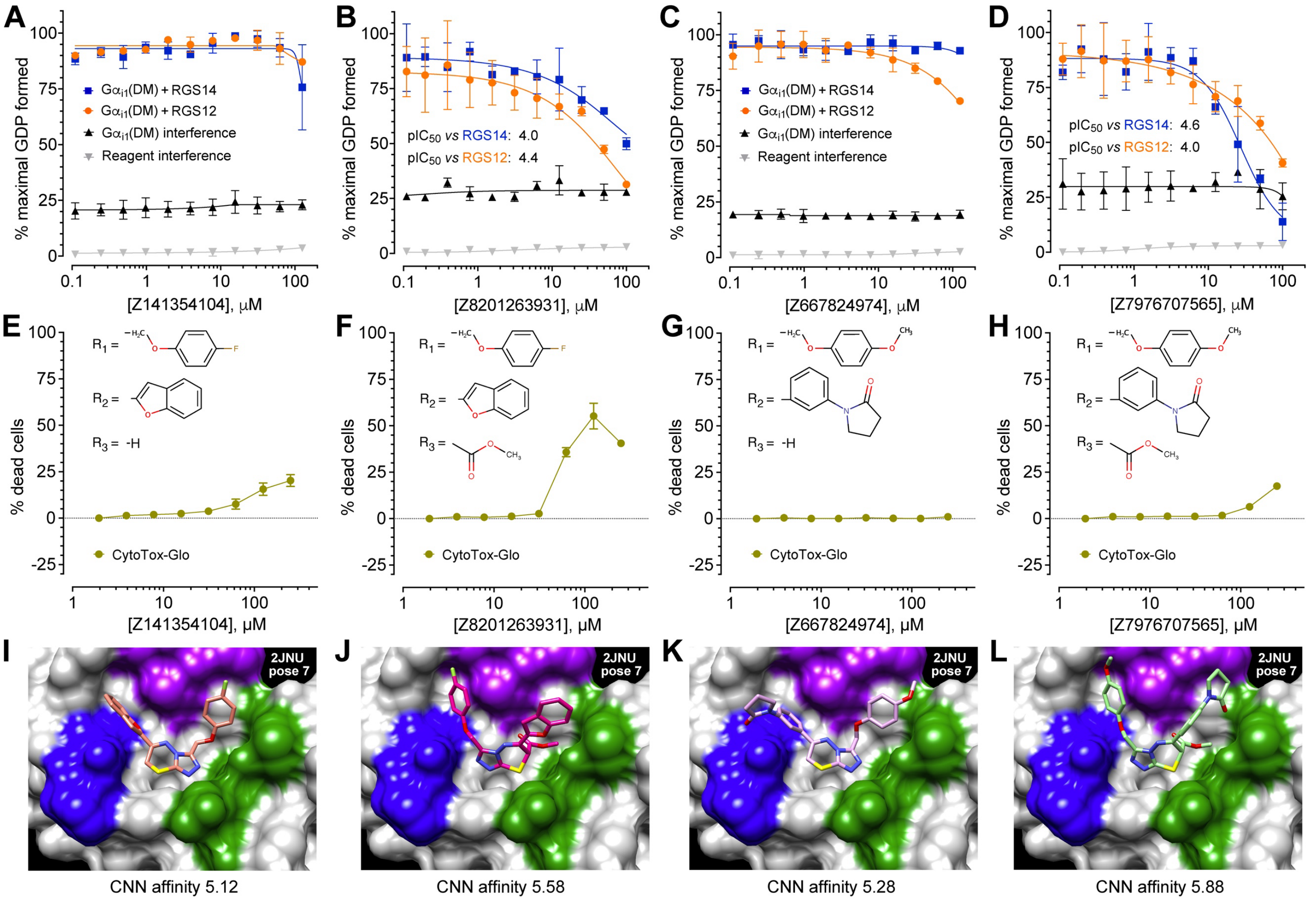

**Supplementary Figure S10. Inhibitors of the triazolothiadiazine chemotype increase in potency against RGS14 GAP activity, but also in cytotoxicity, upon provision of a methyl ester $R_3$-group.** (**A-D**) Transcreener® GDP-detection RGScreen™ assay measuring GDP production as a function of compound concentration for representative members of the active triazolothiadiazine series. GDP production was catalyzed by $G\alpha_{i1}$(DM) protein and accelerated by the indicated RGS-box protein. Additional controls demonstrate lack of signal interference by the tested compound alone (*i.e.*, on detection reagents) or intrinsic $G\alpha_{i1}$(DM) GTPase activity. (**E–H**) Cytotoxicity profiles for the same compounds in THP1-Dual™ cells using the CytoTox-Glo assay (Promega). Compounds were added in a 2-fold dilution series beginning at 250 µM and incubated overnight. Luminescent signal corresponding to dead-cell protease activity was measured first, followed by digitonin lysis and total-cell signal acquisition. Percent cytotoxicity was calculated per the manufacturer's instructions. (**I–L**) GNINA docking results for Z141354104 (panel I) and related congener Z8201263931 (panel J), and for Z667824974 (panel K) and related congener Z7976707565 (panel L), into pose 7 of the RGS14 RGS-box (PDB id 2JNU). Each pair represents a switch from lacking a formal R3-group (left) to a methyl ester (right), resulting in increased CNN affinity scores (*e.g.*, from 5.12 to 5.58 for panels I *vs* J) and a reciprocal "flip" in predicted binding orientation, whereby R1 and R2 substituents project into opposite faces of the shallow canyon at the presumptive Gα-binding interface. We are defining this predicted flipped orientation binding behavior as "ambidextrous" binding.

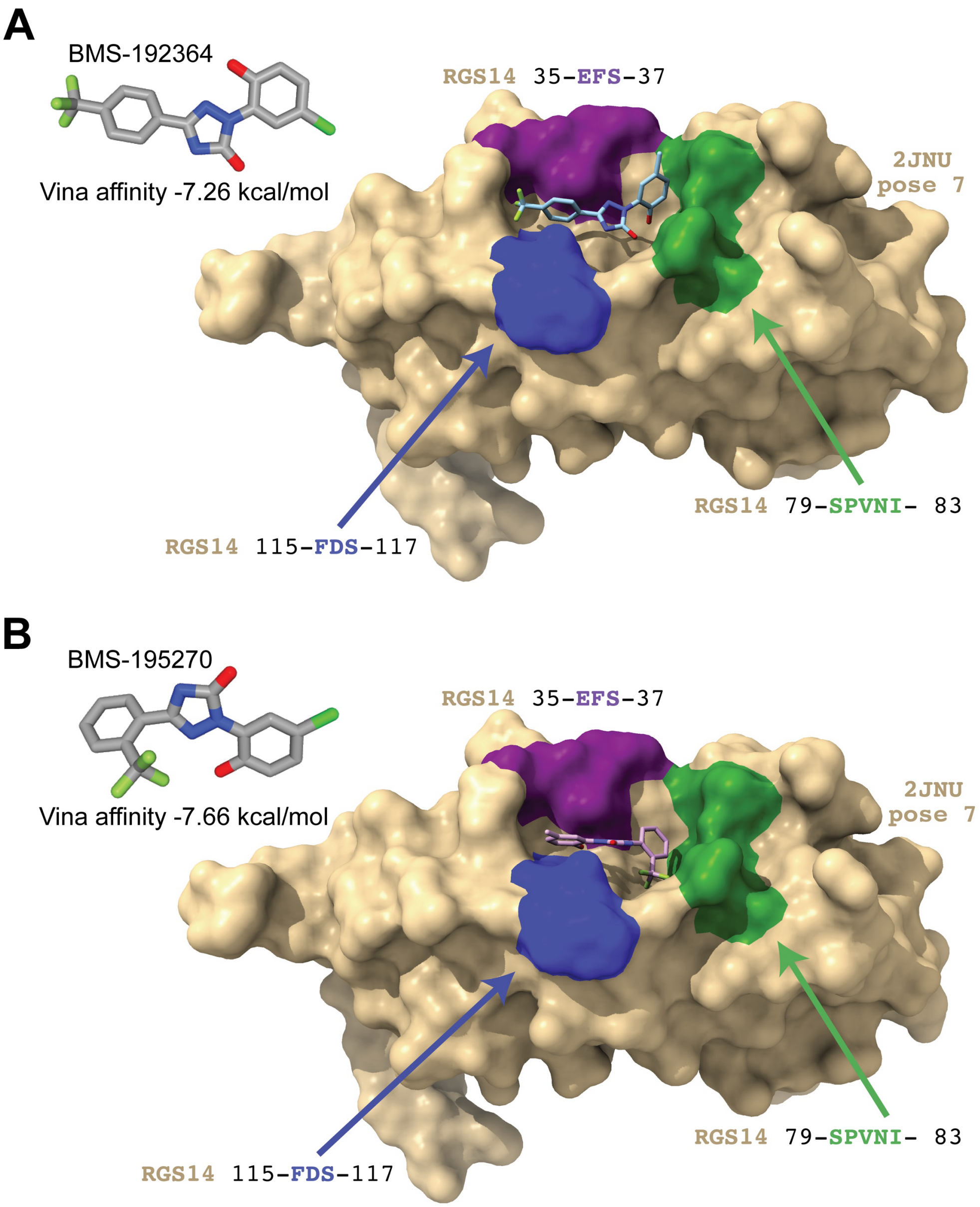

**Supplementary Figure S11. Comparison of predicted binding modes for original BMS diaryl-triazolone inhibitors docked into the presumptive Gα-binding groove of RGS14. (A)** Predicted pose of BMS-192364 (Vina affinity score: –7.26 kcal/mol) docked into the shallow, solvent-exposed canyon at the presumptive Gα-binding interface of pose 7 of the RGS14 RGS-box (PDB id 2JNU), as identified by GNINA-based global docking. **(B)** Predicted binding pose of BMS-195270 (Vina affinity score: –7.66 kcal/mol), a structurally related compound shown to be weakly active in vitro, similarly docked into the same receptor conformation. The spatial proximity of RGS14 residues E35, F36, S37, S79, P80, V81, N82, I83, F115, D116, and S117 (labels color-coded to match Figure 1 orientation markers) highlights the shallow nature of the Gα-interaction site and supports its predicted engagement by both ligands. Notably, these BMS compounds also exhibit flipped or "ambidextrous" orientations of substituent groups around their central core, consistent with the alternating R1/R2 projection pattern observed in the active triazolothiadiazine series.

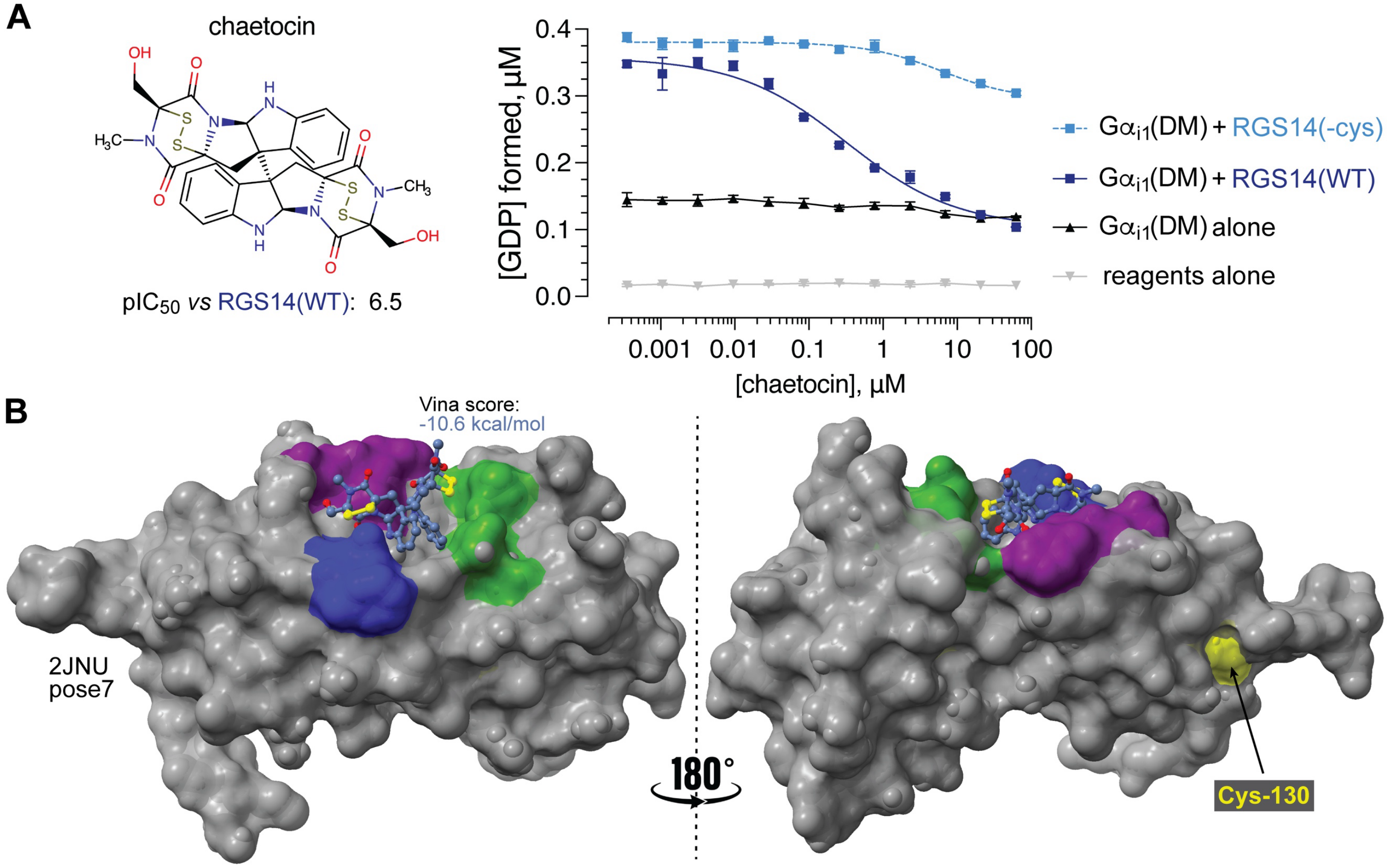

**Supplementary Figure S12. Chaetocin, an inhibitor of thioredoxin reductase and histone methyltransferases, also inhibits RGS14 RGS-box GAP activity *in vitro* but predominantly via cysteine reactivity. (A)** *Left,* structure of chaetocin, a 696.8 Da fungal mycotoxin identified in this study by GNINA-based *in silico* screening of the RGS14 RGS-box shallow canyon (specified by PDB id 2JNU pose 7) against MolPort's 235,298-element "Protein-Protein Interactions Library" of commercially available compounds >500 Da (acquired from molport.com on 01.03.2025). Right, Transcreener® GDP-detection RGScreen™ assay results indicating that chaetocin specifically inhibits RGS14 RGS-box protein GAP activity towards $G\alpha_{i1}$(DM) protein when wildtype RGS14 sequence protein is used ("WT"; estimated $IC_{50}$ of 0.32 μM), and this inhibitory activity is largely lost when using a cysteine-free version of the recombinant RGS14 RGS-box protein ("-cys"), implicating cysteine modification as the predominant mechanism of (allosteric) inhibition. **(B)** Three-dimensional representation of the highest scoring docked pose of chaetocin within the Gα-interacting shallow canyon of the RGS14 RGS-box. The considerable distance between chaetocin's internal disulfide bonds (yellow) and Cys-130 of RGS14 (also yellow), located at the allosteric 'hot-spot' distal to the Gα-binding interface, suggests potential thiol-based covalent modification rather than canonical PPI groove engagement.
